# Divergence of TOR and AMPK regulators across 210 eukaryotes predicts a dispensable TOR in Toxoplasma gondii

**DOI:** 10.64898/2026.09.26.754375

**Authors:** Aditya Patwardhan, Euwen Brennan, Bruno Martorelli Di Genova

## Abstract

AMP-activated protein kinase (AMPK) and target of rapamycin (TOR) are the catalytic cores of the two major energy-sensing pathways in eukaryotes. Most of what is known about them is derived from model organisms. Previous comparative surveys have cataloged which AMPK and TOR pathway components are present in early-branching eukaryotes. However, they have not examined whether the present orthologs preserve the residues and interfaces mapped in model organisms. Here, we compared functionally characterized regions of 10 AMPK and TOR pathway orthologs across 210 eukaryotic species.

Our findings show that most core members of the AMPK and TOR pathways are broadly conserved among eukaryotes. The exceptions fall almost entirely within the medically significant phylum Apicomplexa. For example, the AMPKγ nucleotide-sensing subunit and the N-terminal HEAT-repeat region of TOR are significantly different in this phylum compared with their mammalian orthologs. Also, apicomplexan TOR orthologs have HEAT-repeat domains more than twice the human span, and this expansion accompanies repeated loss of the TOR-complex scaffolds RAPTOR and RICTOR.

These observations suggest that TOR signaling has changed in Apicomplexa, but sequence data alone cannot test this hypothesis. We deleted the sole TOR ortholog in Toxoplasma gondii (TgTOR). Three independent, sequence-validated TgTOR-deleted clones grew no differently from the wild-type control. These findings are important because TOR is a central regulator of eukaryotic growth, yet TgTOR-knockout parasites can still replicate. They also show that the presence of an ortholog alone does not demonstrate functional conservation.

**Significance statement:** Comparative studies of signal transduction pathways typically ask only whether an ortholog is present. We assessed both presence and sequence divergence in 10 core AMPK and TOR pathway members across 210 eukaryotes. The greatest departures were found in Apicomplexa: AMPKγ has expanded, divergent nucleotide-sensing regions; TOR has an expanded HEAT-repeat region; and the TOR-complex scaffolds RAPTOR and RICTOR are absent. Thus, apicomplexans retain TOR orthologs without the scaffolds needed to assemble TOR complexes. We deleted the sole TOR ortholog in *Toxoplasma gondii* (TgTOR). Parasites lacking TgTOR replicated at rates indistinguishable from wild type under standard culture conditions. Since TOR is a central regulator of eukaryotic growth, the ability of a parasite to proliferate without it shows that recognizable orthologs can persist even after their pathway has diverged.

## Introduction

Eukaryotic cells regulate their growth in response to nutritional and energy changes through sensing pathways. AMP-activated protein kinase (AMPK) promotes catabolism during low-energy states. In opposition to AMPK, target of rapamycin (TOR) promotes growth when nutrients are abundant (Hardie et al. 2012; Herzig and Shaw 2017; Saxton and Sabatini 2017; González et al. 2020).

AMPK is a heterotrimeric holoenzyme formed by a catalytic α subunit, a scaffolding β subunit, and a regulatory γ subunit (Xiao et al. 2007; Oakhill et al. 2010; Yan et al. 2018; Figure 1A). The γ subunit binds ATP, ADP, and AMP, and this nucleotide exchange couples the complex to the cellular energy state (Chen et al. 2012). In response to cellular changes, AMPK can be phosphorylated by different upstream kinases, including the liver kinase B1 (LKB1) in animals, and by SAK1, TOS3, and ELM1 in fungi, and by GRIK1 and GRIK2 in plants (Hawley et al. 2005; Woods et al. 2005; Hedbacker and Carlson 2008; Shackelford and Shaw 2009; Shen et al. 2009; Liu et al. 2011).

**Figure 1.**
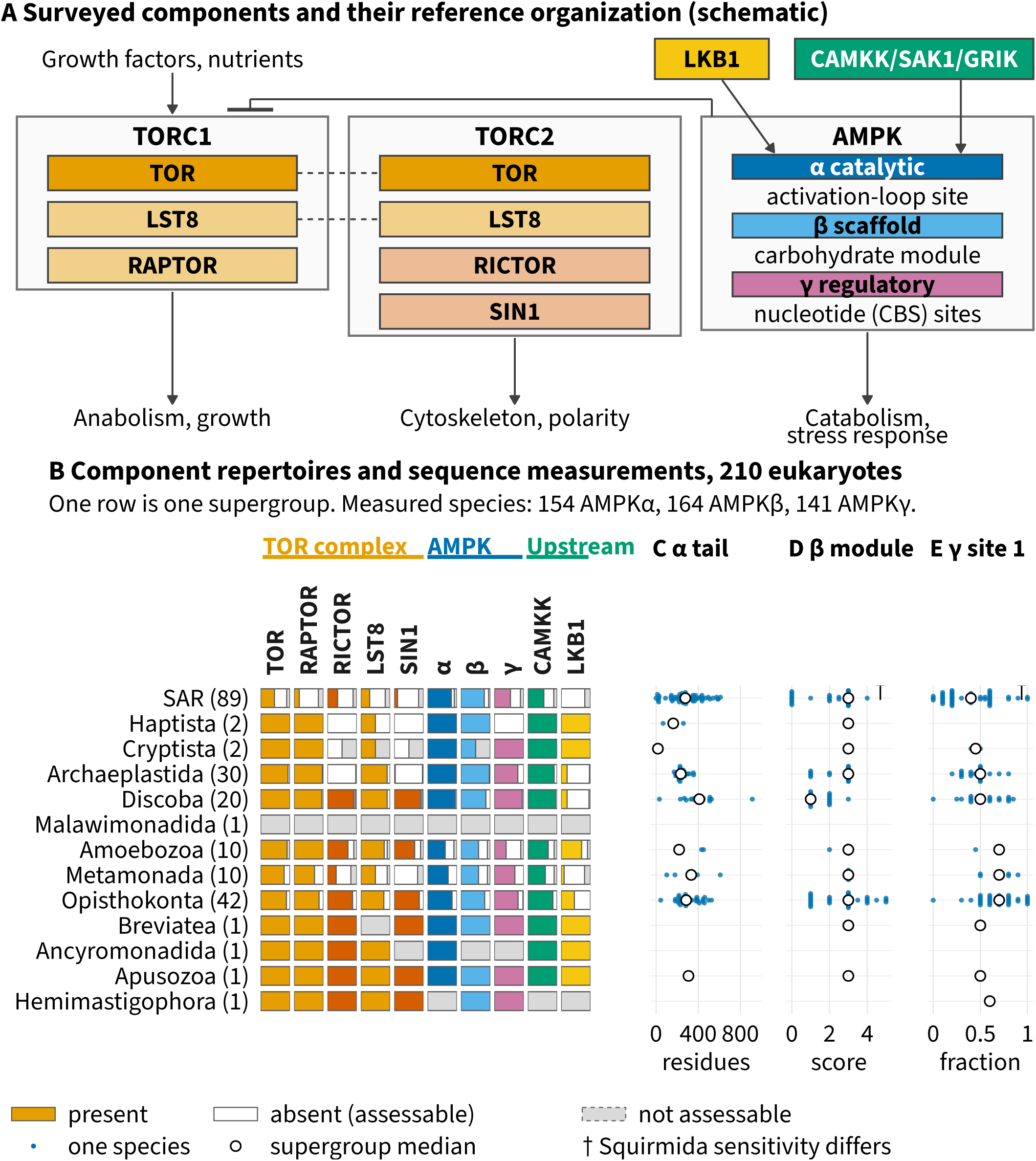
Surveyed TOR and AMPK regulators, their recovery across 210 eukaryotes, and three sequence measurements for the remaining subunits. (A) The surveyed regulators are shown in their respective complexes. Dashed links indicate that TOR and LST8 are shared, and each AMPK subunit is labeled with the region measured in later figures. The T-bar indicates that AMPK inhibits TOR complex 1. This panel is a reference diagram and includes no survey data; box colors identify the regulators in panel B. (B) The status of each of the 10 regulators in each of 13 supergroups across all 210 species. Supergroup sizes are shown in parentheses, and rows are ordered by the dated tree (Supplementary Figure S1B). Each cell indicates presence (regulator color), absence (white), or not assessable (gray); cell widths represent the fraction of sampled species in each supergroup. Apicomplexa, the focus of panels C–E and Figures 2–5, includes 41 sampled species. (C) AMPKα tail length, measured as the number of residues C-terminal to the anchor position. (D) AMPKβ module score, based on matches at five reference positions, on a scale from 0 to 5. (E) AMPKγ site 1 match fraction, shown as a species median and calculated using a different denominator from the per-copy count in Figure 2C. In panels C–E, each dot represents one species from a shared list of 34 phyla grouped under the supergroups in panel B. Crosses in the shaded NA strip mark species without a measurable value, open circles mark phylum medians, and n/N gives the number of measurable values among sampled species. The Squirmida sensitivity analysis and species with no phylum-ranked ancestor are listed in Supplementary Table S10; the same matrix for 37 clade blocks is shown in Supplementary Figure S1; and the species-by-target matrix is provided in Supplementary Table S7. Alt text: Panel A diagrams TOR complex 1 (TOR, LST8, RAPTOR), TOR complex 2 (TOR, LST8, RICTOR, SIN1), and the AMPK heterotrimer (α, β, γ), labeling regions measured in later figures. Dashed lines show TOR and LST8 shared between TOR complexes. Growth factors and nutrients feed into TOR complex 1; LKB1 and CAMKK/SAK1/GRIK feed into AMPKα. AMPK inhibits TOR complex 1, and arrows connect the complexes to anabolism and growth, cytoskeleton and polarity, or catabolism and stress response. Panel B shows 13 supergroups (SAR, 89 species, down to groups with one) across 10 regulators, grouped as TOR complex, AMPK, and upstream. Outlined bars indicate regulator presence (colored), absence (white), or not assessable (gray). Most regulators occur in most Opisthokonta and Archaeplastida; RICTOR and SIN1 are largely absent from Archaeplastida and Haptista. The lone Malawimonadida species is unassessable for all regulators. Panels C–E show three dot plots for 34 phyla, grouped under bold supergroup headings: SAR, Haptista, Archaeplastida, Discoba, Malawimonadida, Amoebozoa, Metamonada, Opisthokonta, and Hemimastigophora. They measure AMPKα tail length (residues), AMPKβ module score (0–5), and AMPKγ site 1 match fraction (0–1), respectively. Filled dots represent species; open circles, phylum medians. Shaded NA strips separated by dashed lines contain crosses for species without measurable values. Each panel’s n/N column reports measurable versus sampled species; for Apicomplexa, values are 23/41, 27/41, and 21/41, with daggers on β and γ.

TOR functions as a complex whose activity is regulated by its binding partners. TOR complex 1 (TORC1) requires RAPTOR and LST8 and promotes growth and protein synthesis. The other complex, TOR complex 2 (TORC2), is formed with RICTOR, SIN1, and LST8 and regulates a distinct metabolism, cytoskeleton, and other downstream cellular processes (Kim et al. 2002; Jacinto et al. 2006; van Dam et al. 2011; Saxton and Sabatini 2017).

Most of our current understanding of the AMPK and TOR pathways comes from model organisms, which show that both pathways are essential for cell survival (González et al. 2020). The Apicomplexan phylum is formed by early divergent single-celled eukaryotes of medical importance, including the causative agents of malaria, cryptosporidiosis, and toxoplasmosis (Arisue and Hashimoto 2015). As early branching eukaryotes, apicomplexans provide an opportunity to examine how conserved nutrient and energy sensing pathways diverged over time. *Toxoplasma gondii* is one of the most widespread human parasites (Ben-Harari and Connolly 2019). During its life cycle, it alternates between developmental stages that impose different metabolic demands (Messina et al. 2025; Rimple et al. 2025). Recent studies have shown that AMPK signaling is required for *T. gondii* metabolic adaptation. *T. gondii* AMPK (TgAMPK) activity is highly activated in extracellular parasites and returns to basal levels after invasion of the host cell. Also, knockdown of TgAMPK subunits suppresses parasite growth (Li et al. 2023). Whether TOR has any comparable role in these events has not been established.

Previous works have shown that many early-branching eukaryotes AMPK and TOR pathway core regulators (van Dam et al. 2011; Roustan et al. 2016; Johnson et al. 2025). While these studies reported which orthologs are present in these species, they did not analyze whether the present orthologs have diverged from their counterparts in well-characterized model systems. The distinction matters. For example, *Arabidopsis thaliana* SnRK1 is homologous to AMPK but differs drastically from mammalian AMPK in its regulatory behavior (Emanuelle et al. 2015). The same is the case for AMPK orthologs in *Saccharomyces* (Rubenstein et al. 2008; Mayer et al. 2011).

The domains of AMPK and TOR have been mapped in detail in model organisms (Scott et al. 2004; Xiao et al. 2007; Chen et al. 2012; Li et al. 2015; Hardie et al. 2016). Using these data, we have surveyed and analyzed the sequences of 10 AMPK and TOR pathway regulators across 210 eukaryotic species, with particular emphasis on early-diverging lineages. Each ortholog was assessed for conservation of the functional domains. First, we analyzed each species for the presence of orthologs of the 10 targets. The successfully identified sequences were scored for several functional and evolutionary analyses. Altogether, the data allowed us to ask two important biological questions: 1) did the discovered orthologs preserve the functional residues and domains mapped in model organisms, and 2) did the loss of an interacting ortholog coincide with increased sequence divergence.

## Results

### Retention of the ten TOR and AMPK regulators across 210 eukaryotes

We surveyed 210 predicted proteomes to determine which of the ten AMPK regulators (AMPKα, AMPKβ, AMPKγ, LKB1, CAMKK) and TOR regulators (TOR, LST8, RAPTOR, RICTOR, SIN1) are present across major eukaryotic lineages (Figure 1A; Supplementary Figure S1). For this search, we focused primarily on single-celled eukaryotes, using model organisms as controls (Supplementary Tables S1 and S2) and focusing primarily on the Apicomplexa phylum. For each target, orthologs were identified using two independent lines of evidence. We inferred orthogroups for all 10 targets using OrthoFinder (Emms and Kelly 2019) and SonicParanoid (Cosentino and Iwasaki 2019) in parallel. As a separate analysis, we used domain-based search using HMMER (Eddy 2011), CDD signatures with rpsblast (Wang et al. 2023), and InterProScan signatures (Jones et al. 2014; Blum et al. 2025) (Supplementary Tables S3 and S4). We aimed to be as thorough as possible, especially because the species cohort we chose is diverse and divergent. After obtaining the candidate pool for each target, we used phylogentic tree to assess orthology. For this, we ran phylogenetic analyses with positive controls (Orthologs from model organisms) and negative controls (sequences with similarity that are not orthologs) as previously done (van Dam et al. 2011) (Supplementary Table S5; Supplementary Figure S2). We classified each regulator as present, absent, or not assessable. A target was scored present when at least one candidate was identified using any of the methods described after the phylogenetic evaluation (Supplementary Figure S3A; Supplementary Table S6). Absence was scored only in genome-derived proteomes (Supplementary Table S7)

In the analyzed species cohort, we recorded ortholog absence for all 10 targets (Figure 1B and Supplementary Tables S1 and S2a). Our results are consistent with previous comparative surveys (van Dam et al. 2011; Roustan et al. 2016; Johnson et al. 2025;) (Supplementary Table S8).

In our searches, AMPKα was the most prevalent ortholog, whereas LKB1 was the least prevalent. Our results also show that TOR was present in only 74.7% of assessable species (136/182; Wilson 95% CI 68.0 to 80.5), while AMPKα was present in 86.8% (158/182; Wilson 95% CI 81.1 to 91.0) (Supplementary Table S1).

### Sequence measurements on the AMPK subunits by phylum

How closely the orthologs resemble their counterparts in model organisms has not been examined before. We assessed three sequence features implicated in AMPK function across all orthologs and compared them across phyla.

One of the first changes we measured was the C-Terminal domain length in AMPKα orthologs (Figure 1C). C-Terminal length spanned roughly a fourfold range across phyla, from a median of 130.5 residues in Chytridiomycota to 610 in Preaxostyla; among the six phyla with at least ten measurable species the range was narrower, from 231 residues in Streptophyta to 418 in Euglenozoa (Figure 1C and Supplementary Table S9; Supplementary Table S10). We also tested whether AMPKα tail length repeatedly evolved in lineages that lost the same pathway regulators. Two convergence analyses detected no signal, although the small number of independent losses limits the strength of this result (Supplementary Figure S4; Supplementary Table S11). We then assessed the conservation of the AMPKβ carbohydrate-binding module. Among the phyla with at least ten measurable species, the conservation was lowest in Ascomycota and Euglenozoa, both with a median module score of 1, and highest in Apicomplexa, Chlorophyta and Ciliophora, all at 3; across all phyla the highest medians were in Porifera, Chordata and Blastocladiomycota (Figure 1D; Supplementary Table S12; Supplementary Table S10).

Lastly, we investigated the conservation of the AMPKγ residues implicated in AMP binding. To do so, we aligned crystal structures of human AMPKγ1 and identified orthologs for all species. Using these alignments, we calculated how many amino acids that bind AMP in human AMPKγ1 are conserved (Supplementary Table S13 and Figure 1E; Supplementary Figure S5; Supplementary Table S14).

These patterns were independent of alignment strategy. Pairwise alignments to the human reference and a joint alignment of all orthologs differed for 79 of the 227 copies scorable by both procedures and 62 of 139 species. However, downstream model estimates were unchanged: for every site 1 coefficient, the 95% interval spanned zero under both procedures (Supplementary Figure S5; Supplementary Figure S6A; Supplementary Table S13).

Coccidia, a subclass of the Apicomplexa phylum, has AMPKγ orthologs that are more than twice the length of typical eukaryotic ones, with a median of 909 residues against 396 elsewhere (Figure 2A and E; Supplementary Table S15; Supplementary Figure S6C). Sarcocystids carried the longest orthologs, Eimeria intermediate ones, and Cryptosporidium the shortest, all of which overlap the upper range observed in other eukaryotes (Figure 2E; Supplementary Table S15).

**Figure 2.**
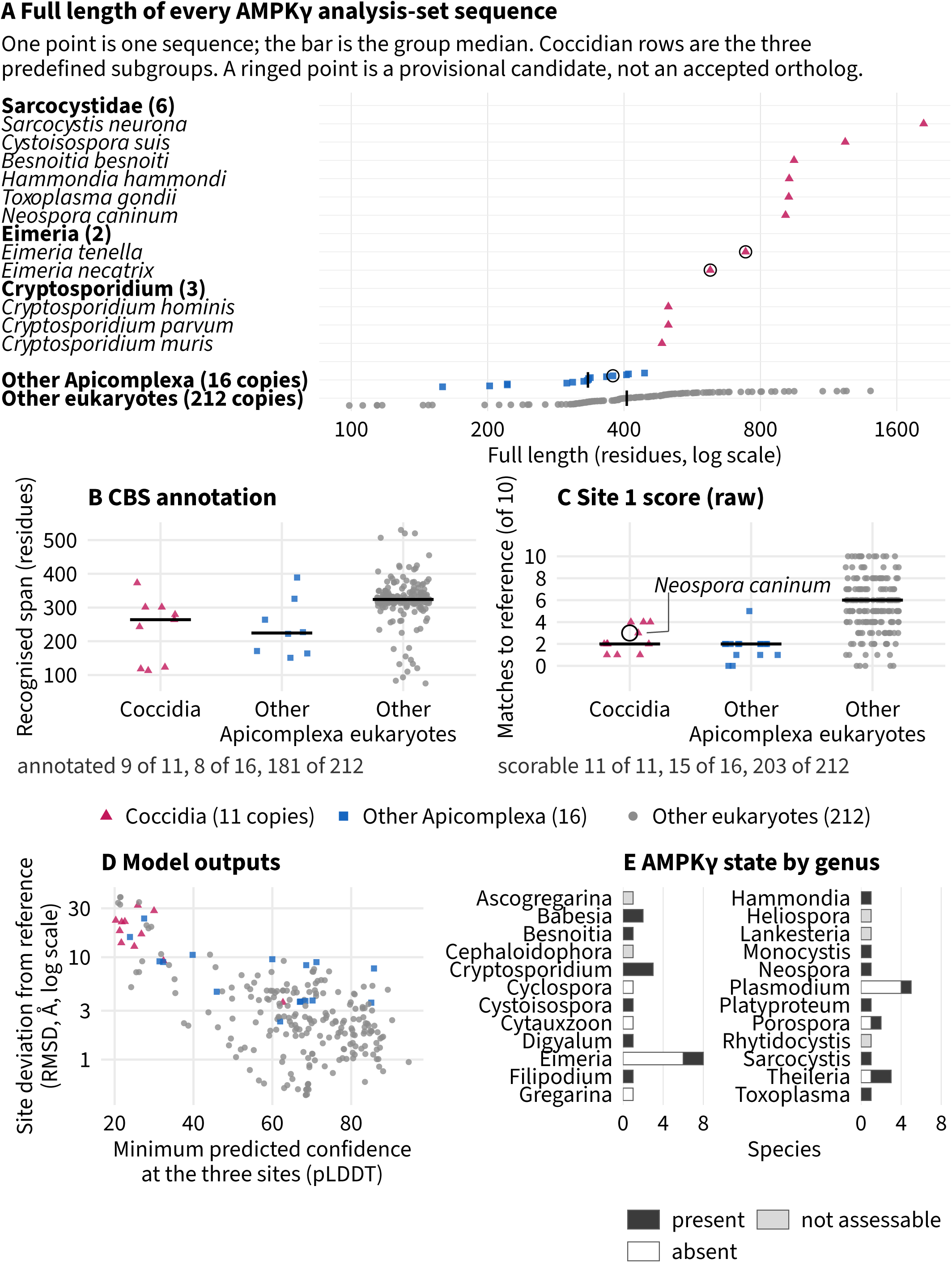
AMPKγ sequences in the focal coccidian cohort, compared primarily by length. The 239 AMPKγ analysis-set sequences from 142 species are grouped into 11 from the coccidian cohort, 16 from other sampled Apicomplexa, and 212 from other eukaryotes; color and shape encode these groups. Open symbols indicate the three provisional AMPKγ candidates. (A) Full length of each copy on a logarithmic axis, with the cohort resolved into individual sequences and grouped as Sarcocystidae, *Eimeria*, and *Cryptosporidium*. A bar marks the median for each comparison group: 333 residues in other sampled Apicomplexa and 405.5 in other eukaryotes. (B) Span of each copy recognized by annotation databases as cystathionine β-synthase (CBS) domains; the panel caption identifies the copies with annotations in each group. (C) Complete pairwise site 1 contact score: the number of the ten site 1 positions matching the human residue, plotted only when all ten positions are scorable—11 of 11, 15 of 16, and 203 of 212. The circled *Neospora caninum* copy has no secure residue correspondence between the two mapping procedures (Supplementary Table S15). (D) Median minimum predicted confidence across the three nucleotide sites versus median deviation of each site from the reference; both axes describe the model. (E) AMPKγ state of all 41 sampled apicomplexan species, grouped by genus and shown in the three states from panel 1B. Counts and medians are from Supplementary Table S15. Alt text: Panel A plots focal-copy lengths on a log scale. Headings group six Sarcocystidae, two *Eimeria,* and three *Cryptosporidium* copies, followed by individual species; bottom rows summarize other Apicomplexa and eukaryotes, with bars marking group medians. Sarcocystidae copies are longest (900–1,800 residues), followed by *Eimeria* (620 and 740) and *Cryptosporidium* (∼500). Comparison groups center near 330 and 405, overlapping the cohort’s lower range. Panels B–C show dot strips for three groups, median bars, and available counts. Panel D plots minimum predicted confidence against site deviation (log scale); coccidian copies have low confidence and high deviation. Panel E stacks genus-level counts as present, absent, or not assessable; *among eight Eimeria* species, six are absent and two present.

Intriguingly, several apicomplexans do not have an AMPKγ ortholog. AMPKγ was absent from six of the eight sampled *Eimeria* species and from *Cyclospora cayetanensis*, whereas the two recovered *Eimeria* sequences were detectable only through genome-based profile searches. AMPKγ is also absent in four of the five sampled *Plasmodium* species, the exception being *Plasmodium gallinaceum*, whereas all three sampled *Cryptosporidium* species retain a single short ortholog (Figure 2E; Supplementary Table S15).

We asked whether the additional residues in coccidian orthologs corresponded to extra nucleotide-binding domains. Interestingly, coccidian orthologs contained proportionally less recognizable cystathionine β-synthase (CBS) domain sequence than other eukaryotic orthologs (median annotation fraction 0.29 versus 0.77), suggesting that most additional residues are not nucleotide-binding domains (Supplementary Figure S6C). The contact residues themselves diverged furthest. Coccidian orthologs have a median of only two of the ten residues corresponding to the AMP-contact positions of human γ1 site 1 (Figure 2B-D; Supplementary Table S15; Supplementary Figure S5). Despite these changes, TgAMPKγ remains essential for *T. gondii* replication. Depleting this subunit prevents AMPKα phosphorylation and suppresses parasite replication (Li et al. 2023).

Our analysis also showed that divergence from the human sequence was concentrated at the CBS domains. Outside these domains, coccidians differ from the human sequence to a similar extent as other species, with median identities of 0.232 and 0.247 across the remaining CBS-domain positions. (Supplementary Table S15). Putative interactors are listed in Supplementary Table S13 and shown schematically in Supplementary Figure S6.

Divergence at this scale could reflect either loss of constraint or adaptation to an alternative function. We therefore compared evolutionary rates at these positions with those at burial-matched positions in the ortholog. Across 141 species, nucleotide-contact residues evolved faster than the protein-wide average (mean relative substitution rate 1.75) but significantly more slowly than 10,000 burial-matched control sets (2.34 and 2.40 under the two burial definitions; *P* = 0.036 and 0.016; Supplementary Table S16). These results suggest that residues have diverged considerably from the mammalian state while evolving more slowly than exposure-matched positions, that is, under relative rather than absolute constraint.

To infer the consequences of changes in the CBS domain, we replaced the ten CBS contact residues of human γ1 with the *Toxoplasma* sequence (Supplementary Figure S7). This modeling resulted in displacing the site by roughly half as much as alanine substitution. The nucleotide-binding pocket of yeast Snf4, a γ ortholog that is not allosterically activated by AMP (Mayer et al. 2011), clustered with the altered constructs rather than with intact mammalian orthologs at CBS1 (Supplementary Figure S7). Together, these results suggest that the nucleotide-sensing surface of apicomplexan AMPKγ has diverged while evolving more slowly than exposure-matched positions. This change is compatible with an altered nucleotide-sensing function, which remains to be tested biochemically.

### Length and repeat density of the TOR N-terminal HEAT region

Across TOR orthologs, our results show that divergence is concentrated in the N-terminal HEAT-repeat region. In coccidians, this region has more than doubled in length while the catalytic FAT-kinase-FATC module remains comparatively conserved. For example, *T. gondii* carries a 2,944-residue HEAT region, compared with 1,330 residues in human mTOR, whereas the catalytic FAT-kinase-FATC module is expanded only 1.63-fold, from 1,037 to 1,693 residues (Figure 3A; Supplementary Tables S17 and S18). A similar expansion evolved independently in the brown alga *Ectocarpus siliculosus*, which carries a 2,846-residue HEAT region despite being distantly related to apicomplexans (Supplementary Table S17).

**Figure 3.**
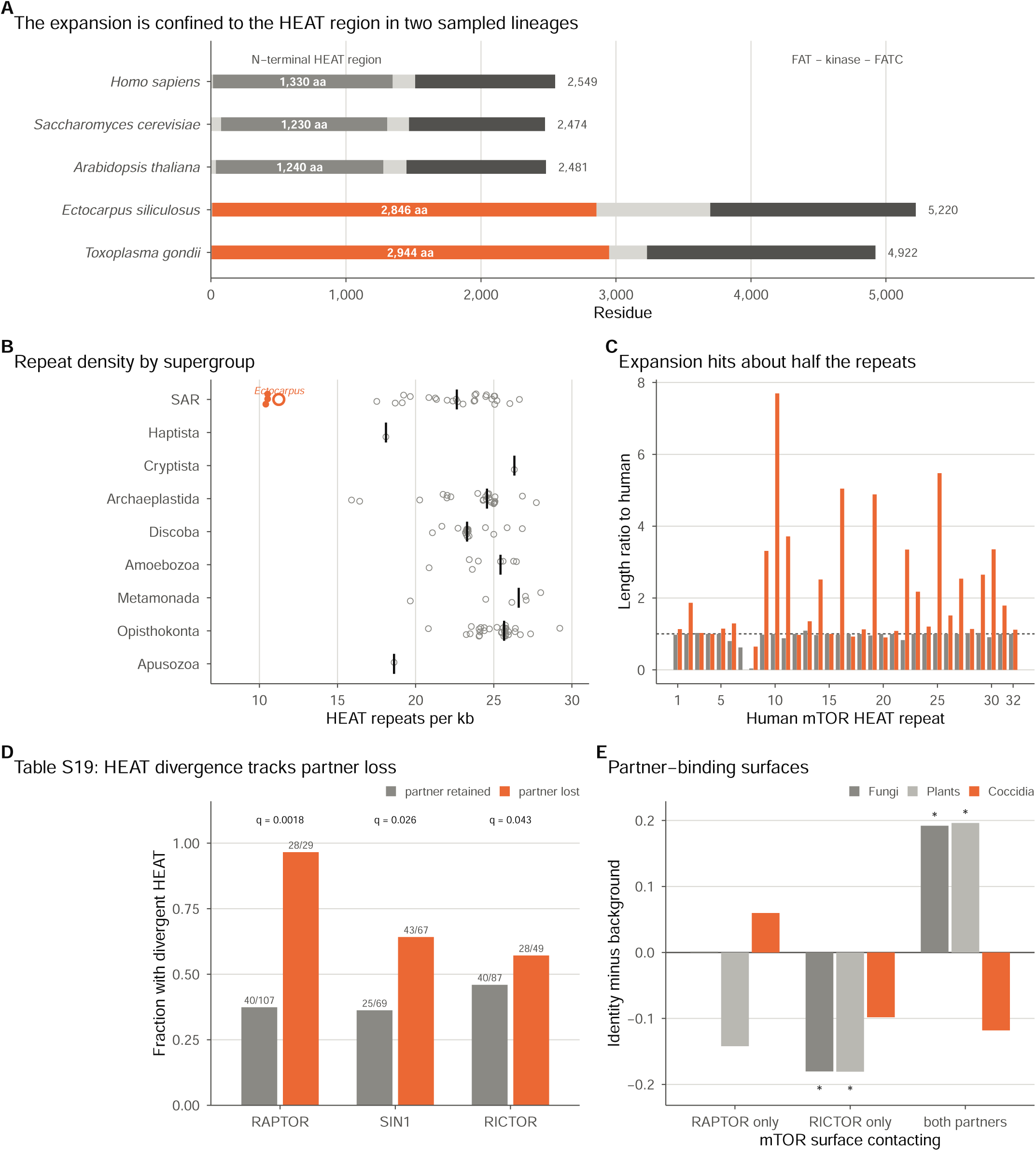
TOR N-terminal HEAT architecture across eukaryotes and its relationship to TORC partner loss. One copy per species was included (the longest usable copy for the tree); presence-only copies were excluded. Colors distinguish Apicomplexa from other eukaryotes. (A) Scaled domain architectures for five species, with the mapped HEAT span shown in color and the C-terminal FAT-kinase-FATC module shown in dark. (B) HEAT repeat density (repeats/kb): points represent individual species, and bars show supergroup medians for 127 species whose mapped regions cover at least half of the human reference array (Supplementary Table S17). (C) Median mapped length of each human reference repeat relative to its length in humans, comparing three expanded coccidian copies with all others. Repeat 7 is unmapped in coccidian copies; positions transferred by alignment are less robust than span totals (Supplementary Table S18). (D) Fraction of 136 genome-grade species with divergent-coded HEAT regions, grouped by retention of each TORC partner (Supplementary Table S19). Counts appear above bars; q values are Holm-adjusted (Supplementary Table S19). (E) Human mTOR identity at partner-binding surfaces minus identity at non-interface HEAT positions in fungi, plants, and coccidia, across RAPTOR-specific, RICTOR-specific, and shared surfaces. Asterisks indicate permutation *P* < 0.05; the coccidian comparison includes five copies (Supplementary Table S20). Alt text: Five panels. Panel A shows one TOR ortholog per species; *Homo sapiens*, *Saccharomyces cerevisiae*, *Arabidopsis thaliana*, *Ectocarpus siliculosus,* and *Toxoplasma gondii*; with a scale bar divided into an N-terminal HEAT region and a dark FAT-kinase-FATC module (0–∼5,200 residues). The orange HEAT region spans 1,230–1,330 aa in the first three species, whose orthologs end at 2,474–2,549 aa; in *E. siliculosus* and *T. gondii*, it spans 2,846–2,944 aa, and orthologs end at 5,220 and 4,922 aa. Panel B plots HEAT repeats per kb across nine supergroups, with supergroup medians marked; most species range from ∼15–30, while orange coccidian points and *E. siliculos*us are near 10–11 in SAR. Panel C shows median mapped lengths of 32 human mTOR HEAT repeats relative to human length (dotted line = 1). Orange coccidian bars reach ∼2–8 for about half the repeats; gray bars for other eukaryotes remain near 1. Panel D compares the fraction of species with divergent HEAT regions by partner status (present, gray; lost, orange): RAPTOR, 40/107 vs. 28/29 (q = 0.0018); SIN1, 25/69 vs. 43/67 (q = 0.026); RICTOR, 40/87 vs. 28/49 (q = 0.043). Panel E plots identity minus background (−0.2 to 0.2) at RAPTOR-only, RICTOR-only and shared-partner surfaces in fungi, plants and coccidia. Fungi and plants decrease ∼0.18 at the RICTOR-only surface and increase ∼0.19 at the shared surface (asterisks); coccidia decrease at the shared surface.

We measured the number of HEAT repeats per unit length. All three coccidian TOR orthologs contained fewer HEAT repeats per unit length than any other sampled eukaryotic TOR, indicating that a similar repeat architecture spans a much larger sequence length (Figure 3B; Supplementary Table S17). Approximately half of the mapped HEAT repeats were more than 1.5 times the length of their human counterparts, whereas most repeats in other eukaryotes remained close to the human state (Figure 3C; Supplementary Table S18).

Additionally, species lacking a RAPTOR ortholog almost invariably have a divergent HEAT region (Supplementary Table S19). Of the 29 RAPTOR-absent species, 28 fell below the median HEAT-region identity relative to the human reference, compared with only 40 of 107 species that retain RAPTOR (Figure 3D; Supplementary Table S19). More broadly, divergence of the HEAT region and loss of RAPTOR, SIN1, and RICTOR occur on the same branches of the tree more often than expected by chance (Holm-adjusted P = 0.0018, 0.026, and 0.043) (Supplementary Table S19). We did not detect changes when RICTOR is present, indicating that loss of a single binding partner does not relax sequence constraint at its interaction interface (Figure 3E; Supplementary Table S20).

### Evidence for Coordinated Loss Within, but Not Between, AMPK and TOR Modules

Given the differences we observed in the AMPK heterotrimer across the 210 species we analyze, we assessed the conservation of the AMPKα activation site. Our results show that, despite changes in the AMPKα C-terminus, the activation site is significantly conserved across the analyzed species (Figure 4A-C and Supplementary Table S21). Every sampled species with an AMPKα ortholog had at least one AMPKα copy conserving the activating threonine residue, and no species carried only non-threonine variants.

**Figure 4.**
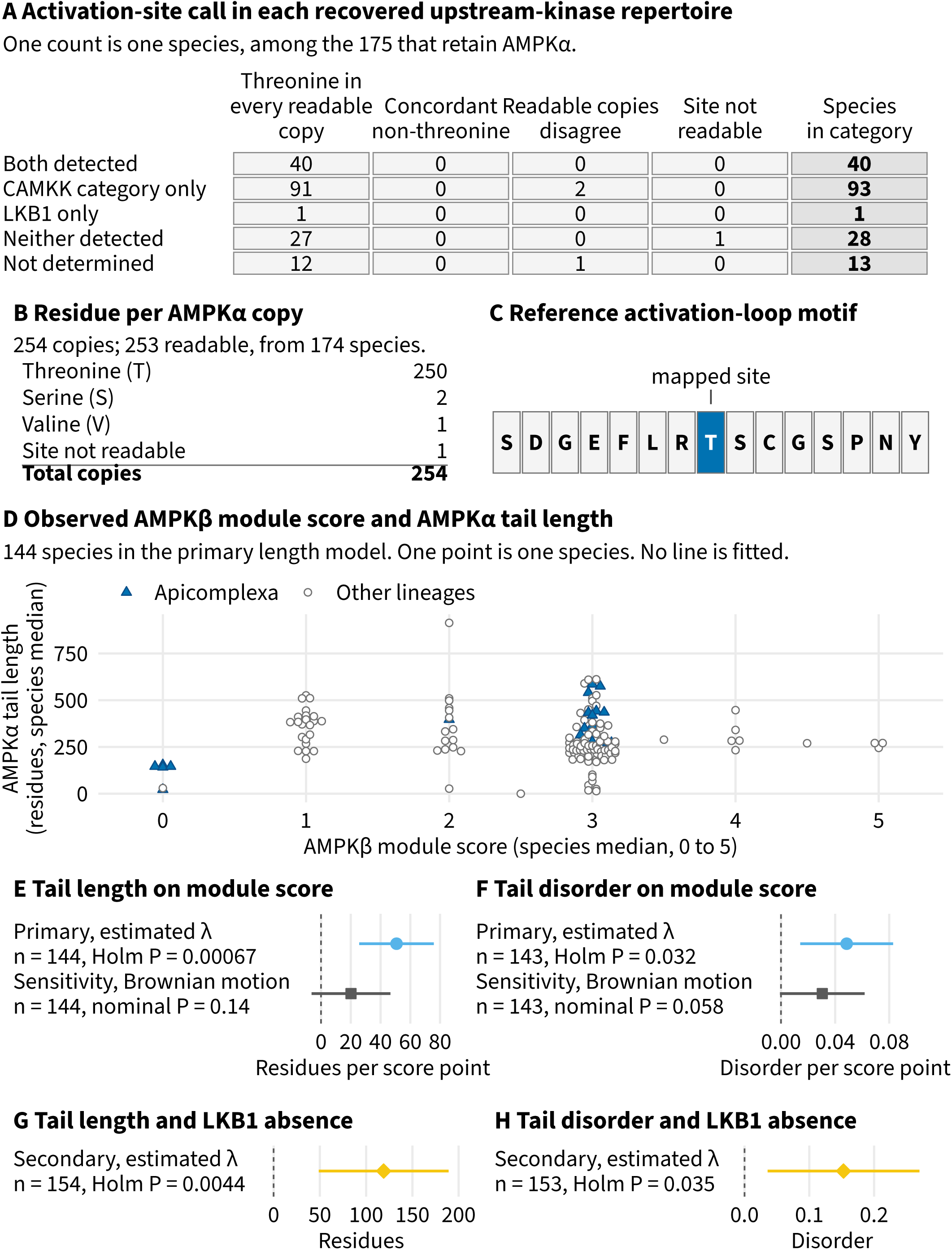
AMPKα activation site, recovered upstream kinases, and AMPKβ module versus AMPKα tail. (A) Activation-site calls across 175 species retaining AMPKα, cross-tabulated by recovered upstream-kinase repertoire. Rows show five kinase categories; columns indicate threonine in all readable copies, concordant non-threonine, disagreement among readable copies, or an unreadable site. (B) Residue at the mapped activation site for each AMPKα copy, counted separately from the species-level calls in A. (C) Reference activation-loop motif, with the mapped site marked. Per-copy calls and kinase data are shown in Supplementary Figure S9. (D) Species medians for AMPKβ module score and AMPKα tail length in the 144-species primary length model; color and shape indicate Apicomplexa. Points are offset along the score axis to separate ties, while lengths remain unchanged; no line is fitted. (E, F) Primary coefficients for tail length and disorder on module score under estimated phylogenetic signal, alongside full-sample Brownian-motion sensitivity estimates in the same units. Primary estimates are 50.7 residues and 0.049 disorder per score point. Holm-adjusted *P* values apply to the primary six-test family; Brownian-motion *P* values are nominal. (G, H) Secondary coefficients for the same responses by assessable LKB1 absence, from a separate four-test family. Bars show nominal 95 percent Wald intervals, and the dashed line marks zero. Each panel reports its sample size (Supplementary Tables S22 and S23). Alt text: Panel A is a five-row, five-column table of species counts. The rows are both detected, CAMKK category only, LKB1 only, neither detected, and not determined. The columns are threonine in every readable copy, concordant non-threonine, readable copies disagree, site not readable, and a bold species-in-category total. The counts are 40, 0, 0, 0, 40; 91, 0, 2, 0, 93; 1, 0, 0, 0, 1; 27, 0, 0, 1, 28; and 12, 0, 1, 0, 13. Panel B is a short count list: threonine, 250; serine, 2; valine, 1; site not readable, 1; total, 254 copies. Panel C is a strip of fifteen boxed letters (S D G E F L R T S C G S P N Y) with the eighth box, threonine, highlighted and labeled as the mapped site. Panel D plots AMPKα tail length in residues against the AMPKβ module score, from 0 to 5, for 144 species. Apicomplexan species are shown as filled blue triangles, and other lineages as open gray circles; points at each score are spread sideways into a symmetric cloud. Panels E and F each show two horizontal estimates with 95 percent intervals against a dashed zero line: the primary estimate under an estimated phylogenetic signal, whose interval lies entirely to the right of zero, and the Brownian-motion sensitivity below it, whose interval crosses zero. Panels G and H, beneath a rule and heading identifying them as the secondary four-test family, each show one estimate with an interval to the right of zero.

Given this result, we assessed the conservation of the AMPKα-activating kinases, LKB1 and the CAMKK family (Supplementary Figure S2). In contrast to conservation of the AMPKα activation site, the upstream regulatory repertoire is broadly divergent, and the human activating kinases, especially LKB1, are absent in several of the analyzed species (Figure 1B). When comparing the presence of AMPK and TOR regulators, we showed that species that have lost TOR were also more likely to lack detectable AMPK activators (adjusted *P* = 0.018; Supplementary Figure S8; Supplementary Table S22).

Surprisingly, even in species with no detectable AMPK upstream kinases, the AMPKα activation site remained conserved (Figure 4A-C; Supplementary Figure S9). This result highlights a limitation of ortholog analyses, as the absence of these activators does not mean the absence of AMPK regulation in these organisms. For example, TgAMPKα is required for *T. gondii* proliferation and its phosphorylation status changes during the parasite life cycle (Li et al. 2023; Yang et al. 2024). However, our analysis did not identify reliable AMPKα activators.

When comparing AMPKα C-terminal tail expansion, our analysis found that it correlated with the sequence properties of other AMPK members, rather than their presence or absence (Supplementary Table S22). Species with AMPKβ carbohydrate-binding domains more similar to the human domain tended to have longer, more disordered AMPKα tails (Figure 4D-F; Supplementary Figure S10B; Supplementary Tables S22 and S23). Species lacking LKB1 also had substantially longer, more disordered tails than those retaining the kinase (Figure 4G and H; Supplementary Table S22). After correction for multiple testing, tail length and disorder showed no consistent association with the presence or absence of TOR, RAPTOR, RICTOR, AMPKβ, or AMPKγ (Supplementary Tables S22 and S9; Supplementary Figure S10G).

Neither the AMPKγ nucleotide-contact score nor conservation of the AMPKβ carbohydrate-binding module showed consistent associations with loss of pathway regulators or disruption of the canonical complexes (Supplementary Tables S22 and S12; Supplementary Figure S10G).

Species lacking a TOR ortholog were also more likely to lack AMPK subunits (Supplementary Table S24). In addition, AMPKγ was absent in roughly two-thirds of species without TOR, compared with only one-sixth of species with TOR (Supplementary Table S24). The same pattern was observed for AMPKα, AMPKβ, RAPTOR, and RICTOR (Figure 5A; Supplementary Table S24).

**Figure 5.**
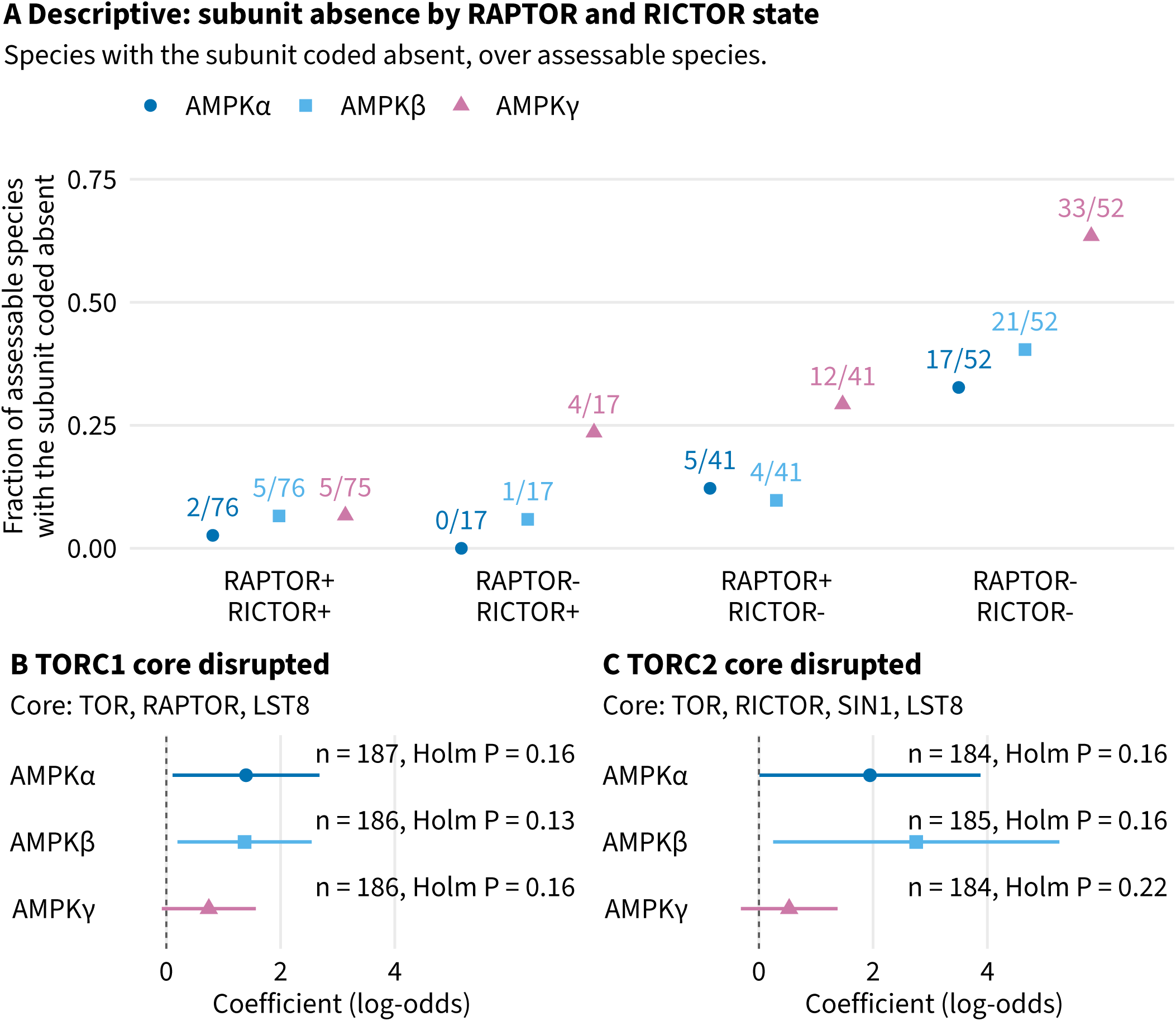
AMPK subunit absence by TOR-complex state: descriptive and model-based. (A) Fraction of assessable species lacking each AMPK subunit across the four observed RAPTOR/RICTOR combinations (+, present; −, assessable absence). Values above points show the number of absences and species assessed, with a separate denominator for each subunit. These are tip-level categories, not an evolutionary sequence; panels B and C use defined core complements rather than observed states. (B, C) Six prespecified phylogenetic logistic coefficients estimate AMPK subunit absence in relation to disruption of the TORC1 or TORC2 core (Materials and Methods); disruption means an assessable absence of at least one core regulator (Supplementary Figure S16). Rows show sample sizes and Holm-adjusted *P* values. Both panels use the same scale; bars indicate nominal 95 percent Wald intervals, and the dashed line marks zero (Supplementary Tables S22 and S24). Alt text: Panel A plots the fraction of assessable species lacking AMPKα, AMPKβ, or AMPKγ across four RAPTOR and RICTOR state categories. Each point is labeled with its count: 2 of 76, 5 of 76, and 5 of 75 when both scaffolds are present, increasing to 17 of 52, 21 of 52, and 33 of 52 when both are absent. Panels B and C are aligned forest plots on a shared log-odds scale, each with three rows and a dashed line at zero. In B, coefficients range from about 0.7 to 1.4, sample sizes from 186 to 187, and Holm P-values from 0.13 to 0.16. In C, coefficients range from about 0.5 to 2.8, sample sizes from 184 to 185, and Holm P values from 0.16 to 0.22. Four of the six intervals exclude zero, but none passes the family correction.

Most associations reflected coordinated loss within individual complexes rather than between the two pathways. Of the 90 pairwise comparisons, seven remained significant after correction, spanning five distinct regulator pairs (Supplementary Figure S11; Supplementary Table S25). Four pairs linked orthologs within the same complex: AMPKα with AMPKβ, AMPKα with AMPKγ, AMPKβ with AMPKγ, and RICTOR with SIN1. The only significant cross-pathway association was between LST8 and AMPKβ. Results were similar with more conservative coding of transcriptome-derived proteomes (Supplementary Table S25).

Across all 28 pruned trees examined, three regulator pairs showed correlated loss: AMPKα and AMPKβ, RAPTOR and LST8, and RICTOR and SIN1. No association between an AMPK regulator and a TOR-complex regulator was consistently recovered (Supplementary Table S26). Species that lost AMPKα often also lacked detectable AMPK activators (Supplementary Tables S27 and S25). However, ancestral-state reconstruction identified multiple branches where the kinase repertoire and AMPKα followed different evolutionary histories, indicating that their losses were not strictly coupled (Figure 6; Supplementary Tables S28 and S29; Supplementary Figure S12B).

**Figure 6.**
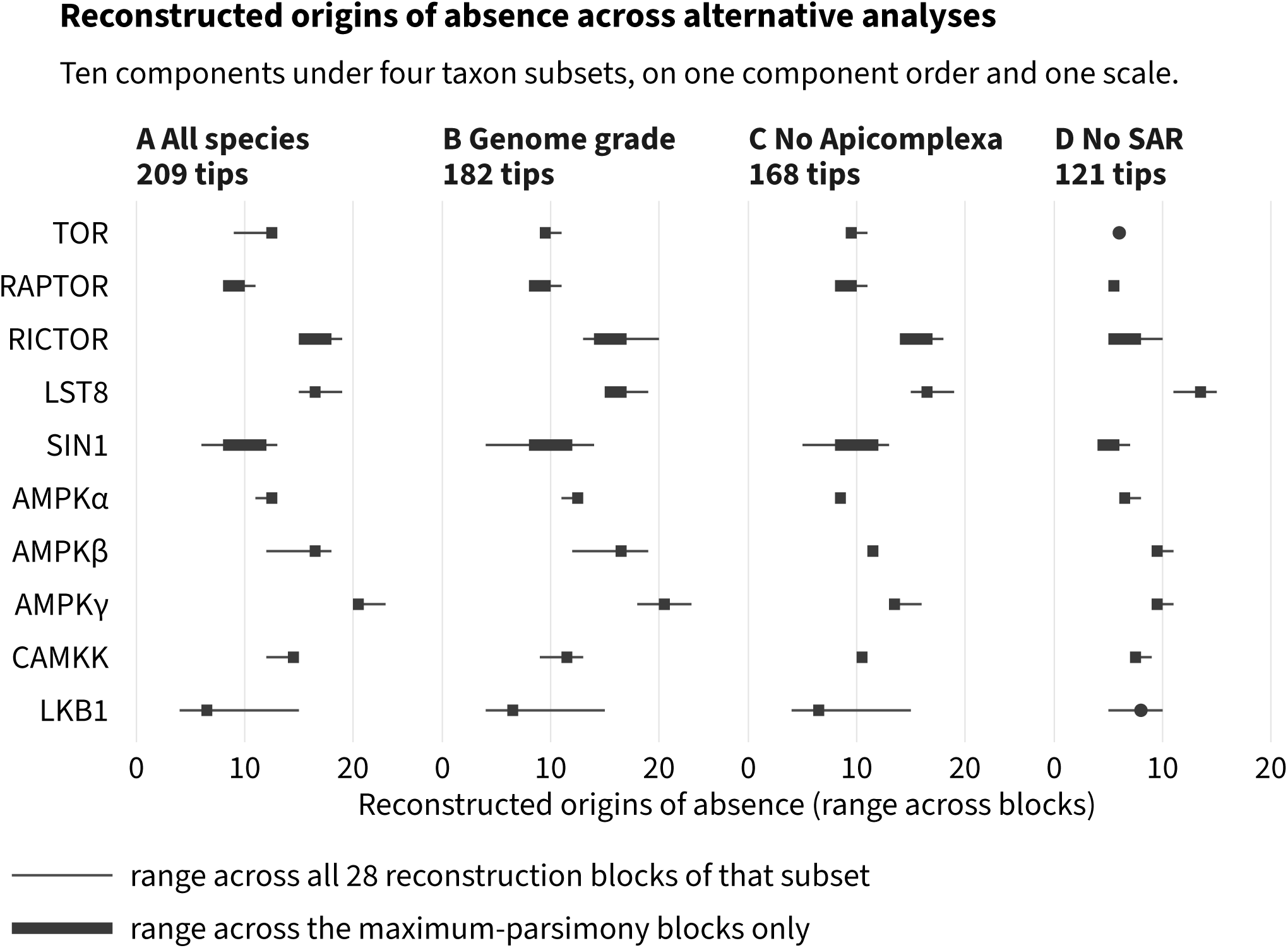
Reconstructed number of origins of absence across analyses. For each of ten regulators, ranges are shown across four taxon subsets using a shared regulator order and horizontal scale: (A) all 209 tips, (B) 182 genome-grade tips, (C) 168 tips excluding Apicomplexa, and (D) 121 tips excluding SAR. Each subset includes 28 reconstruction blocks, each combining topology, rooting, and substitution model. Thin lines span all blocks; thick bars span maximum-parsimony blocks only. Coincident endpoints are shown as a single point. Ranges reflect alternative analyses, not confidence intervals, and are not summed. State 1 denotes absence (Supplementary Figure S12A; Supplementary Table S28). Alt text: Four aligned panels share a list of ten regulators—TOR, RAPTOR, RICTOR, LST8, SIN1, AMPKα, AMPKβ, AMPKγ, CAMKK, and LKB1—in that order, along with a horizontal axis showing reconstructed origins of absence from 0 to about 24. The panels are labeled A to D and include all species (209 tips), genome grade (182), no Apicomplexa (168), and no SAR (121). Each row has a thin line showing the range across all 28 reconstruction blocks and a thick bar showing the maximum-parsimony blocks; some rows collapse to a point. In the all-species panel, AMPKγ lies farthest to the right, from 20 to 23, while LKB1 spans the widest range, from 4 to 15. Ranges generally shift left as tips are removed.

Since several of the species we analyzed are protozoan parasites, the changes shown here could be explained by a parasitic lifestyle. Before phylogenetic correction, obligate and intracellular species more frequently lacked AMPKγ and had simplified TOR complexes (Supplementary Table S30). Most associations weakened after accounting for shared ancestry, and nearly all confidence intervals overlapped zero (Supplementary Figure S13; Supplementary Table S30). These data cannot distinguish lifestyle from phylogenetic history.

### Deletion of the sole TOR ortholog of Toxoplasma gondii

Our comparative analyses identified Apicomplexa as a phylum in which TOR signaling differs most from the opisthokont organization. Although a minority of apicomplexans retain a TOR ortholog, eight of the 41 sampled species (Figure 1B), none retains RAPTOR and only one retains RICTOR, so the TORC1 and TORC2 scaffold orthologs are effectively absent from the phylum (Supplementary Table S2). Coccidians also have a highly divergent TOR HEAT-repeat region, whose expansion coincides with the loss of these interaction partners (Figure 3). The retention of TOR despite substantial sequence divergence suggested two possibilities: 1) TgTOR had been functionally rewired but remained essential, as indicated by the TgAMPK results (Li et al. 2023; Yang et al. 2024), or 2) it had lost its ancestral role in growth control. Since sequence data alone could not distinguish between these possibilities, we tested the latter hypothesis by deleting the sole TOR ortholog in *Toxoplasma gondii,* TGME49_316430.

A previous pooled genome-wide CRISPR screen (Sidik et al. 2016) detected no phenotype loss with TGME49_316430 gRNAs. Because pooled screens measure guide-directed disruption rather than the effect of a defined deletion, we deleted the locus directly. Using CRISPR, we deleted the *T. gondii* TOR ortholog (TgTOR) and recovered three independent knockout clones, each validated by PCR and sequencing across both the 5′ and 3′ junctions (Figure 7A and B; Materials and Methods). We generated multiple independent lines to reduce the chance that any resulting phenotype was caused by an unrelated mutation acquired during strain construction. Clones 15 and 17 showed no detectable TgTOR amplification, even with 1 µg of cDNA input per reaction; clone 7 showed a weak signal at that input. Relative to wild type, TgTOR transcript abundance was reduced 399-fold in clone 7 and at least 1,687-fold and 671-fold in clones 15 and 17, respectively (Figure 7C).

**Figure 7.**
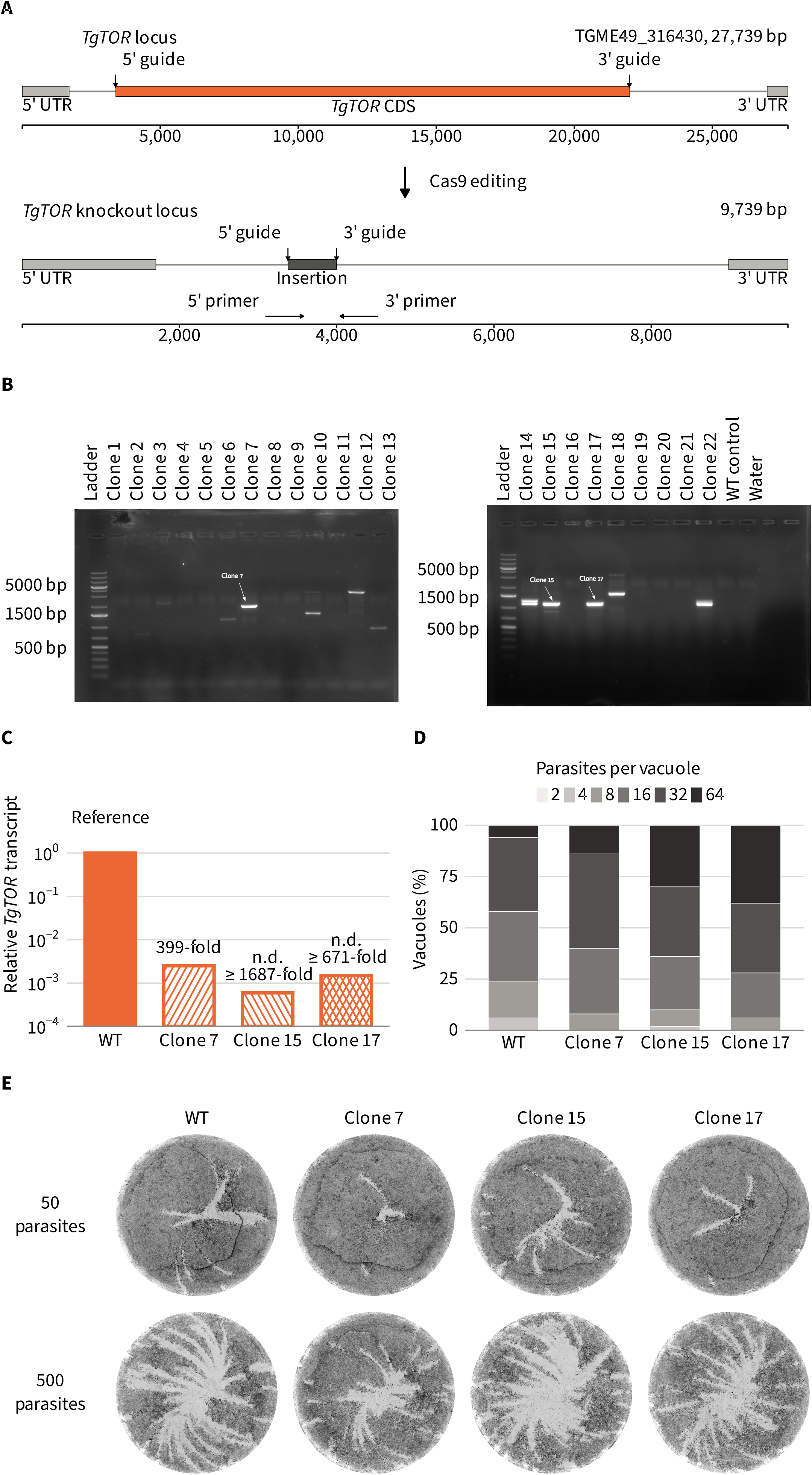
Development and characterization of *T. gondii* TOR-knockout strains. (A) TgTOR locus (TGME49_316430; 27,739 bp), with 5′ and 3′ guide sites flanking the coding sequence, and the 9,739-bp Cas9-edited locus containing an insertion between the cut sites. Arrows indicate primers used to amplify the edited region. (B) Edited-region amplification from 22 clones across two gels, with size ladders, a wild-type control, and a no-template water lane. Arrows mark clones 7, 15, and 17, which were validated by sequencing. (C) TgTOR transcript levels measured by qPCR relative to wild type, normalized to TgSAG1 and displayed on a log scale. Clone 7 shows a 399-fold reduction; transcripts were undetectable in clones 15 and 17, corresponding to reductions of at least 1,687-fold and 671-fold, respectively. Hatched bars indicate the detection floor for each reaction. (D) Parasite counts per vacuole after two days, shown as percentages by size class; 50 vacuoles per strain across three replicates. (E) Representative wells from a seven-day plaque assay with 50 or 500 parasites per well. Alt text: Five panels show TgTOR deletion and strain growth. A: Gene maps depict the 27,739-bp TgTOR locus, with gray 5′/3′ UTRs flanking an orange coding sequence and guide sites at both ends (scale, 25,000). Cas9 editing yields a 9,739-bp knockout, with a short dark insertion replacing the coding region between the guide sites. Facing 5′/3′ sequencing primers are shown below (scale, 8,000). B: Agarose gels show clones 1–13 and 14–22, alongside wild type and a water control; ladders mark 5,000, 1,500, and 500 bp. Arrows identify sequencing-validated clones 7, 15, and 17, each showing a bright band near 1,500 bp. C: Relative TgTOR transcript levels (log scale, 10 to 1): wild type is 1, and clone 7 is ∼2.5 × 10 ³ (399-fold lower). Transcript levels in clones 15 and 17 are undetected (at least 1,687- and 671-fold lower, respectively). D: Stacked bars show vacuole percentages for wild type and clones 7, 15, and 17, grouped by parasite count: 2, 4, 8, 16, 32, or 64. E: Eight plaque-stained wells show four strains at inocula of 50 or 500 parasites. Pale plaques on dark monolayers appear similar across strains at each dose.

We measured potential effects of TgTOR on parasite replication. For that, we infected human fibroblasts for two days and calculated the number of parasites per vacuole as a growth readout. The number of parasites per vacuole showed that all three knockout clones were indistinguishable from wild type (Figure 7D). To confirm these findings, we did a long-term growth assay. Seven-day plaque assays, performed with either 50 or 500 parasites per well, also showed no difference between wild type and any TgTOR-null clone (Figure 7E). Since plaque area reflects repeated rounds of invasion, replication, and egress, the seven-day assay can detect per-cycle replication differences that a two-day vacuole count cannot. Under the conditions tested, TgTOR is not essential for completion of the *T. gondii* lytic cycle.

## Discussion

AMPK and TOR core pathway members show variable distributions across eukaryotes (van Dam et al. 2011; Roustan et al. 2016; Johnson et al. 2025). Our calls agree with these surveys for 350 of 360 compared cells (van Dam et al. 2011), 620 of 675 (Roustan et al. 2016), and 714 of 820 (Johnson et al. 2025; Supplementary Table S8a). Agreement across three surveys based on different datasets and detection strategies supports the overall classification framework. The framework also resolves difficult candidates that profile-based approaches could not confidently classify (Johnson et al. 2025) (Supplementary Figure S3B). The clearest example is SIN1, which we recover in 18 species scored as absent or left unresolved by Johnson et al. (2025) (Supplementary Table S8b). Also, only 11 of the 1,255 present cells rely entirely on orthologs placed differently by the maximum-likelihood tree and the site-heterogeneous model: three in TOR, six in LST8, and one each in AMPKα and AMPKβ. These cells remain scored as present because the site-heterogeneous model served as a cross-check rather than a classifier (Supplementary Table S31; Supplementary Figure S3E).

Absence calls depend on genome annotation quality, even when genome-derived proteomes are prioritized (Supplementary Table S32). Transcriptome-derived proteomes never receive absence calls, so nine transcriptome-grade apicomplexans (eight gregarines and *Rhytidocystis pertsovi*, plus four of the six colpodellids) have only present or not-assessable states. Consequently, each apicomplexan loss node is polarized using just two genome-grade outgroup genera: *Chromera* and *Vitrella*. Broader sampling would not help, because all genome-grade apicomplexans outside Gregarinasina lie crownward of the losses, while the more distant outgroups, *Perkinsus* and dinoflagellates, already have genome annotations. Additional taxa would repeat the derived state rather than reveal when it originated. Cultivation is the limiting factor: deep-branching apicomplexans that bracket these nodes cannot be cultured and are known only from cells hand-picked from field-collected invertebrate hosts. As a result, Marosporida, sister to Coccidia and Hematozoa, is represented by a single transcriptome (Gile and Slamovits 2014; Keeling et al. 2014; Janouškovec et al. 2015; Janouškovec et al. 2019; Mathur et al. 2019; Mathur et al. 2021; Boisard et al. 2022; Richter et al. 2022).

The same constraint limits what can be affirmed about irreversibility. A regain, the reverse transition from absence to presence, appears as a present state nested within a clade of absences. A missed ortholog appears the same way. Missed orthologs are documented here: roughly a third of the LST8 and AMPKγ absences, and every AMPKα absence, are candidates the gene tree rejected rather than searches that found nothing. A regain would also require horizontal transfer of a multidomain kinase or scaffold into an obligate intracellular parasite, a route not documented for signaling components in these lineages. Where the reversible model wins, patchy absence calls are the simpler explanation; where the no-regain model wins, it wins on parameter penalty, not fit (Supplementary Table S33). LKB1 is the exception, with nothing detected in 141 of its 142 absences and no assessable apicomplexan retaining it. These comparisons therefore sort the targets by how clean their absence calls are, not by whether loss is permanent. That limit leaves the biological contrast intact: LKB1 is absent coherently across the Apicomplexa phylum, whereas the TOR partners are absent in a scattered pattern that repeated independent loss would equally produce.

Across 210 eukaryotic species, many orthologs differ substantially from their mammalian counterparts in domains involved in signaling and regulation. Apicomplexa offer the clearest examples: coccidian AMPKγ retains only two of the ten residues that contact AMP in human γ1 (Figure 2), while TOR is present but RAPTOR and RICTOR are absent (Figure 3). AMPKγ shows the greatest divergence. Earlier work identified an unusually large γ subunit in *T. gondii* (Li et al. 2023), and our survey shows that this architecture is shared across coccidians (Figure 2). This expansion coincides with divergence at the nucleotide-contact residues that sense AMP in mammals, even as the rest of the CBS region retains its identity. These residues still evolve more slowly than burial-matched controls, though faster than the protein-wide average. Together, these findings indicate that the nucleotide-sensing interface has been rewired rather than abandoned.

Similar departures from the mammalian AMPK mechanism are known elsewhere in eukaryotes. The γ-type subunit of plant SnRK1 differs from mammalian AMPK in its regulation by nucleotides (Emanuelle et al. 2015), while yeast Snf4 is not allosterically activated by AMP and instead responds to ADP (Mayer et al. 2011). While this still needs to be tested in Apicomplexa, these examples demonstrate that substantial modification of γ-subunit regulation has evolved repeatedly.

TOR shows a similar combination of retention and divergence in Apicomplexa. A TOR ortholog is retained in a minority of apicomplexans, eight of the 41 sampled species, and coccidians among them have an expanded HEAT-repeat region and lack the TORC1 and TORC2 scaffolds RAPTOR and RICTOR (Figures 1B and 3; Supplementary Table S2). HEAT-region divergence correlates with scaffold loss. Our analyses establish this association but do not resolve its mechanism: they show repeated scaffold loss alongside divergence, but not how the absence of RAPTOR and RICTOR affects TOR signaling or whether HEAT-region expansion compensates.

The nucleotide-sensing interface of AMPKγ has also diverged extensively across Apicomplexa (Figures 1E and 2). By contrast, the activating threonine in the AMPKα activation loop is present in 250 of the 254 scored copies, and no species carries only non-threonine copies (Figure 4A-C; Supplementary Table S21). Conservation of the activation loop, even in species without detectable upstream kinases, suggests that some aspects of AMPK regulation remain strongly constrained despite extensive changes elsewhere in the pathway (Supplementary Figure S9D). This contrast argues against a model in which all regulators change equally; different regulatory features appear to face different constraints.

Deleting TgTOR showed that these genomic changes have functional consequences. Three independently generated, sequencing-validated TgTOR-null clones grew and formed plaques indistinguishable from those of wild-type parasites (Figure 7). Since all three independently derived lines showed the same phenotype, an off-target lesion introduced during strain construction is unlikely to explain it.

We are unaware of any other eukaryote reported to complete a replicative cycle after deletion of its sole TOR ortholog. The requirement for TOR in growth is among the best-established findings in eukaryotic cell biology. In *Saccharomyces cerevisiae*, TOR2 is essential, and disruption of both TOR1 and TOR2 arrests cells in G1 (Kunz et al. 1993). In mice, disruption of mTOR is lethal shortly after implantation: proliferation fails in both the embryo proper and the trophoblast, and embryonic stem cells lacking mTOR kinase activity arrest (Murakami et al. 2004). Even in the protozoan Leishmania, which has 4 TOR orthologs, deletion of some orthologs is lethal (Myburgh et al. 2024).

Our findings suggest that conservation of the catalytic ortholog alone is not enough to infer conservation of biological function. At least in *T. gondii*, the TOR ortholog is dispensable for growth under standard culture conditions. However, TgTOR may be needed for *T. gondii* growth under stress, such as nutrient limitation, developmental differentiation, or *in vivo* infection. Yet genome-screen data from nutritionally impaired parasites show no TgTOR-associated phenotype (Rimple et al. 2025). Alternatively, other regulatory systems may carry out functions traditionally attributed to TOR. Distinguishing among these possibilities will require testing TgTOR knockout parasites under nutrient limitation, during differentiation, and animal infection. The viable TOR-null phenotype nonetheless constrains evolutionary interpretations, as do patterns of regulator loss. Within these limits, the comparative measurements and TOR knockout phenotype show that a recognizable ortholog can persist without the regulatory role it has in model systems.

## Materials and Methods

### Candidate discovery and orthology assignment

We identified candidates for ten targets (TOR, RAPTOR, RICTOR, LST8, SIN1, AMPKα, AMPKβ, AMPKγ, LKB1, and the CAMKK/SAK1/GRIK functional category) using two method families: orthogroup inference and domain-based searches. We inferred orthogroups with OrthoFinder (Emms and Kelly 2019), SonicParanoid (Cosentino and Iwasaki 2019), (Supplementary Table S4). For domain-based searches, we used profile HMMs with HMMER (Eddy 2011), CDD signatures with rpsblast (Wang et al. 2023), and InterProScan signatures (Jones et al. 2014; Blum et al. 2025). We fixed settings for each method in advance using a panel of positive controls and decoys (Supplementary Methods; Supplementary Table S5). Fungal and plant AMPK-activating kinases are not orthologous to mammalian CAMKKs (Supplementary Figure S2). We therefore treated the CAMKK/SAK1/GRIK group as a functional category (eggNOG KOG0585; Huerta-Cepas et al. 2019), rather than as an orthogroup (Supplementary Table S34; Supplementary Figure S2D).

For the TOR and AMPK targets, orthology was assigned from gene trees using the per-sequence criterion described by van Dam et al. (2011). Candidates were accepted if they clustered with curated positive controls and excluded if they grouped with a neighboring non-orthologous family marked by a boundary marker (Supplementary Figure S3C). Ambiguous cases were resolved using a two-arm rule: assignments required support from both the orthogroup and domain-based arms. LKB1 and CAMKK assignments relied primarily on candidate discovery; gene trees were used only to exclude candidates that grouped with confusable families (Supplementary Table S35; Supplementary Figure S2C).

Gene trees were inferred with IQ-TREE (Wong et al. 2026) from MAFFT alignments (Katoh and Standley 2013); ModelFinder (Kalyaanamoorthy et al. 2017) selected the substitution model. We also inferred trees under a site-heterogeneous model (Wang et al. 2018) and in RAxML (Stamatakis 2014), and orthology calls under all tree inferences are reported in Supplementary Table S5. TOR alignments were screened for recombination with GARD (Kosakovsky Pond et al. 2006), which detected no recombination events (Supplementary Table S36).

### Species states and composition

Each of the 210 species was assigned one of three states for each target: present, absent, or not assessable. A target was considered present if at least one ortholog was identified. Absence was recorded only for genome-derived proteomes lacking an accepted ortholog; transcriptome-derived proteomes were classified as not assessable (Supplementary Table S7).

We report prevalence in two ways: detection prevalence, calculated across all 210 species, and assessable prevalence, calculated across the 182 genome-derived proteomes. Wilson 95% confidence intervals (Wilson 1927) are provided for assessable prevalence only. Results are tabulated overall and by supergroup (Supplementary Table S1).

Taxonomic composition was summarized at the phylum level, and species without a phylum-level NCBI assignment (Schoch et al. 2020) were reported individually (Supplementary Table S2). Comparisons between phyla included not-assessable species as bounded ranges, and a difference was considered strict when every compatible value fell on the same side of zero (Supplementary Methods; Supplementary Table S2). For phylogenetic analyses, not-assessable states were treated as missing data, and taxa carrying them were pruned when an analysis required complete states. Sensitivity analyses also treated every transcriptome-derived observation as uncertain.

### Sequence phenotypes of orthologous proteins

We defined four sequence phenotypes by comparison with human reference proteins and summarized each species’ value as the median across ortholog copies. Copies that did not meet a metric’s requirements were treated as unavailable.

The AMPKγ contact score quantified conservation across ten AMP-contact residues relative to human PRKAG1. Contact positions were derived from mammalian AMPKγ1 crystal structures (Xiao et al. 2007; Xiao et al. 2011) and grouped into three sets corresponding to AMP-binding sites 1, 3, and 4 (Supplementary Table S37). Site 1 was the primary analysis; sites 3 and 4 served as sensitivity analyses. Evolutionary constraint at contact positions was assessed using rate4site (Pupko et al. 2002).

For AMPKα, we measured the length and predicted disorder of the C-terminal region beyond the residue homologous to PRKAA1 position 290 (Supplementary Table S9). Disorder was predicted using metapredict (Emenecker et al. 2021). Transcriptome-derived sequences were excluded from the primary length analyses. The AMPKβ module score quantified conservation at five carbohydrate-binding module residues relative to human PRKAB1 (Supplementary Table S12).

The TOR N-terminal HEAT region was evaluated in 246 TOR orthologs from 152 species, using sequence signatures, AlphaFold structural models, and InterProScan annotations (Supplementary Table S17). Repeat density was calculated as repeats per kilobase of the mapped region, using one representative copy per species (Supplementary Table S17).

### Structure prediction and AMPKγ pocket geometry

All 239 AMPKγ analysis-set sequences and five control sequences were modeled as single chains using ColabFold with AlphaFold2 (Jumper et al. 2021; Mirdita et al. 2022) (Supplementary Table S15; Supplementary Figure S7). The controls comprised human and rat γ1, yeast Snf4, and two human γ1 variants with substitutions at the ten site 1 AMP-contact residues (Supplementary Figure S7).

Three AMP-binding sites were identified in a rat γ1 structure (Xiao et al. 2007). A reference AMP molecule was transferred to each model, and pocket geometry and local prediction confidence were measured for each site (Supplementary Methods; Supplementary Table S15; Supplementary Figure S7). No confidence threshold was applied because confidence was itself an outcome variable. Values that could not be measured were recorded as unavailable. Human γ1 was also modeled with AlphaFold-Multimer as a benchmark, and its measurements were analyzed separately (Supplementary Figures S14 and S15).

### Generation and characterization of TgTOR knockout strains

*Toxoplasma gondii* was maintained in human fibroblasts (ATCC SCRC-1041) cultured in DMEM supplemented with 10% FBS. TgTOR knockout lines were generated in RH parasites that constitutively express Cas9 (ATCC PRA-426). The TgTOR coding sequence (TGME49_316430) was replaced by inserting a guide vector using two previously described guide RNAs (Sidik et al. 2016). Genomic sequencing confirmed the intended deletion in three independently derived clones (7, 15, and 17; Figure 7A,B).

TOR transcript abundance was measured by qPCR in three biological replicates per clone, with TgSAG1 (TGME49_233460) as the reference transcript, and is reported relative to wild type. The 117 bp TgTOR amplicon falls within the deleted interval and therefore cannot be amplified from the edited locus. Reactions with no detectable amplification were assigned a Cq value of 40, making the reported depletion a lower bound. Clones 15 and 17 both reached this floor; thus, the difference between their bounds (1,687-fold and 671-fold, or 1.3 cycles) reflects variation in the TgSAG1 reference rather than in TgTOR.

Replication was assessed by infecting confluent human fibroblast monolayers for 2 days. Cultures were then fixed, stained with DAPI and anti-TgSAG1 antibody, and scored for parasites per vacuole (50 vacuoles per strain across three replicates). Each knockout was compared with wild type using Student’s t-test, with vacuoles treated as independent observations. This approach inflates the effective sample size and narrows the confidence interval for the difference, making the failure to detect a difference conservative. Long-term lytic growth was measured by plaque assay (Messina et al. 2025), with triplicate wells inoculated with 50, 500, or 5000 parasites. Wells seeded with 5000 parasites had completely lysed by day 7 and were excluded from the analysis. Plaque area under the two conditions was analyzed using Tukey-adjusted pairwise comparisons, which found no statistically significant difference.

### Use of generative artificial intelligence

A large language model, Anthropic Claude Opus 5, accessed through Claude Code, assisted with writing and adapting the analysis and figure-rendering code described here; it was not used for any other part of this work. It did not select the analyses, establish the coding rules, or determine any classification reported here. The authors supervised, reviewed, and verified all code it produced or adapted, and take responsibility for its content. The authors also used Grammarly (Grammarly Inc.) to correct grammar, spelling, and punctuation during manuscript preparation. They reviewed all suggestions and accepted or rejected them, and take full responsibility for the publication’s content.

## Data Availability

Supplementary data are provided in the article’s single workbook, and figure source data accompany the figures. Analysis code is publicly available at https://github.com/Brunomdg/AMPK-TOR-pipeline. The repository includes the pipeline, helper scripts, documentation, software and environment details, penalized-likelihood search settings, and command lines for each external program run. Job scripts, figure and table generation, independent recomputation checks, and supplementary-only analyses are available from the authors upon request.

The remaining data are deposited in Zenodo (Patwardhan et al. 2026). These include the 210-proteome index and source details; orthology calls; species-state matrix; alignments and gene and dated species trees; and profile panel and control sequences. The fitted models, ancestral-state reconstructions, convergence tests and structure-prediction summaries are reported in the supplementary tables, and the underlying objects are available from the corresponding author on request. Proteomes were obtained from the public resources listed in the references: NCBI RefSeq and GenBank, EukProt, VEuPathDB, ParameciumDB, UniProt, Ensembl Genomes, and MMETSP; a further eight proteomes were provided by their authors (Supplementary Table S32).

## Supplementary Material

The Supplementary Material comprises the Supplementary Methods, 16 supplementary figures and 44 supplementary tables. The tables are supplied as a single Excel workbook, Supplementary Data, which opens with a Contents sheet listing Tables S1 to S44 in numerical order with their titles and legends, followed by one worksheet per table or table part, named by its table number.

## Acknowledgments

We thank the Vermont Advanced Computing Center (VACC) at the University of Vermont for the computing resources used in this study, and the VACC staff who maintain them. This work was supported by the National Institute of General Medical Sciences (NIGMS), National Institutes of Health under award P20GM125498 and by the U.S. National Science Foundation under award 2510406.

