## Supplementary material for "Divergence of TOR and AMPK regulators across 210 eukaryotes predicts a dispensable TOR in Toxoplasma gondii": S1

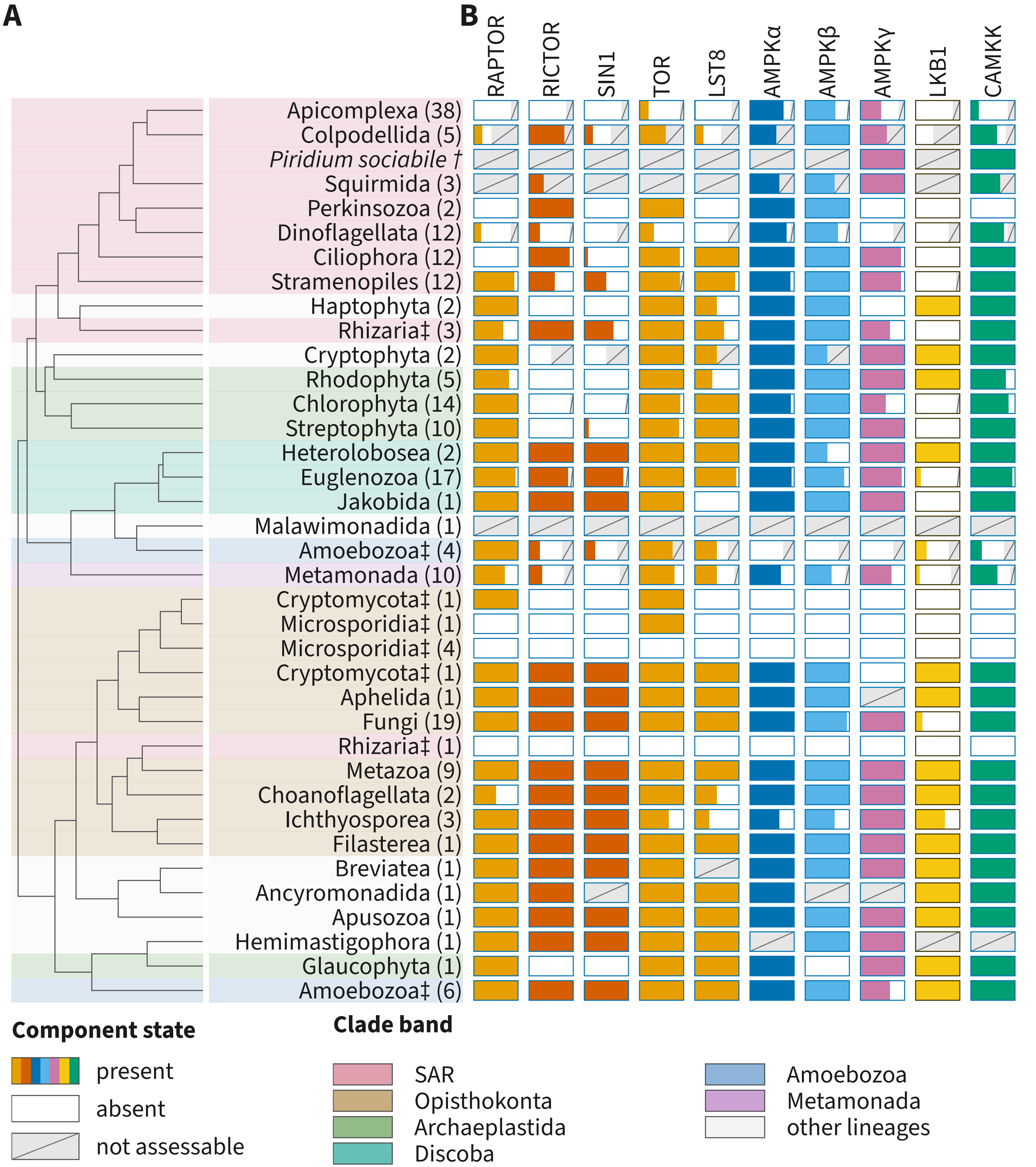
A
B
RAPTOR
RICTOR
SIN1
TOR
LST8
AMPKα
AMPKβ
AMPKγ
LKB1
CAMKK
Apicomplexa (38)
Colpodellida (5)
*Piridium sociabile* †
Squirmida (3)
Perkinsozoa (2)
Dinoflagellata (12)
Ciliophora (12)
Stramenopiles (12)
Haptophyta (2)
Rhizaria‡ (3)
Cryptophyta (2)
Rhodophyta (5)
Chlorophyta (14)
Streptophyta (10)
Heterolobosea (2)
Euglenozoa (17)
Jakobida (1)
Malawimonadida (1)
Amoebozoa‡ (4)
Metamonada (10)
Cryptomycota‡ (1)
Microsporidia‡ (1)
Microsporidia‡ (4)
Cryptomycota‡ (1)
Aphelida (1)
Fungi (19)
Rhizaria‡ (1)
Metazoa (9)
Choanoflagellata (2)
Ichthyosporea (3)
Filasterea (1)
Breviatea (1)
Ancyromonadida (1)
Apusozoa (1)
Hemimastigophora (1)
Glaucophyta (1)
Amoebozoa‡ (6)
Component state
present
absent
not assessable
Clade band
SAR
Opisthokonta
Archaeplastida
Discoba
Amoebozoa
Metamonada
other lineages
