## Supplementary material for "Divergence of TOR and AMPK regulators across 210 eukaryotes predicts a dispensable TOR in Toxoplasma gondii": S2

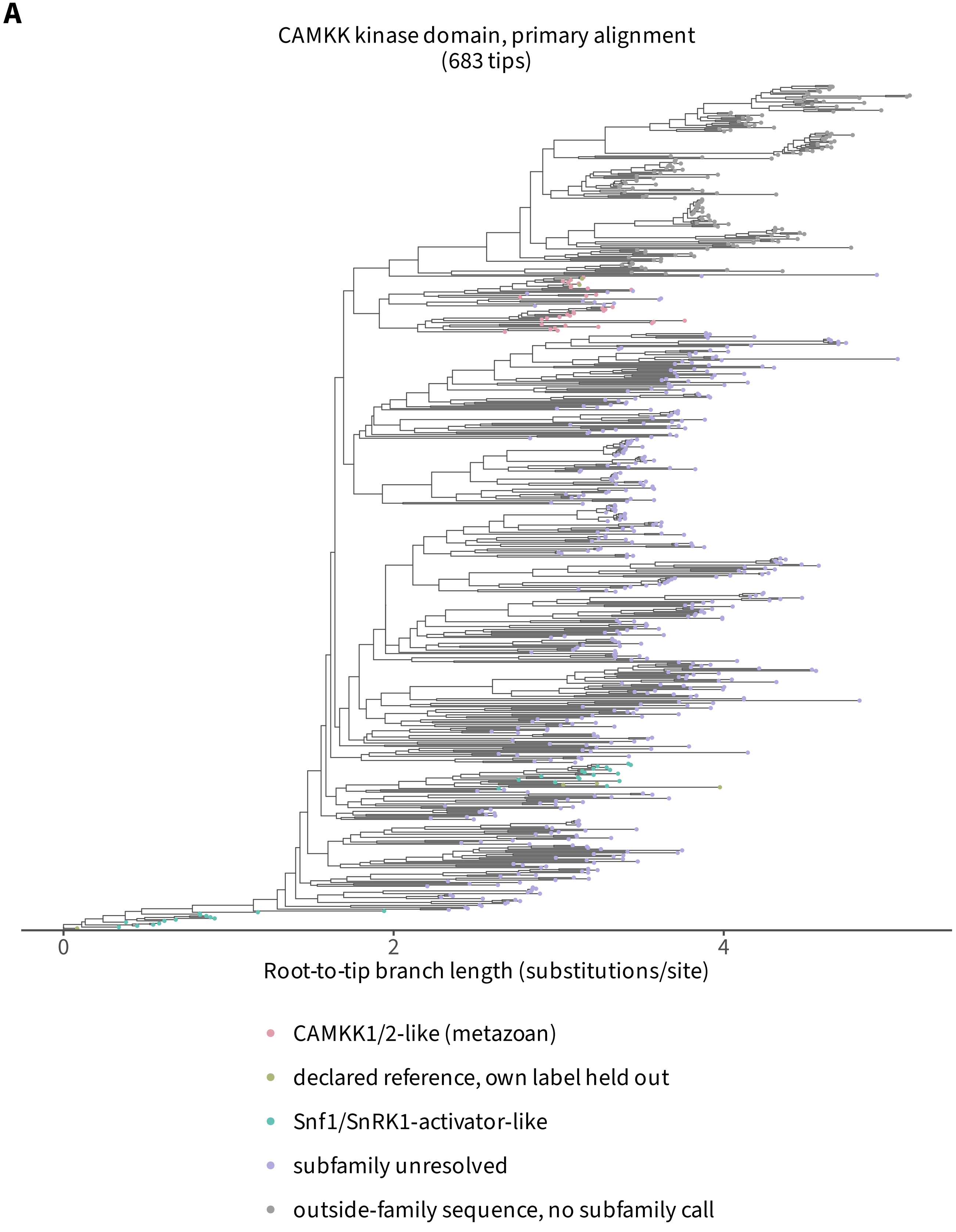
A
CAMKK kinase domain, primary alignment
(683 tips)
0
2
4
Root-to-tip branch length (substitutions/site)
CAMKK1/2-like (metazoan)
declared reference, own label held out
Snf1/SnRK1-activator-like
subfamily unresolved
outside-family sequence, no subfamily call

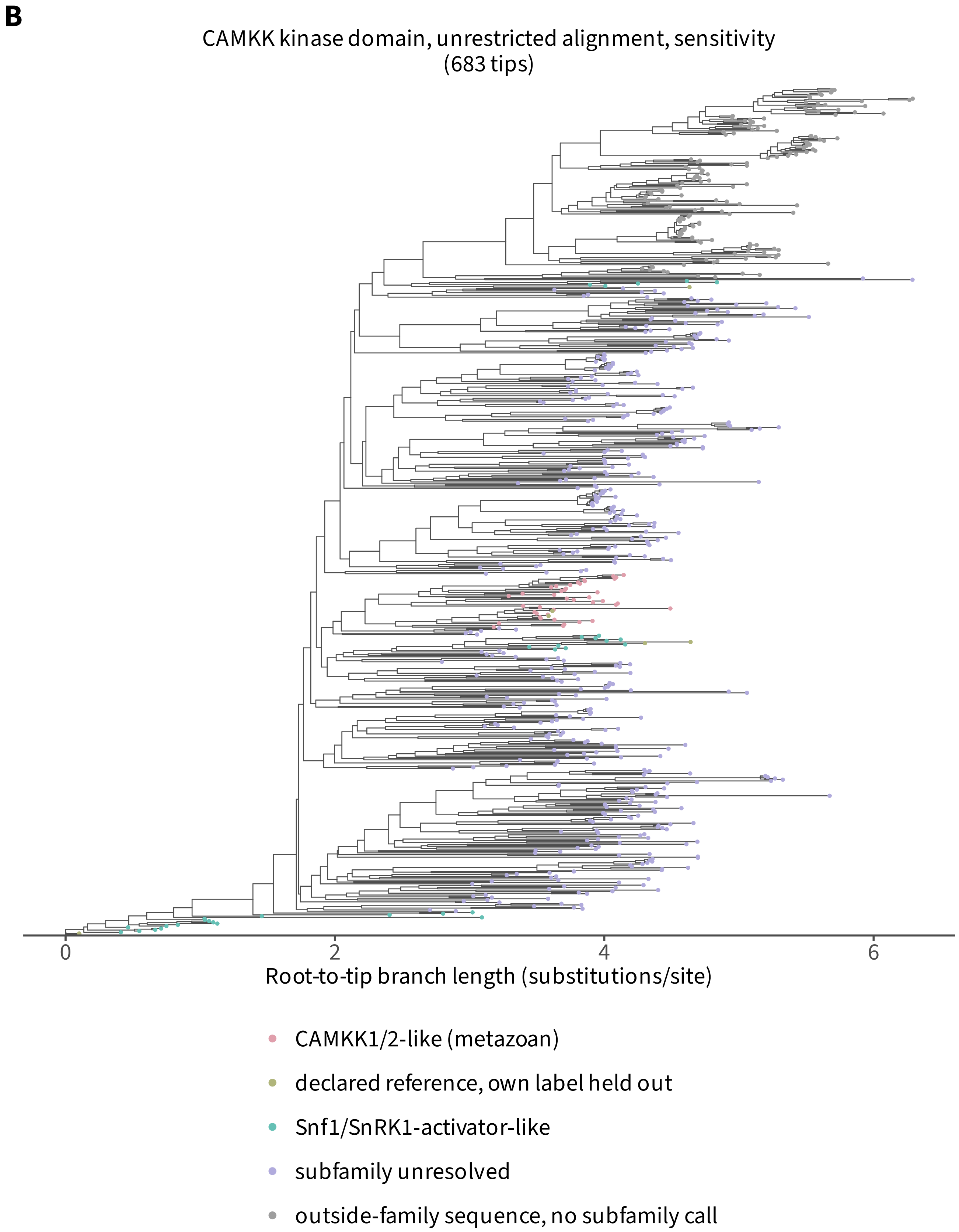

B
CAMKK kinase domain, unrestricted alignment, sensitivity
(683 tips)
0
2
4
6
Root-to-tip branch length (substitutions/site)
CAMKK1/2-like (metazoan)
declared reference, own label held out
Snf1/SnRK1-activator-like
subfamily unresolved
outside-family sequence, no subfamily call

c

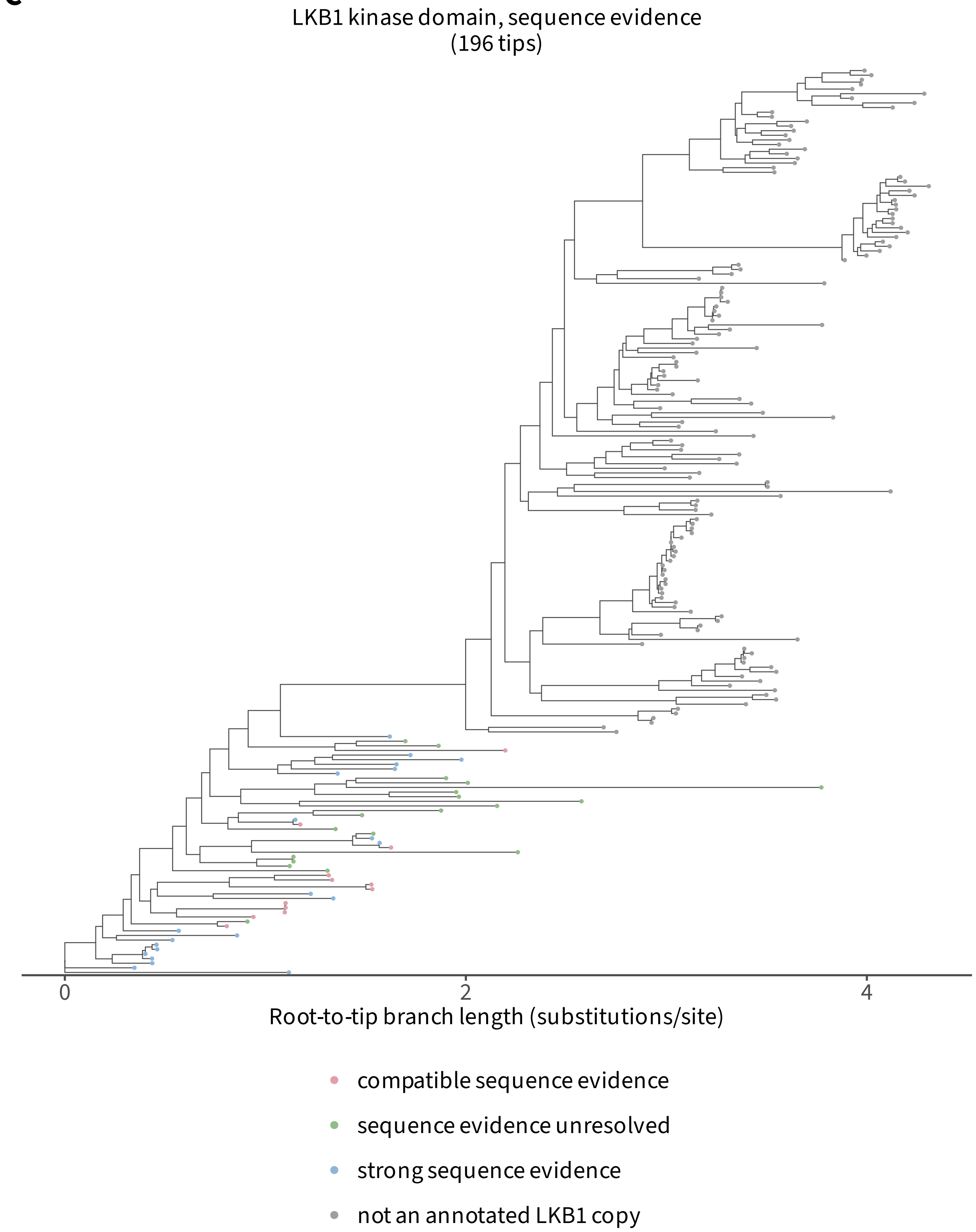

LKB1 kinase domain, sequence evidence
(196 tips)
0
2
4
Root-to-tip branch length (substitutions/site)
compatible sequence evidence
sequence evidence unresolved
strong sequence evidence
not an annotated LKB1 copy

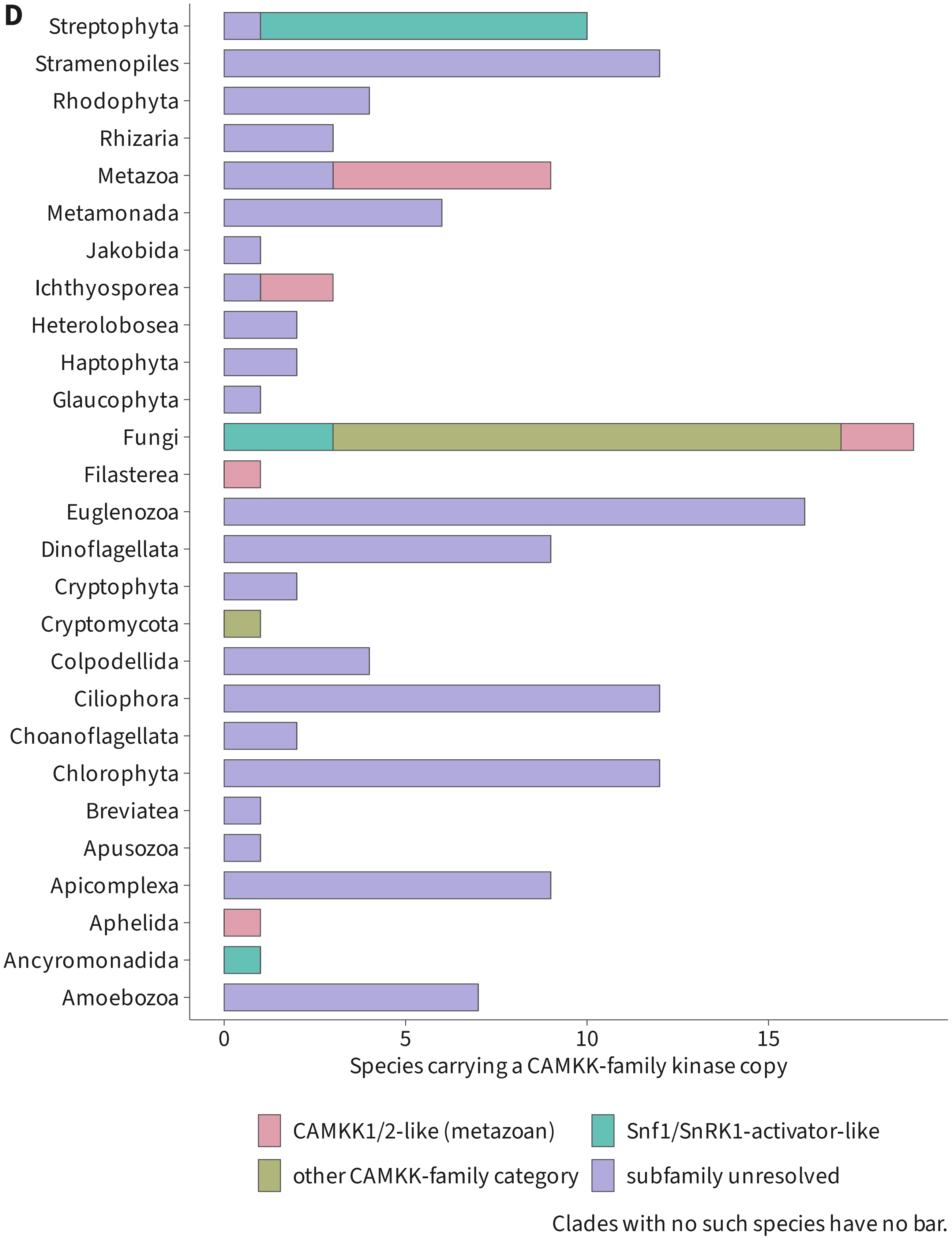
D
Streptophyta
Stramenopiles
Rhodophyta
Rhizaria
Metazoa
Metamonada
Jakobida
Ichthyosporea
Heterolobosea
Haptophyta
Glaucophyta
Fungi
Filasterea
Euglenozoa
Dinoflagellata
Cryptophyta
Cryptomycota
Colpodellida
Ciliophora
Choanoflagellata
Chlorophyta
Breviatea
Apusozoa
Apicomplexa
Aphelida
Ancyromonadida
Amoebozoa
0
5
10
15
Species carrying a CAMKK-family kinase copy
CAMKK1/2-like (metazoan)
Snf1/SnRK1-activator-like
other CAMKK-family category
subfamily unresolved
Clades with no such species have no bar.

### E Reference reassignment

| Reference | Declared class | Recovered class | Support (%) | Placement |
| --- | --- | --- | --- | --- |
| NP_200863.2 | Snf1/SnRK1 activator-like | Snf1/SnRK1 activator-like | 100.0 | supported |
| NP_566876.3 | Snf1/SnRK1 activator-like | Snf1/SnRK1 activator-like | 100.0 | supported |
| NP_757343.2 | CAMKK1/2-like (metazoan) | CAMKK1/2-like (metazoan) | 96.0 | supported |
| XP_047284057.1 | CAMKK1/2-like (metazoan) | CAMKK1/2-like (metazoan) | 96.0 | supported |
| NP_011055.3 | Snf1/SnRK1 activator-like | Snf1/SnRK1 activator-like | 91.0 | provisional placement |
| NP_011336.3 | Snf1/SnRK1 activator-like | Snf1/SnRK1 activator-like | 91.0 | provisional placement |
| NP_012876.1 | Snf1/SnRK1 activator-like | Snf1/SnRK1 activator-like | 99.0 | supported |

### F Tree inference settings

| Tree / alignment | Model | Bootstrap convergence | Tips |
| --- | --- | --- | --- |
| CAMKK, primary alignment (A) | LG+R10 | not converged | 683 |
| CAMKK, unrestricted alignment (B) | LG+I+R9 | not converged | 683 |
| LKB1, primary alignment | LG+I+R6 | converged | 196 |
| LKB1, unrestricted alignment (C) | LG+R7 | converged | 196 |
