## Supplementary material for "Divergence of TOR and AMPK regulators across 210 eukaryotes predicts a dispensable TOR in Toxoplasma gondii": S3

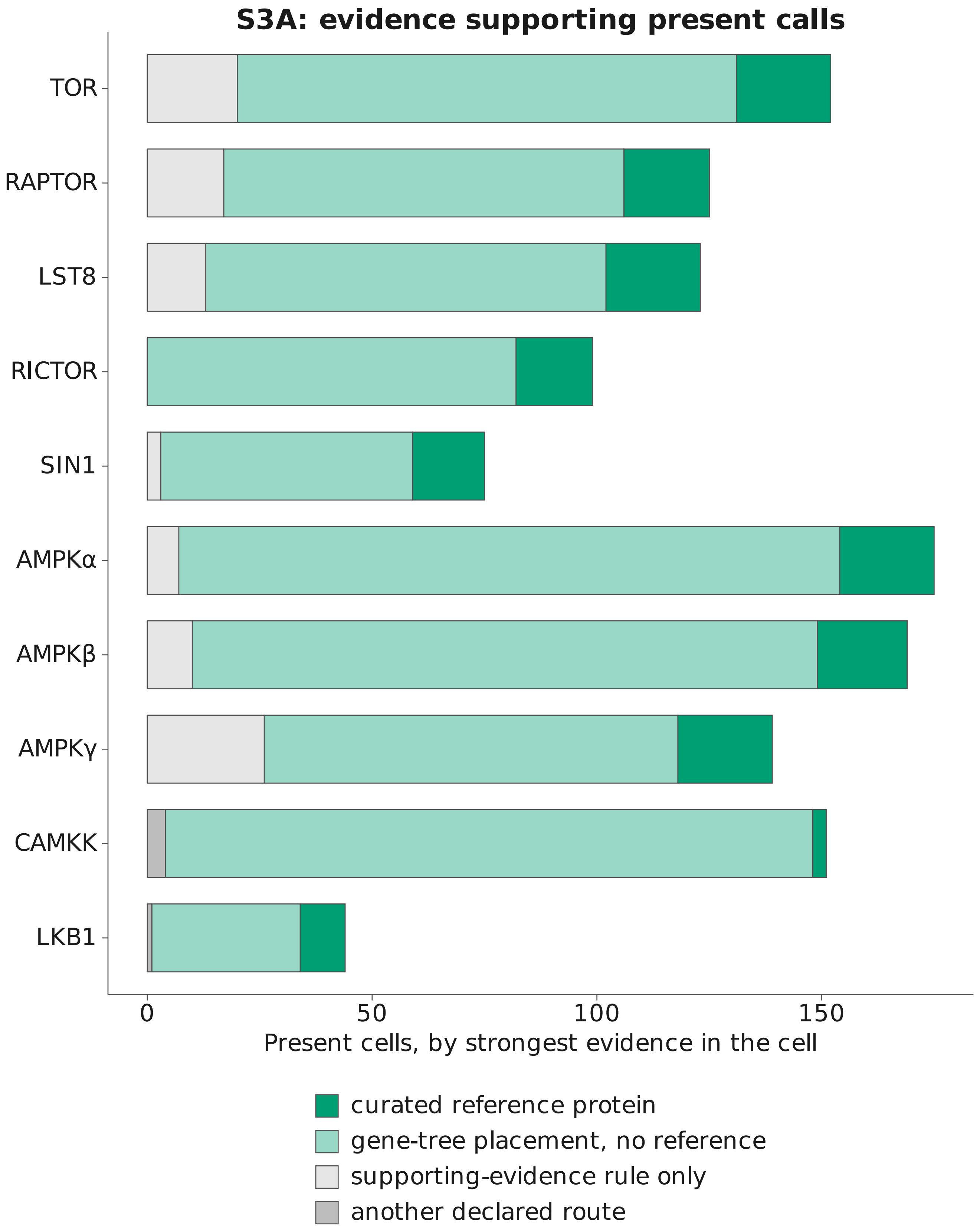
S3A: evidence supporting present calls
TOR
RAPTOR
LST8
RICTOR
SIN1
AMPKα
AMPKβ
AMPKγ
CAMKK
LKB1
0
50
100
150
Present cells, by strongest evidence in the cell
curated reference protein
gene-tree placement, no reference
supporting-evidence rule only
another declared route

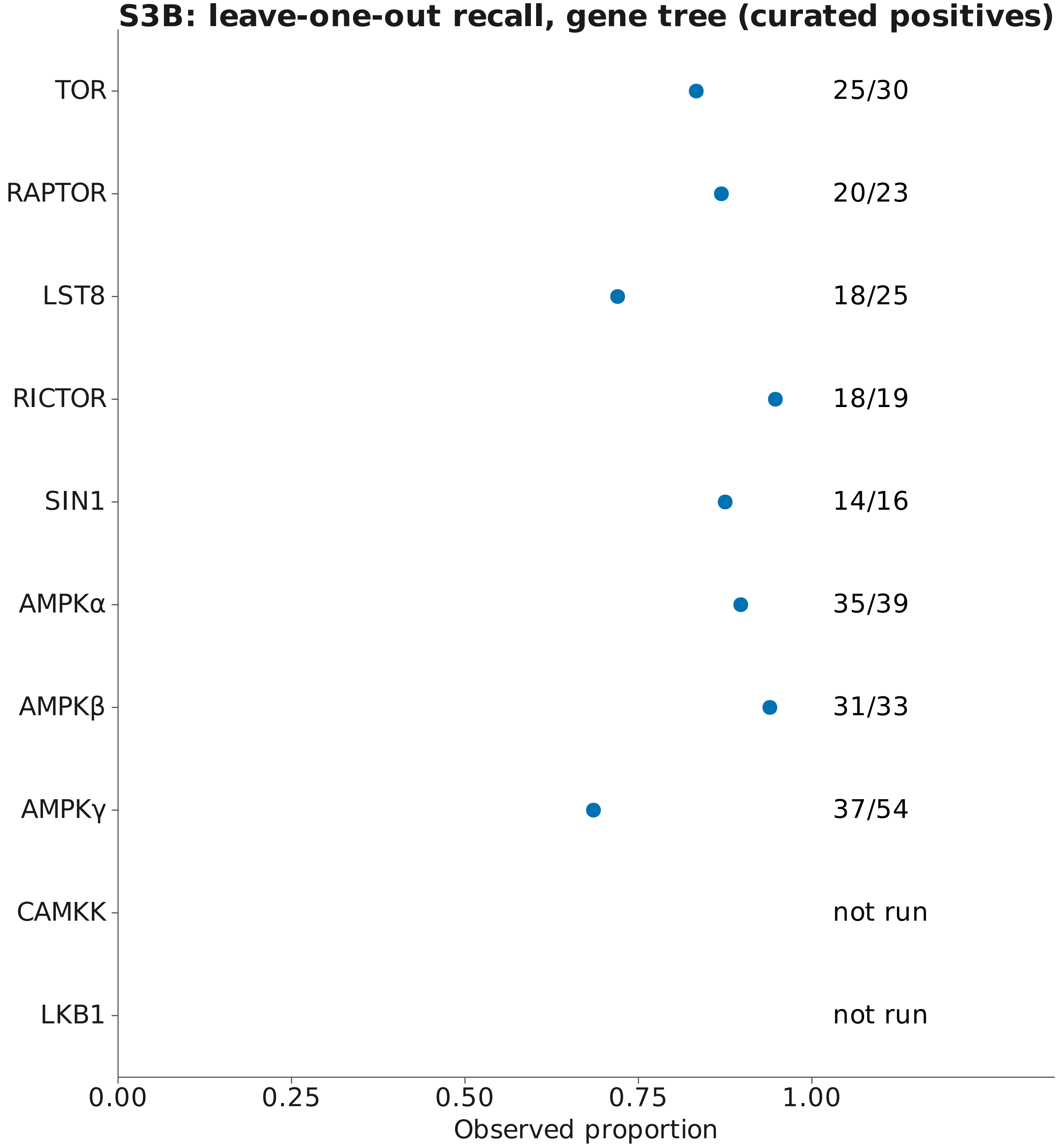

Fractions show the available numerator and denominator.
Cross-check percentages do not independently settle a call.

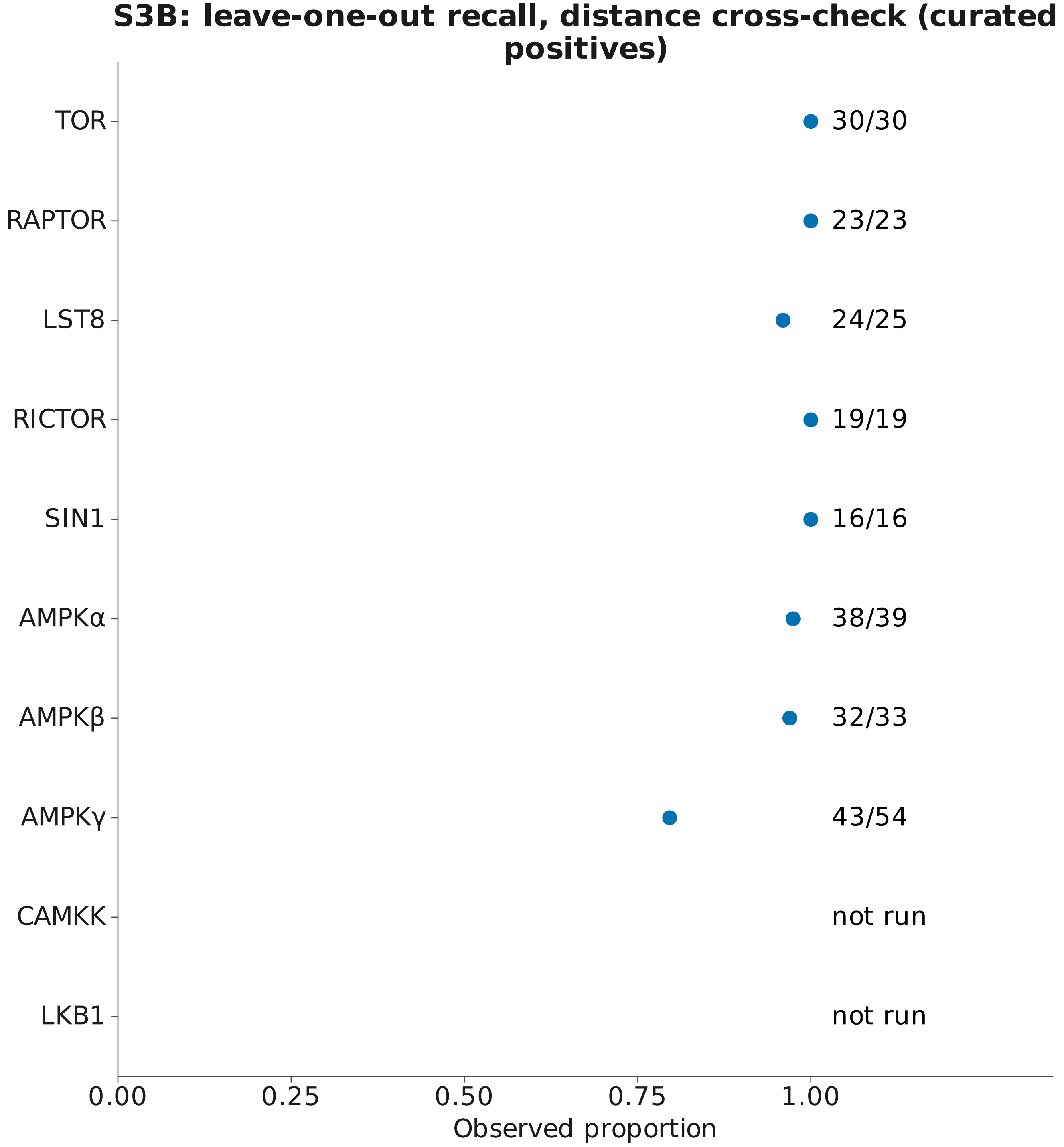


Fractions show the available numerator and denominator.
Cross-check percentages do not independently settle a call.

### S3B: leave-one-out specificity, gene tree (boundary markers)

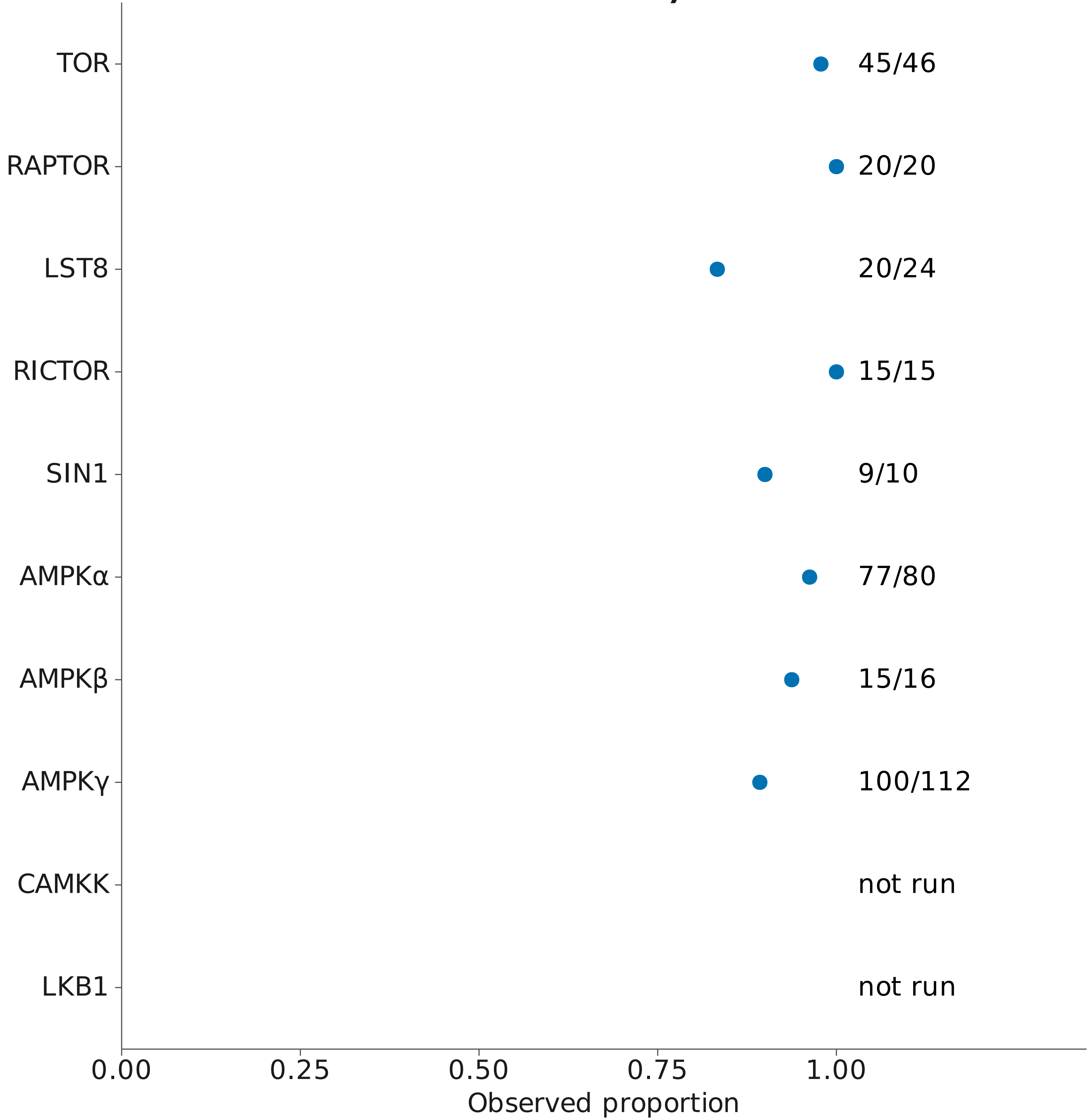


Fractions show the available numerator and denominator.
Cross-check percentages do not independently settle a call.

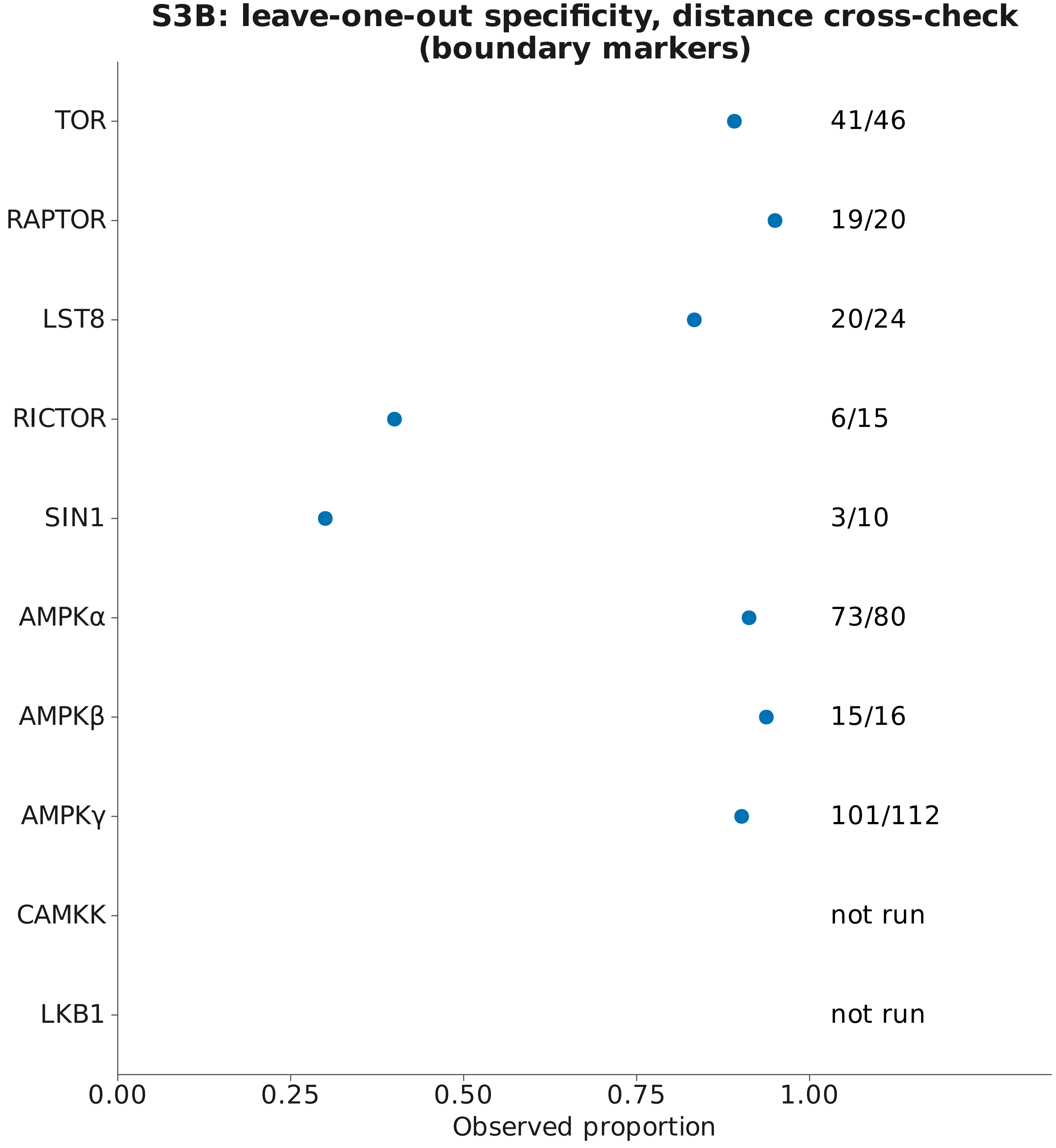

Fractions show the available numerator and denominator.
Cross-check percentages do not independently settle a call.

### S3B: RAxML vs IQ-TREE agreement (cross-check; settles no call)

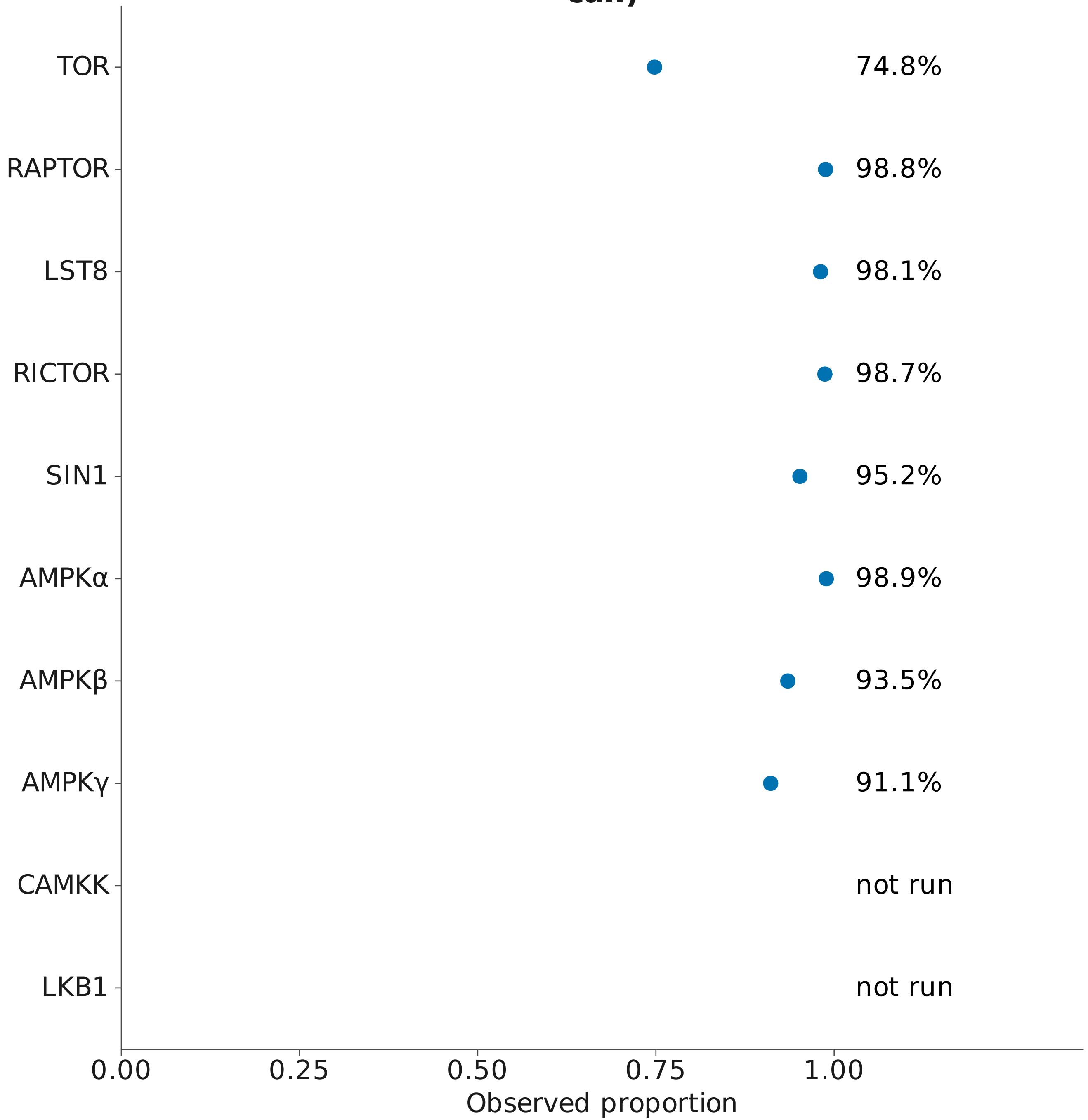


Fractions show the available numerator and denominator.
Cross-check percentages do not independently settle a call.

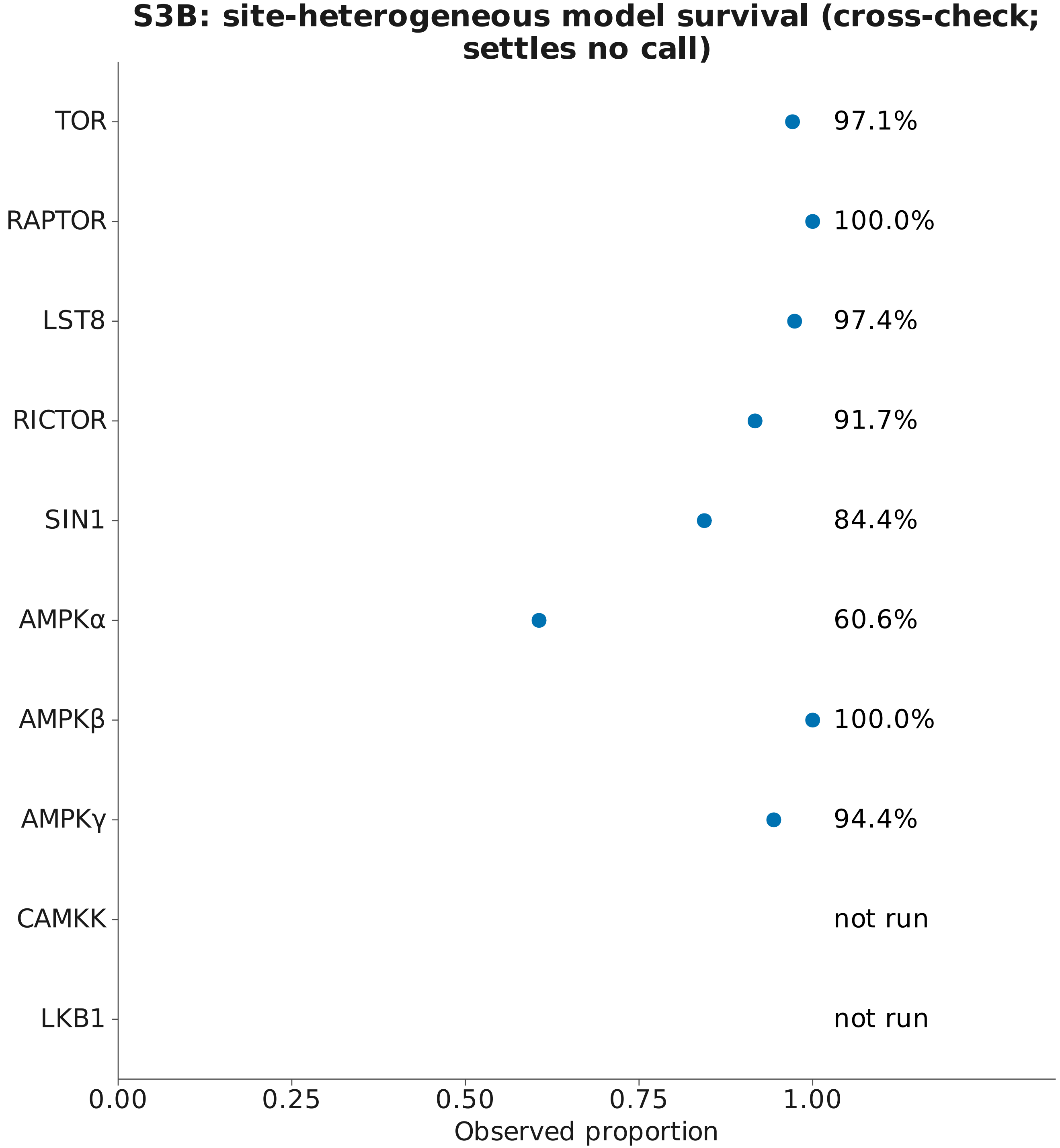


Fractions show the available numerator and denominator.
Cross-check percentages do not independently settle a call.

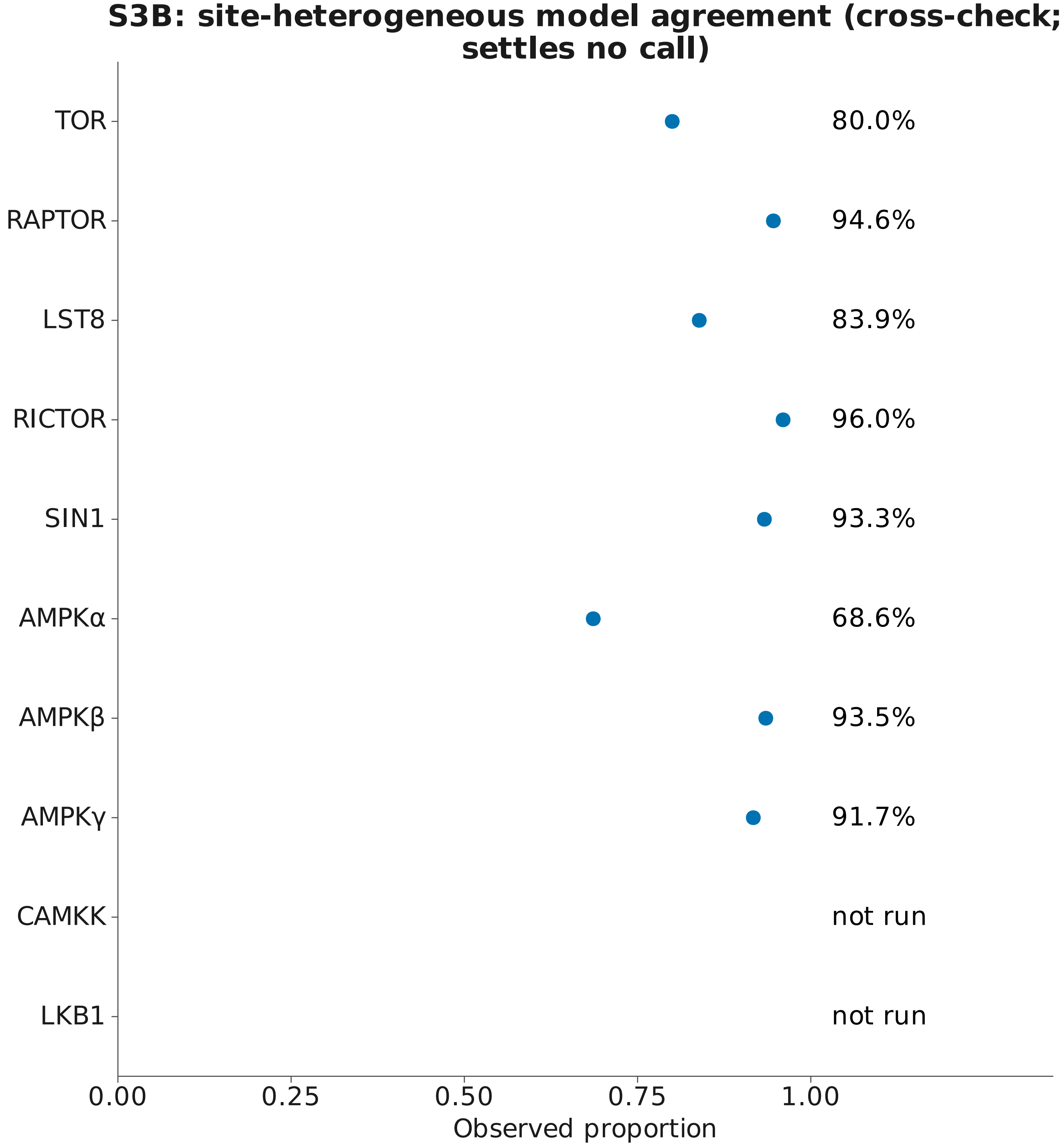


Fractions show the available numerator and denominator.
Cross-check percentages do not independently settle a call.

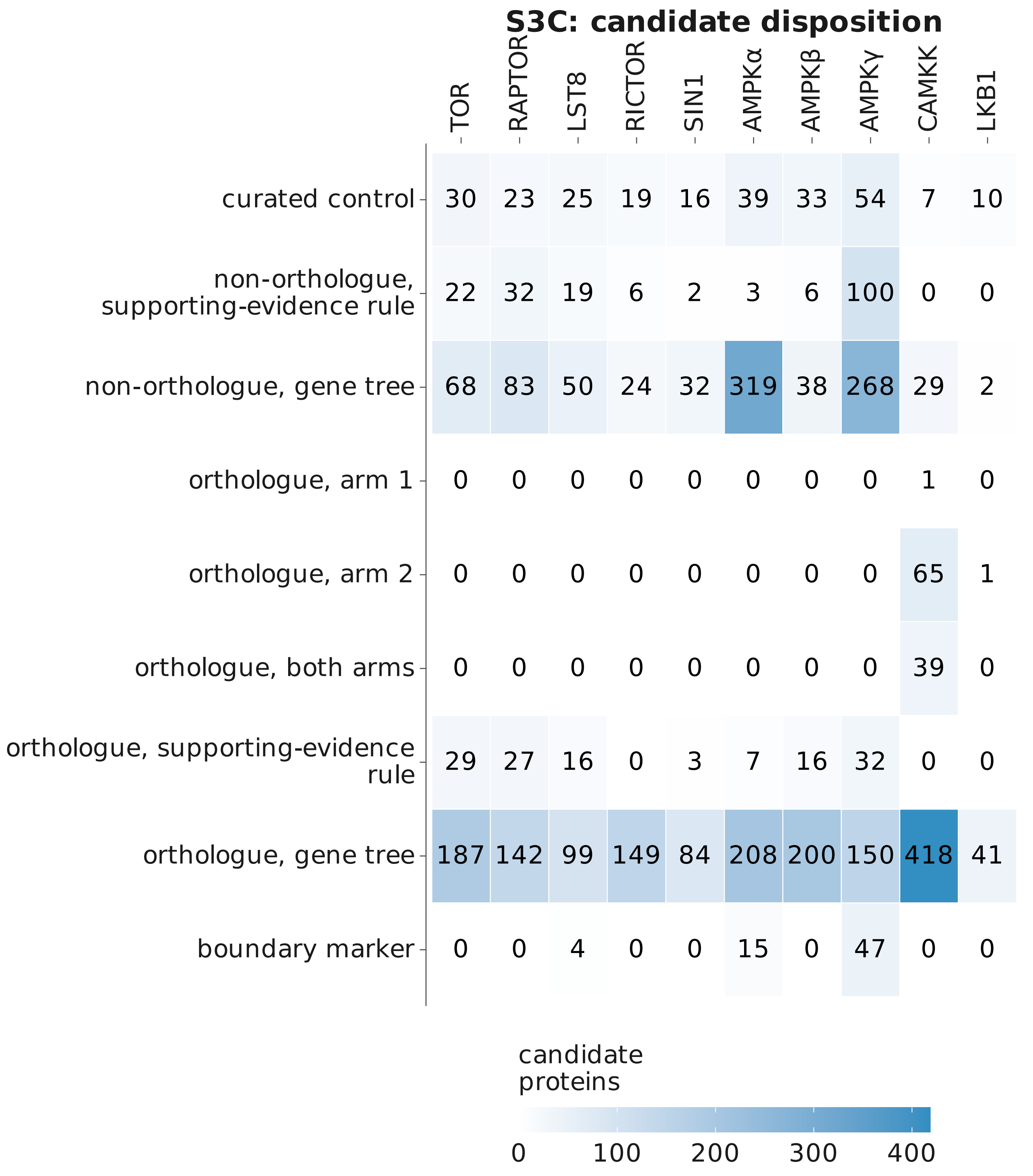
S3C: candidate disposition
TOR
RAPTOR
LST8
RICTOR
SIN1
AMPKα
AMPKβ
AMPKγ
CAMKK
LKB1
curated control
30 23 25 19 16 39 33 54 7 10
non-orthologue, supporting-evidence rule
22 32 19 6 2 3 6 100 0 0
non-orthologue, gene tree
68 83 50 24 32 319 38 268 29 2
orthologue, arm 1
0 0 0 0 0 0 0 0 1 0
orthologue, arm 2
0 0 0 0 0 0 0 0 65 1
orthologue, both arms
0 0 0 0 0 0 0 0 39 0
orthologue, supporting-evidence rule
29 27 16 0 3 7 16 32 0 0
orthologue, gene tree
187 142 99 149 84 208 200 150 418 41
boundary marker
0 0 4 0 0 15 0 47 0 0
candidate proteins
0
100
200
300
400

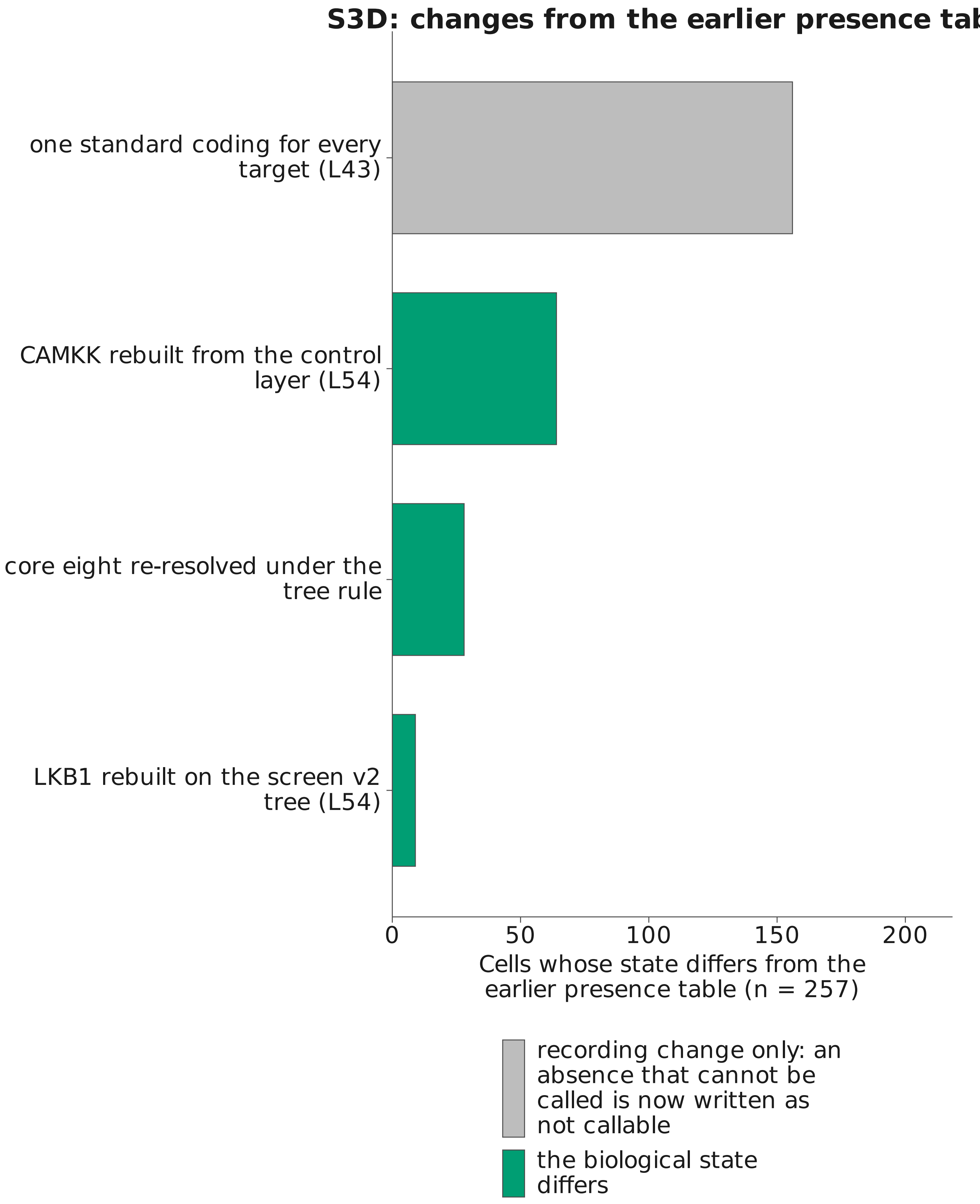
S3D: changes from the earlier presence tab
one standard coding for every target (L43)
CAMKK rebuilt from the control layer (L54)
core eight re-resolved under the tree rule
LKB1 rebuilt on the screen v2 tree (L54)
0
50
100
150
200
Cells whose state differs from the earlier presence table (n = 257)
recording change only: an absence that cannot be called is now written as not callable
the biological state differs

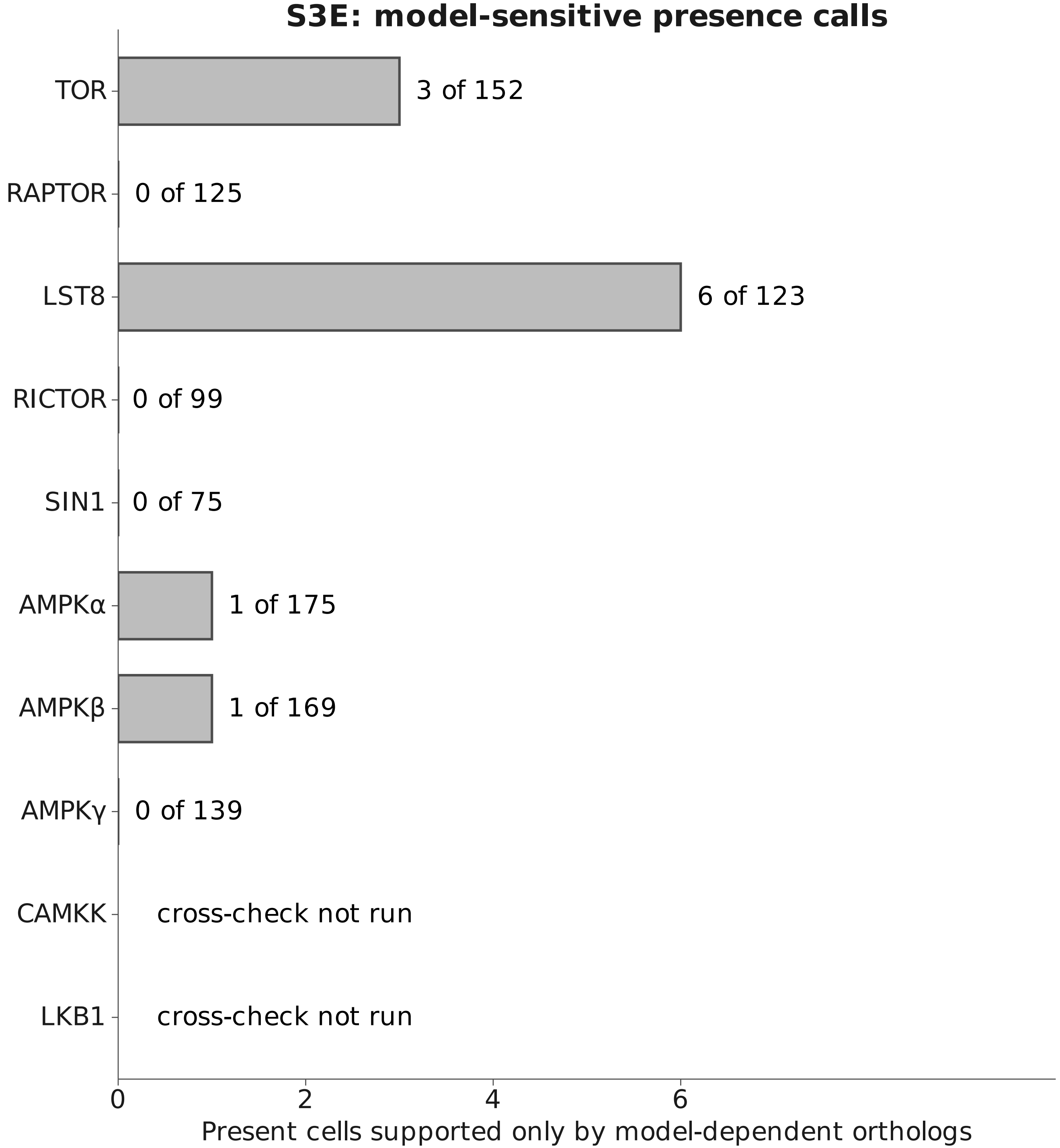

Labels: model-sensitive cells / present cells
Targets without this cross-check are explicitly marked.
