## Supplementary material for "Divergence of TOR and AMPK regulators across 210 eukaryotes predicts a dispensable TOR in Toxoplasma gondii": S4

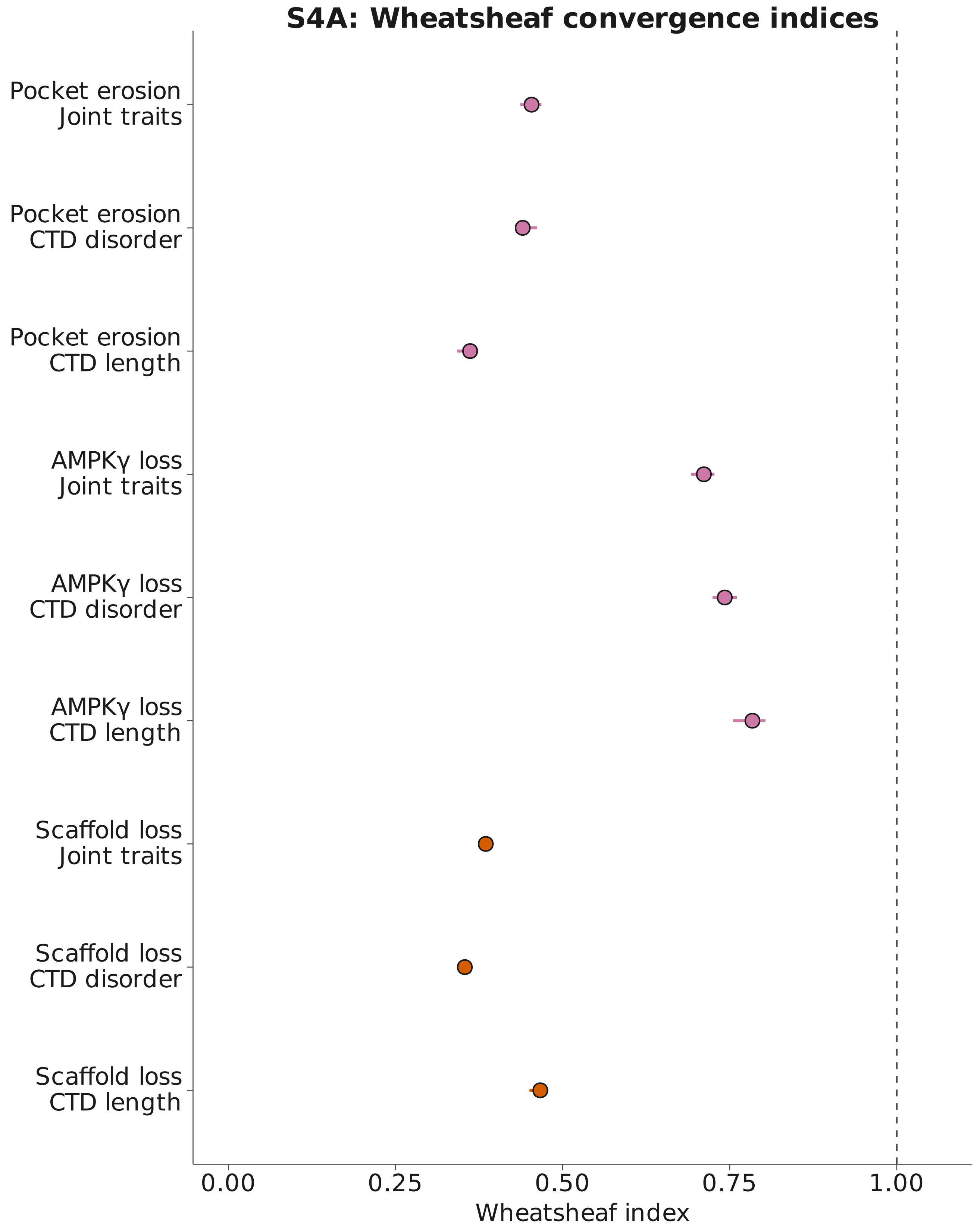


Bars: package-reported 95% intervals. Dashed line: index 1.
Traits use log length or logit disorder; joint analysis uses both.
Failure to detect convergence does not establish equivalence.

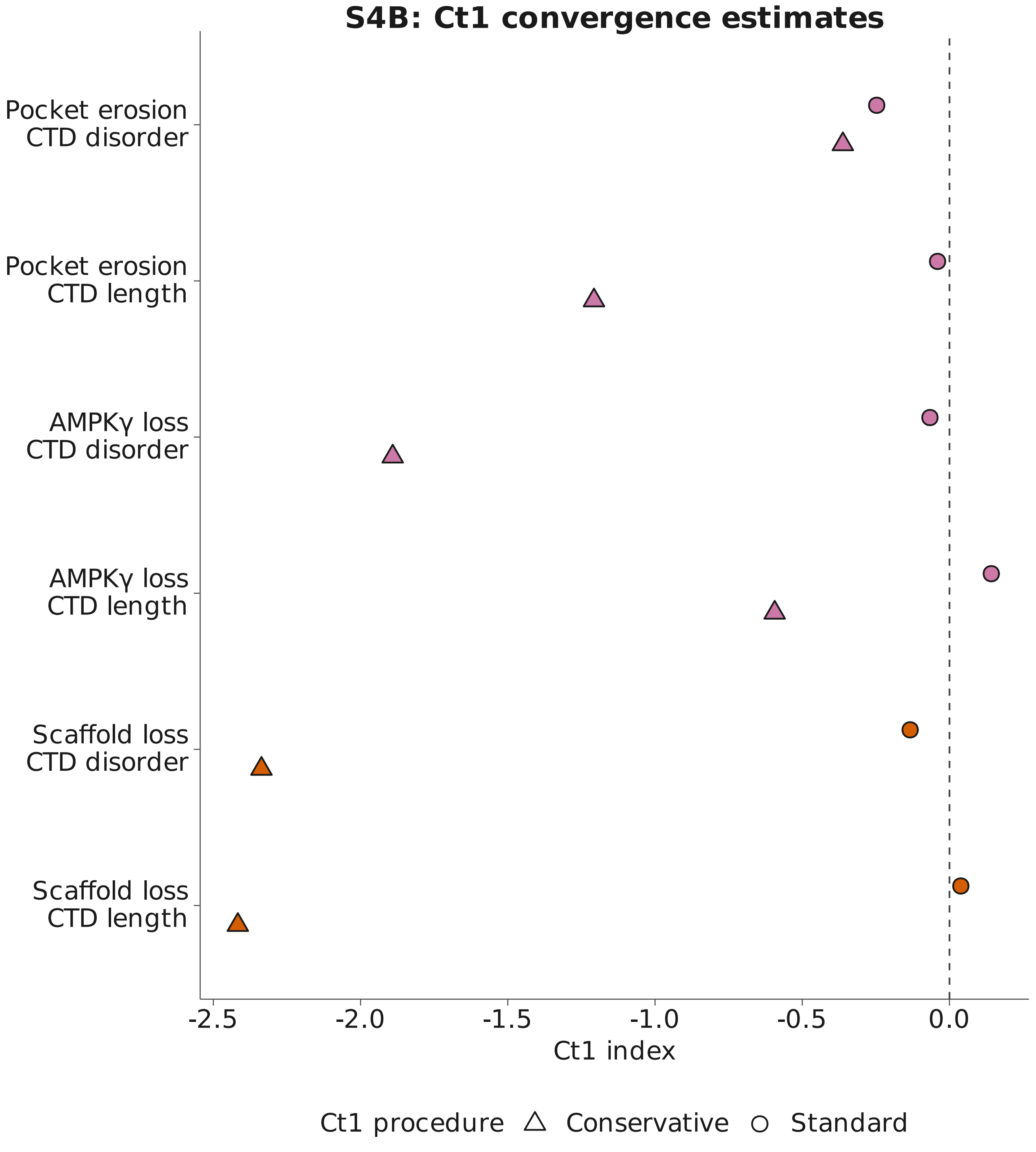


No intervals are available for these Ct1 estimates.
Dashed line: zero. Exact sample sizes and nominal P values are retained in the accompanying source tables.
