## Supplementary material for "Divergence of TOR and AMPK regulators across 210 eukaryotes predicts a dispensable TOR in Toxoplasma gondii": S5

#### S5: site1, pairwise mapping

All 239 analysis-set sequences; recorded positional counts

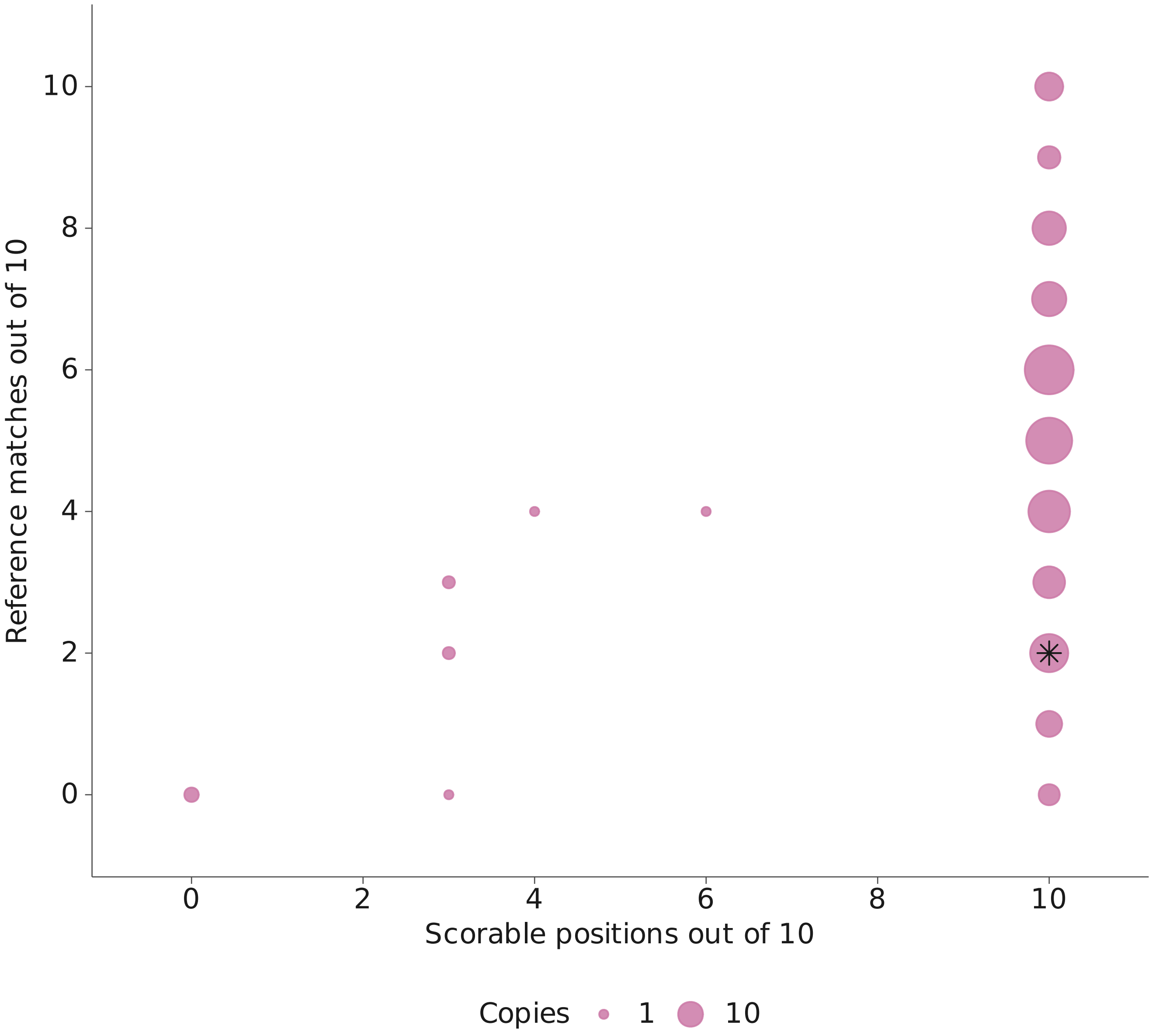


Point area: number of copies. Black star: Toxoplasma gondii.
Zero scorable positions indicates mapping coverage, not loss.
Position sets and alignment procedures remain separate.

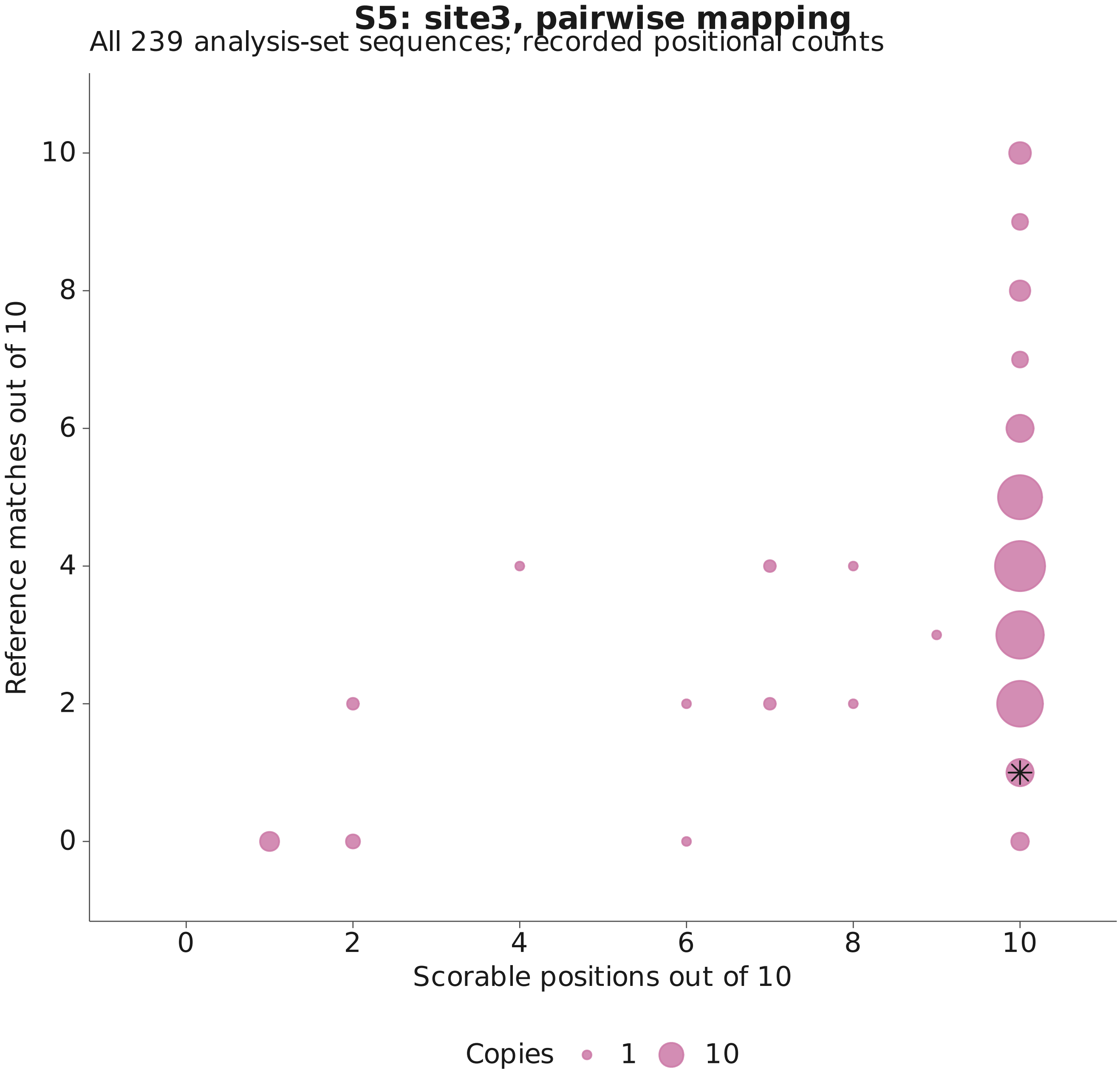


Point area: number of copies. Black star: Toxoplasma gondii.
Zero scorable positions indicates mapping coverage, not loss.
Position sets and alignment procedures remain separate.

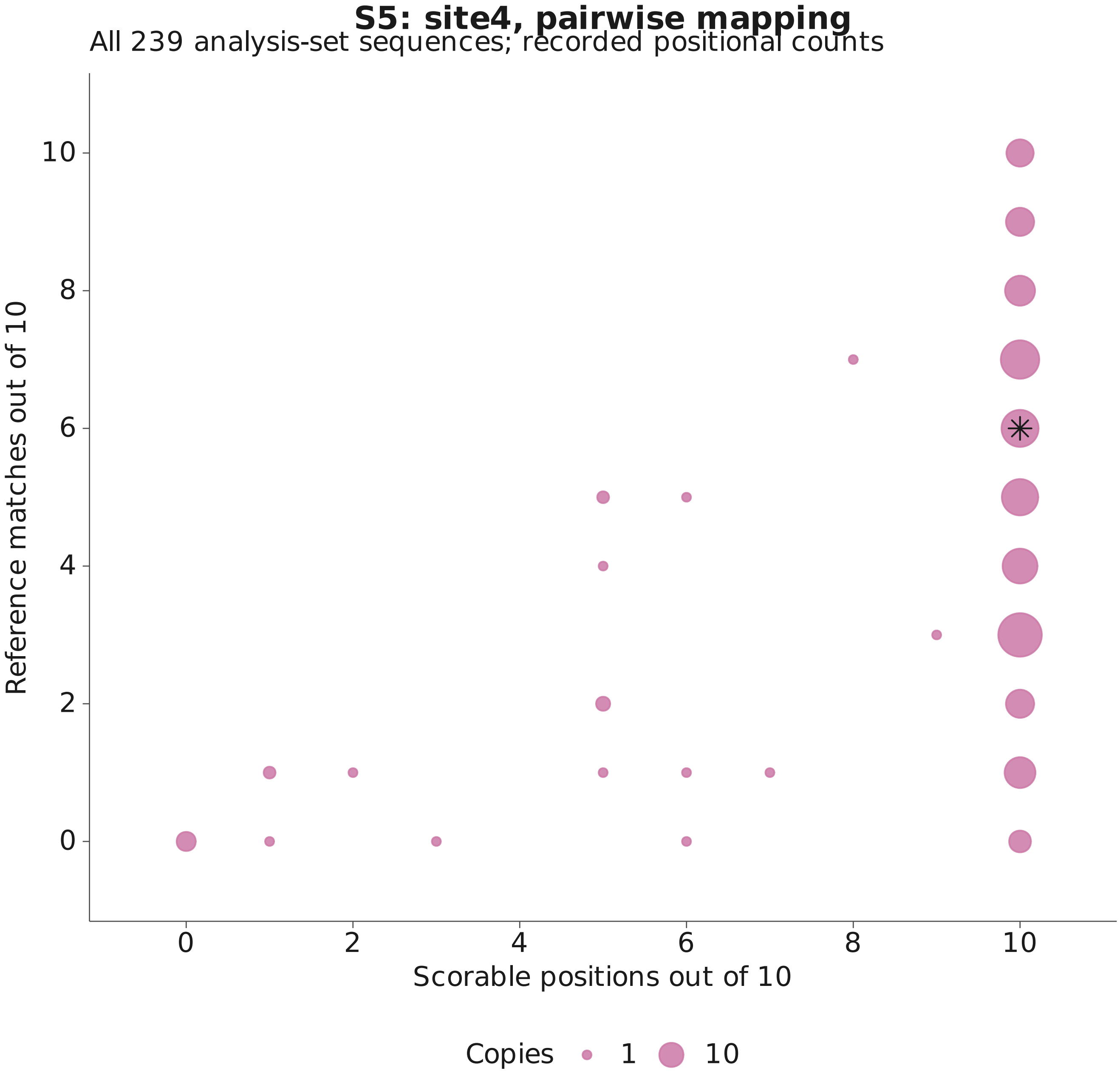


Point area: number of copies. Black star: Toxoplasma gondii.
Zero scorable positions indicates mapping coverage, not loss.
Position sets and alignment procedures remain separate.

#### S5: site1, joint mapping

All 239 analysis-set sequences; recorded positional counts

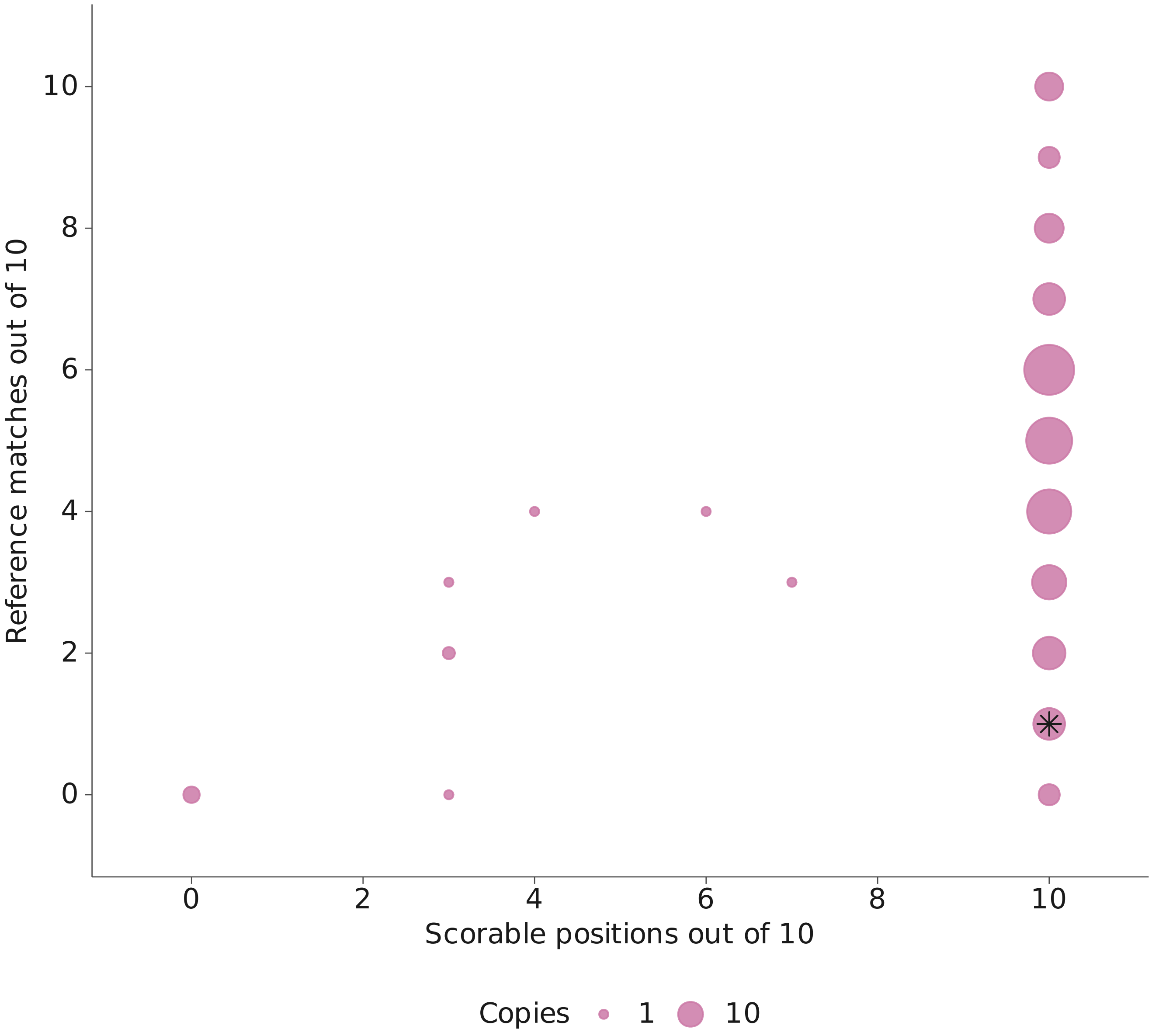


Point area: number of copies. Black star: Toxoplasma gondii.
Zero scorable positions indicates mapping coverage, not loss.
Position sets and alignment procedures remain separate.

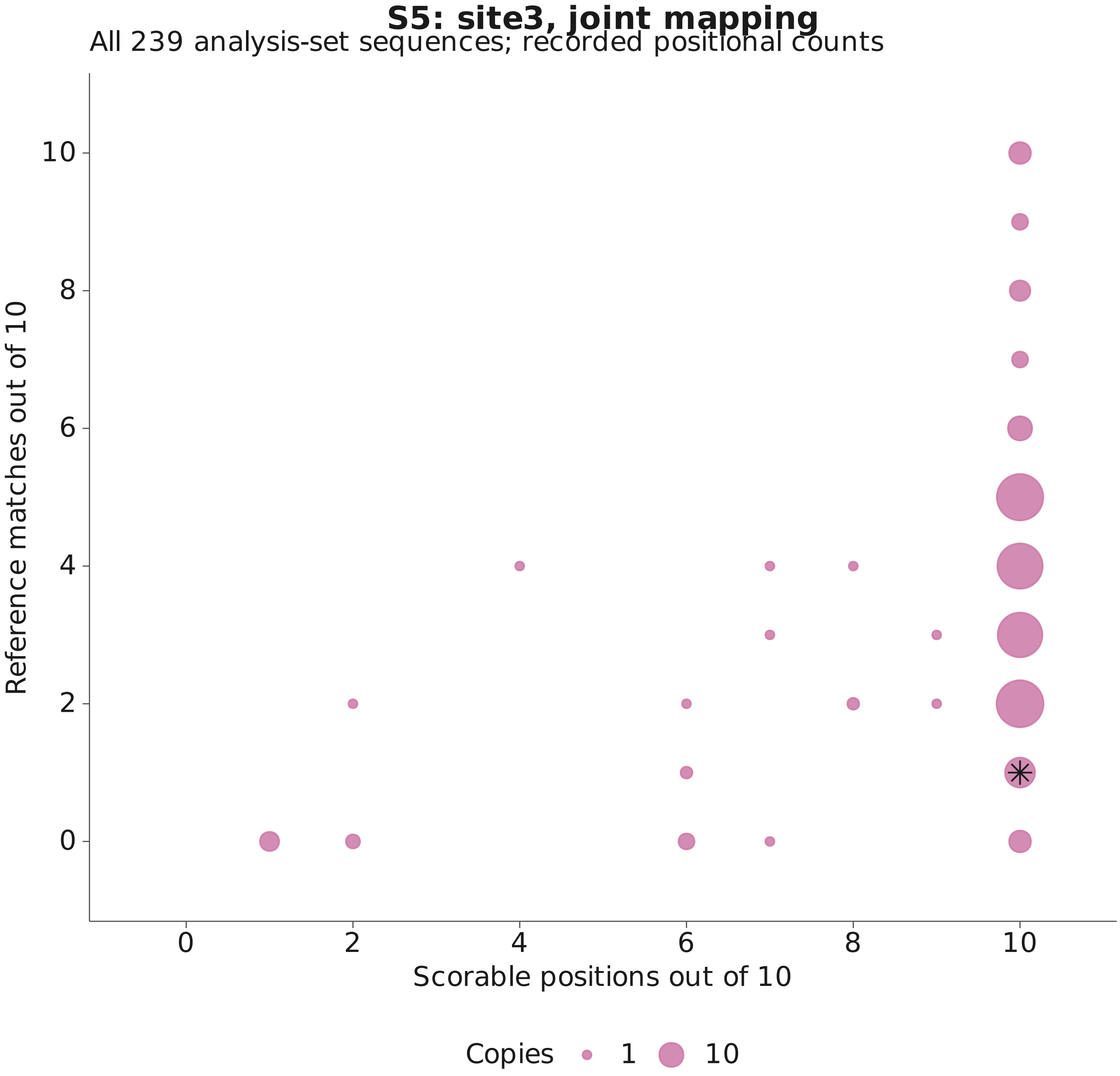


Point area: number of copies. Black star: Toxoplasma gondii.
Zero scorable positions indicates mapping coverage, not loss.
Position sets and alignment procedures remain separate.

### S5: site4, joint mapping

All 239 analysis-set sequences; recorded positional counts

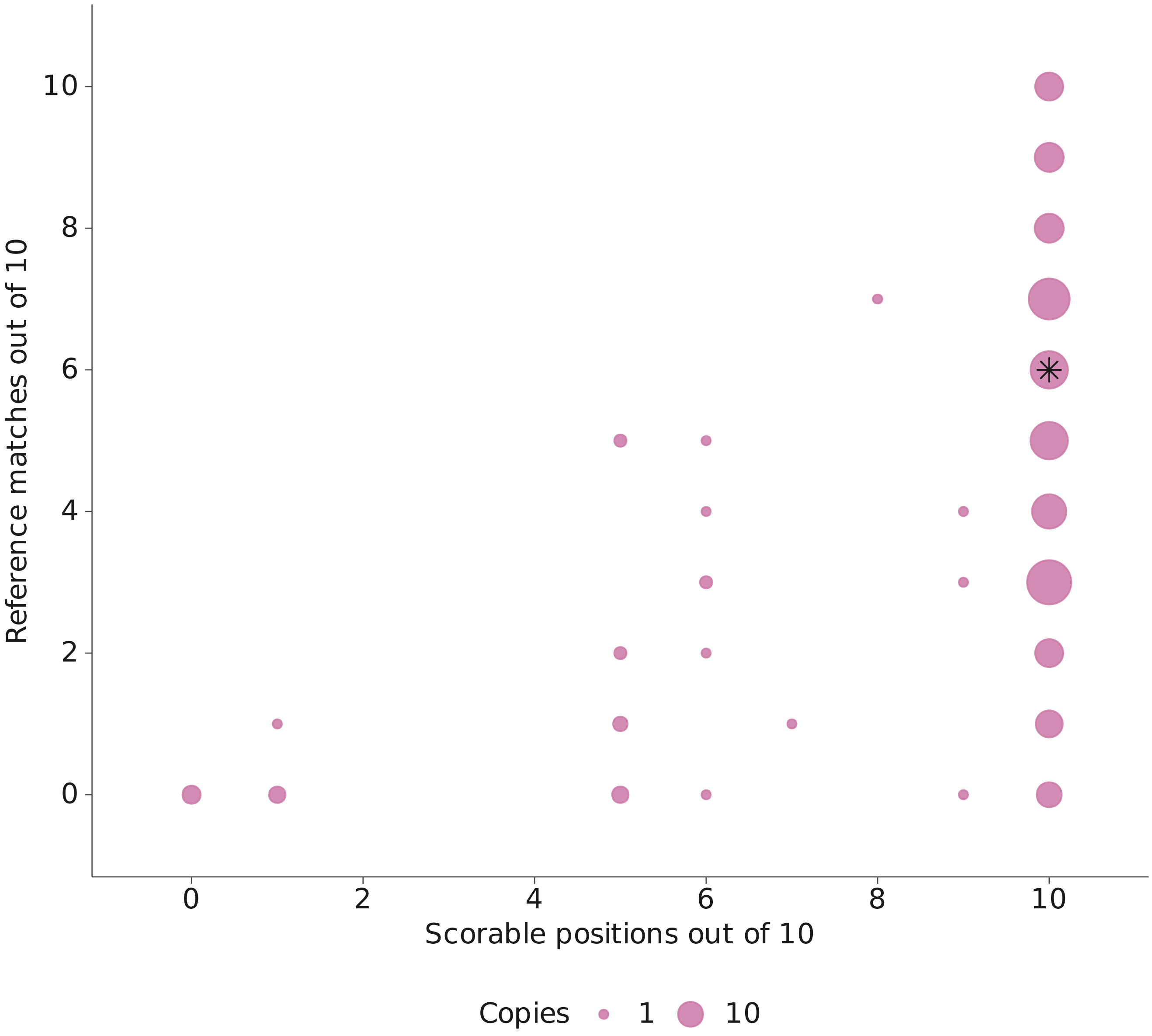


Point area: number of copies. Black star: Toxoplasma gondii.
Zero scorable positions indicates mapping coverage, not loss.
Position sets and alignment procedures remain separate.
