## Supplementary material for "Divergence of TOR and AMPK regulators across 210 eukaryotes predicts a dispensable TOR in Toxoplasma gondii": S6

### S6A: site 1 alignment comparison

239 analysis-set sequences; frozen complete-set scores

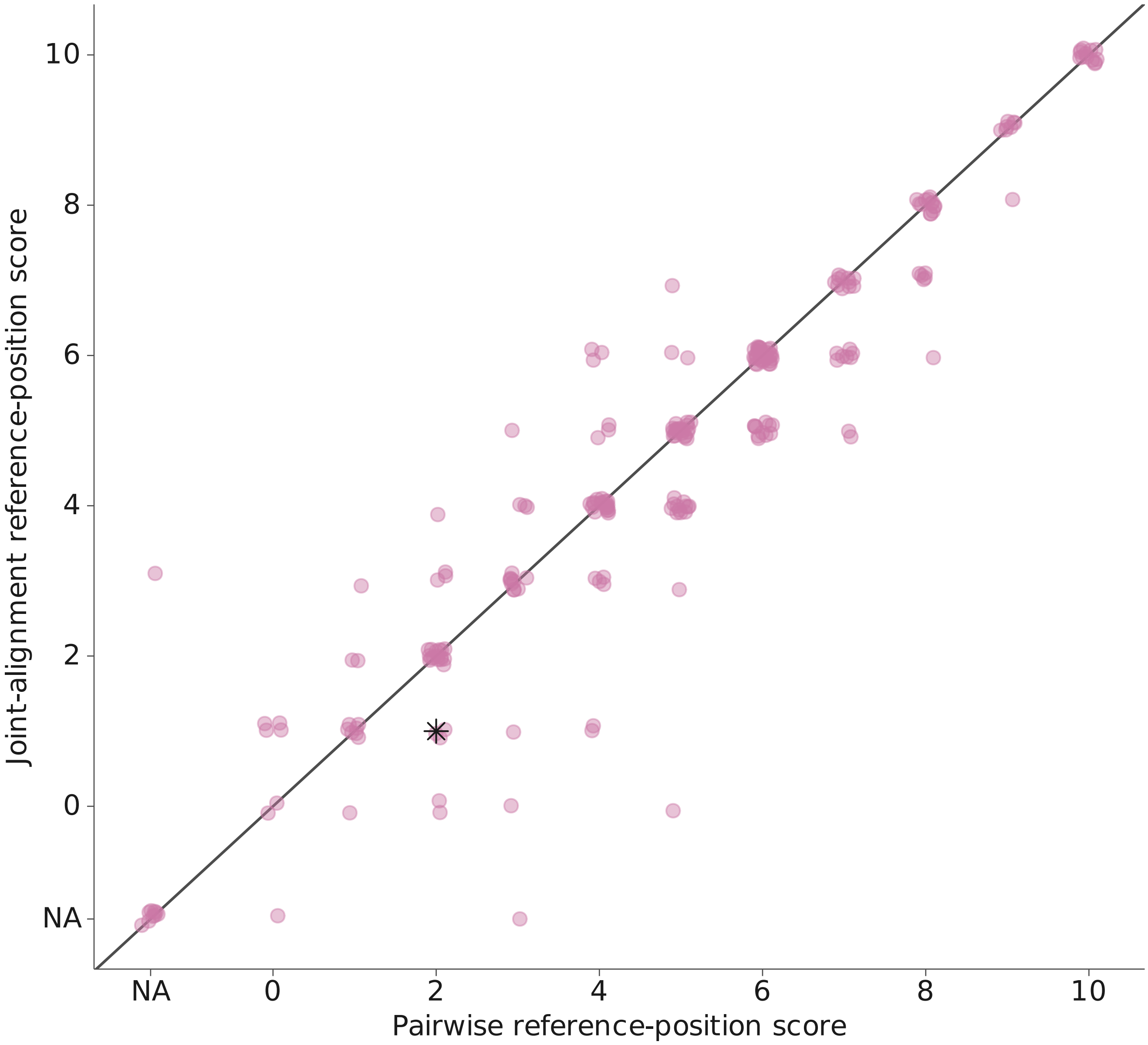


Small fixed jitter reveals overlapping points; source values are unchanged. Black star: Toxoplasma gondii. Unavailable scores remain at NA, separate from observed zero scores.

### S6A: site 3 alignment comparison

239 analysis-set sequences; frozen complete-set scores

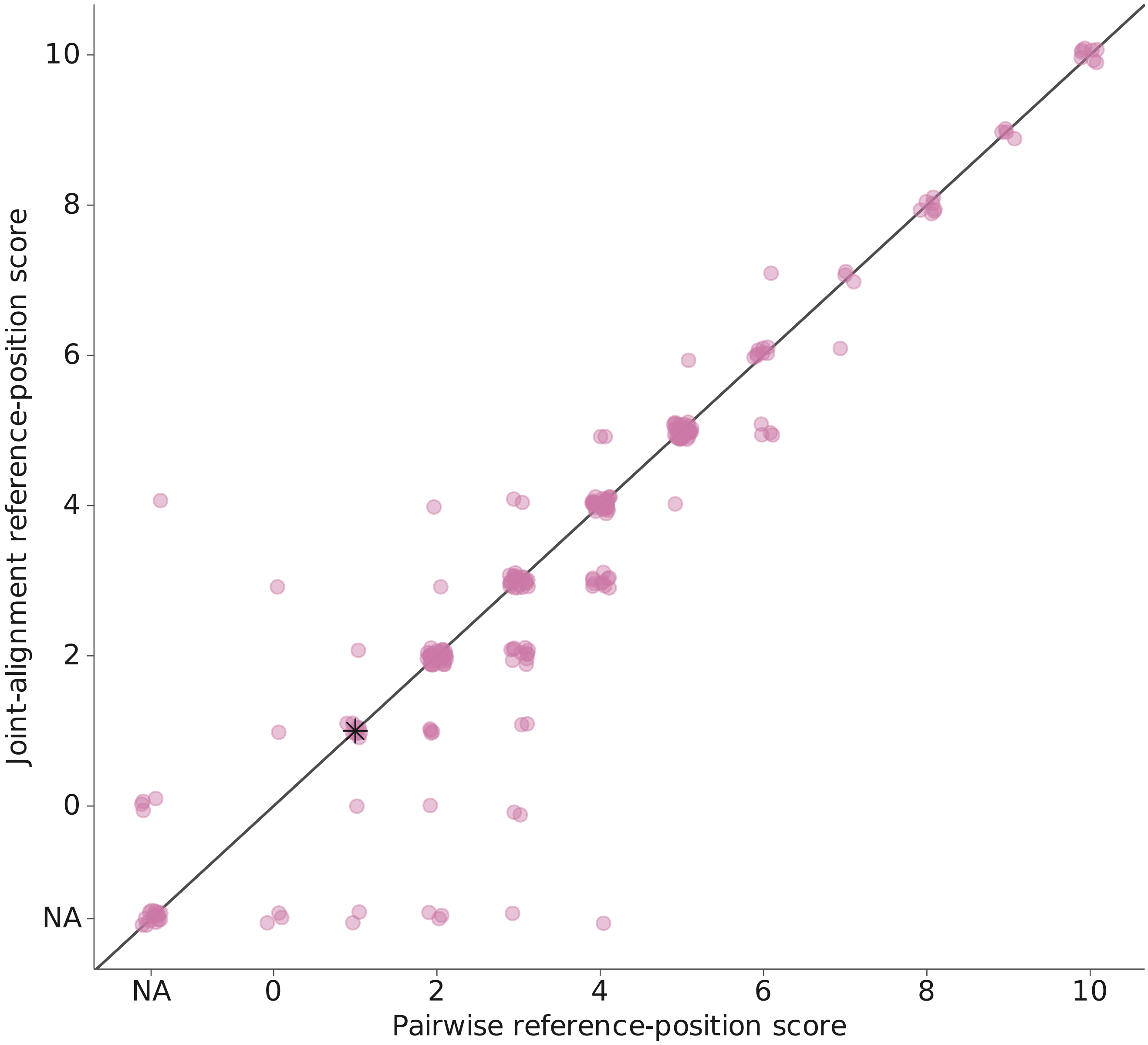


Small fixed jitter reveals overlapping points; source values are unchanged. Black star: Toxoplasma gondii. Unavailable scores remain at NA, separate from observed zero scores.

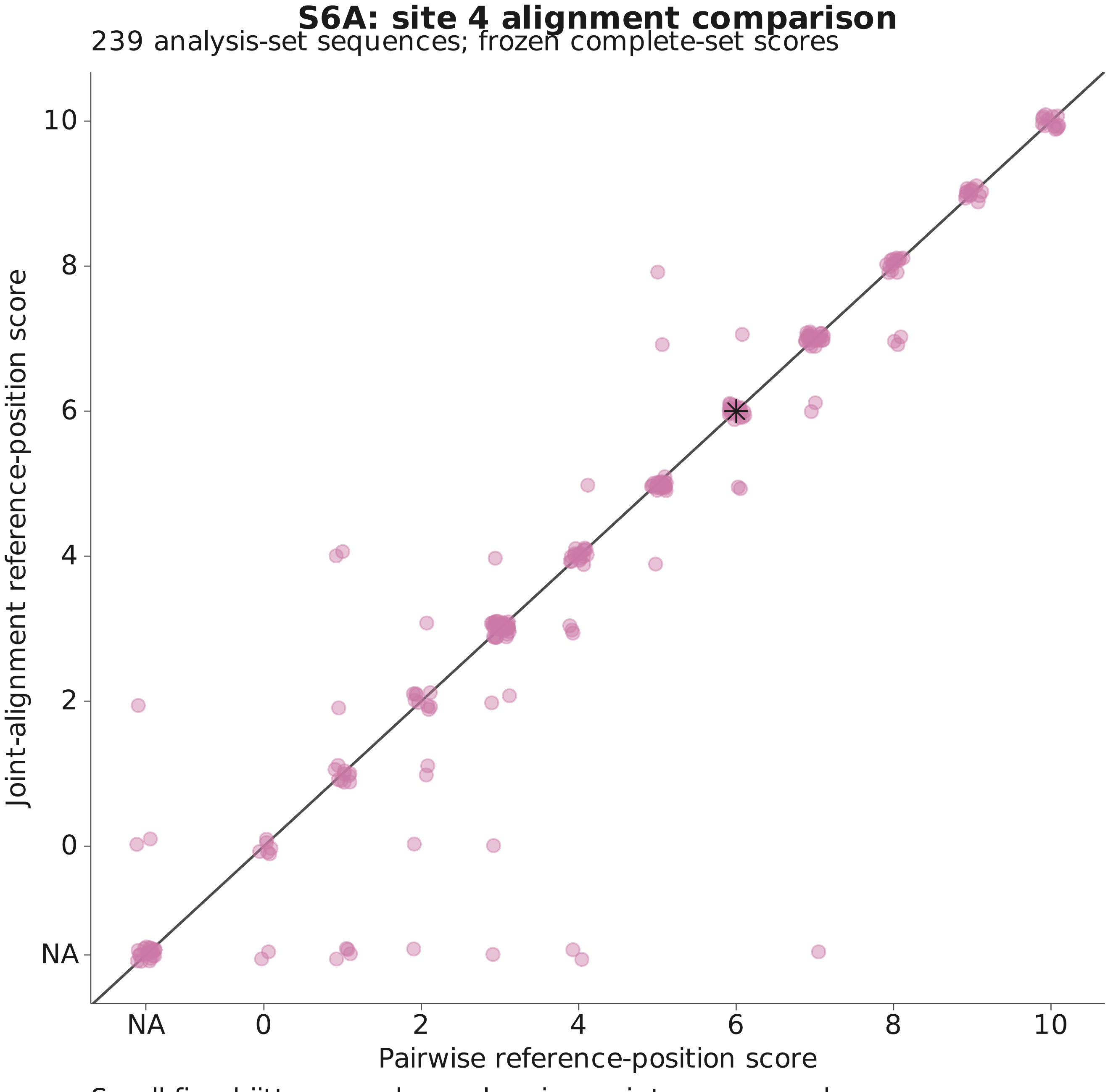


Small fixed jitter reveals overlapping points; source values are unchanged. Black star: Toxoplasma gondii. Unavailable scores remain at NA, separate from observed zero scores.

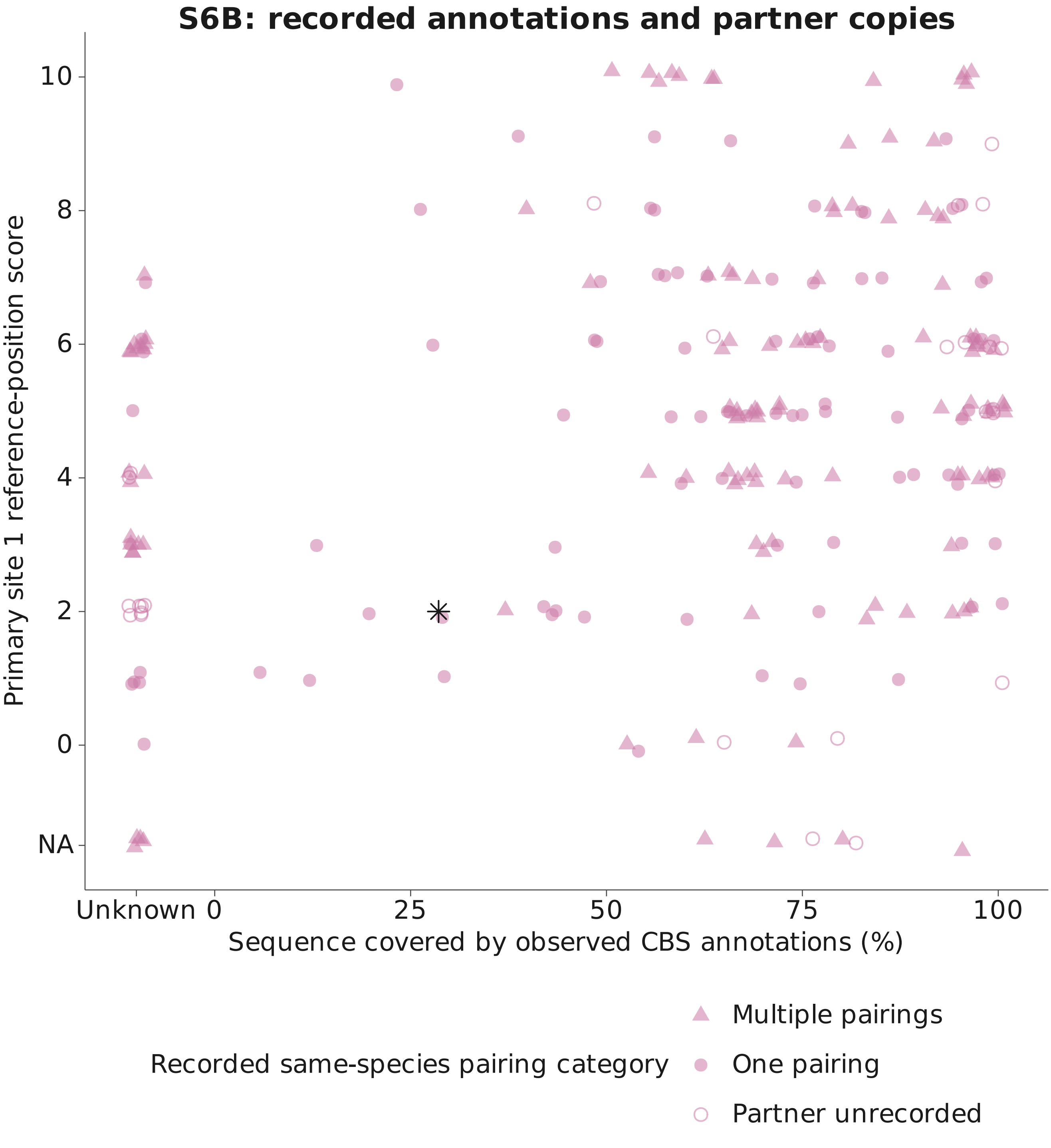


239 copies; annotation intervals retain the source union rule.
Small fixed jitter shows overlap. Black star: Toxoplasma gondii.
Unknown annotations and unrecorded partners are not absences.

#### S6C: length against recorded domain span

239 analysis-set sequences; annotation intervals retain the source union rule

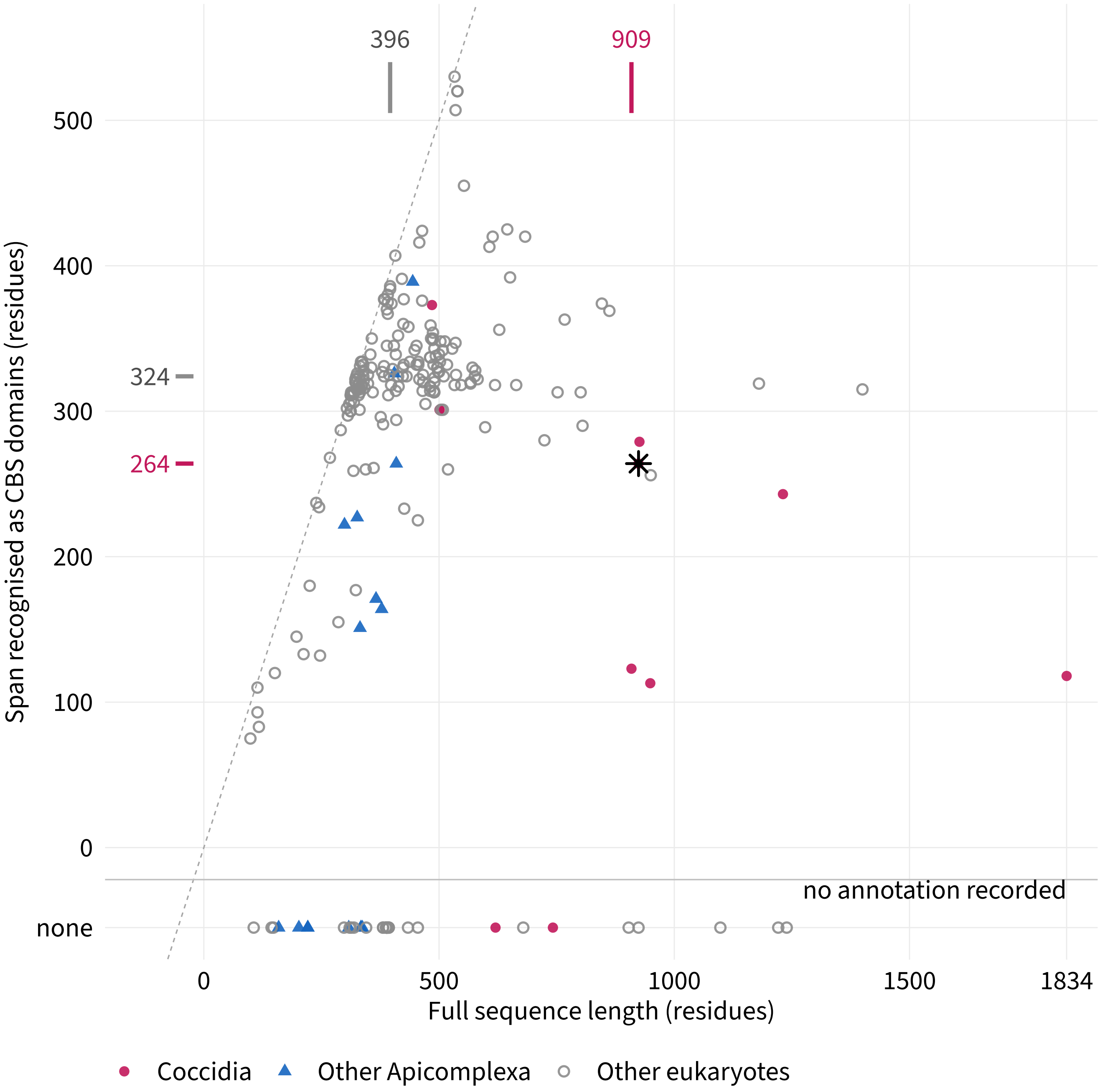


Dashed: the whole sequence recognised. Black star: Toxoplasma gondii.
Ticks outside the axes mark the group medians.
Unknown annotations are shown as none, not as a span of zero.
