## Supplementary material for "Divergence of TOR and AMPK regulators across 210 eukaryotes predicts a dispensable TOR in Toxoplasma gondii": S7

### AMPKgamma structural predictions: AMP1327

239 copies and 5 predicted controls; all queries accounted for

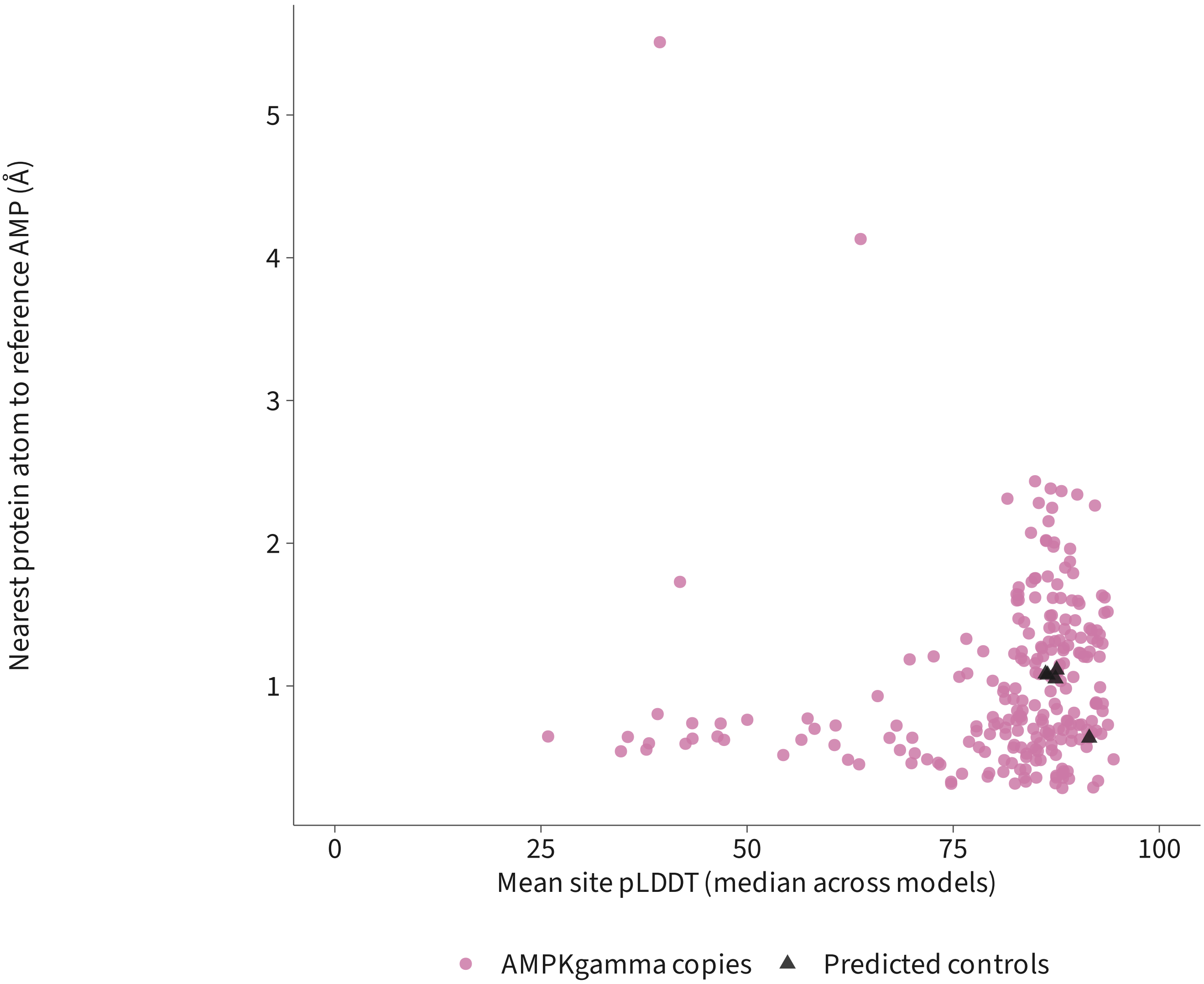


| | Measurement unavailable | Point shown |
|---|---|---|
| Predicted controls | 0 | 5 |
| AMPKgamma copies | 4 | 235 |

One point per query: median of available model measurements.
Counts include unavailable measurements; no confidence cutoff applied.
Computational descriptions, not binding or functional-loss measurements.

### AMPKgamma structural predictions: AMP1327

239 copies and 5 predicted controls; all queries accounted for

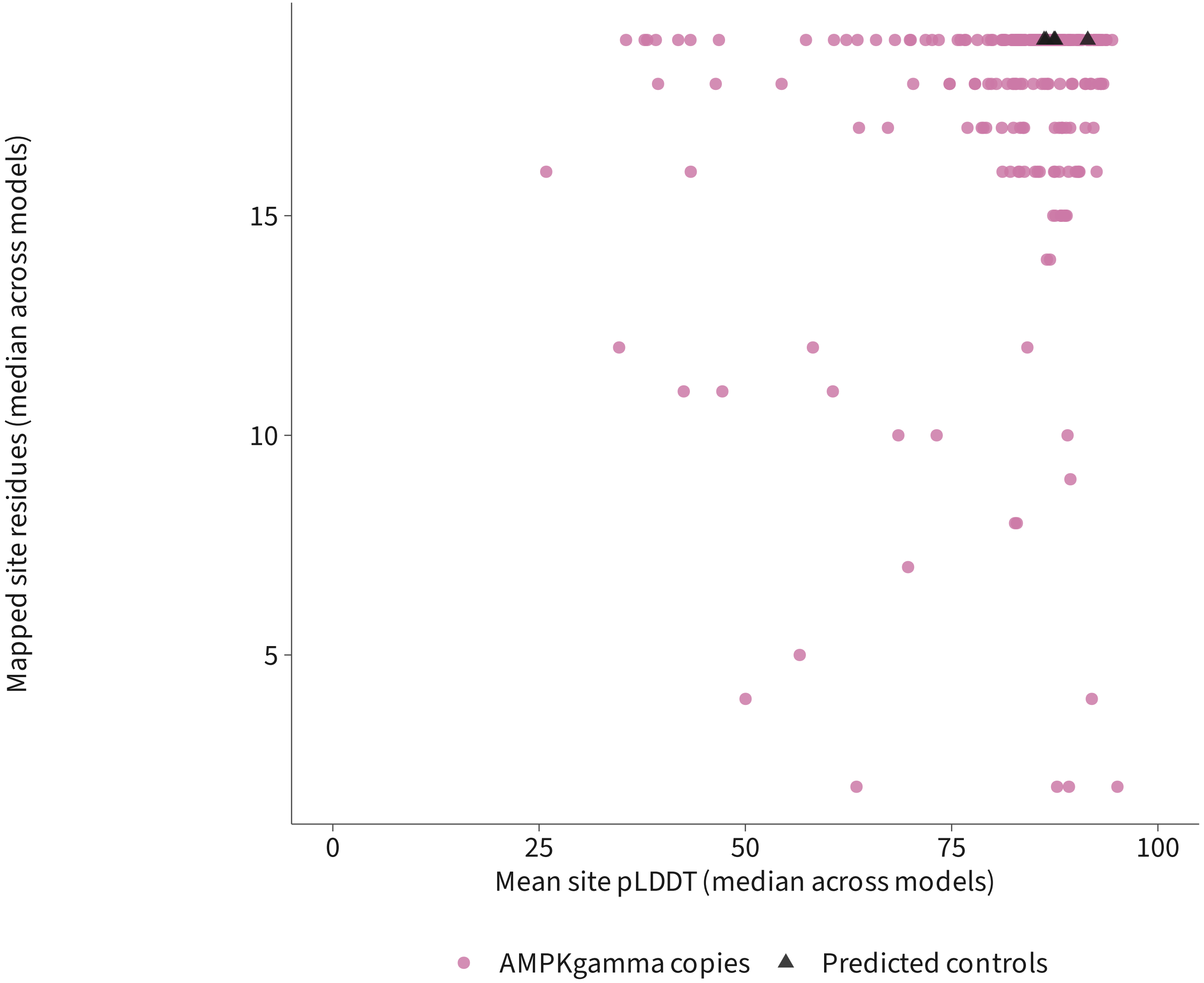


| | Measurement unavailable | Point shown |
|---|---|---|
| Predicted controls | 0 | 5 |
| AMPKgamma copies | 0 | 239 |

One point per query: median of available model measurements.
Counts include unavailable measurements; no confidence cutoff applied.
Computational descriptions, not binding or functional-loss measurements.

### AMPKgamma structural predictions: AMP1327

239 copies and 5 predicted controls; all queries accounted for

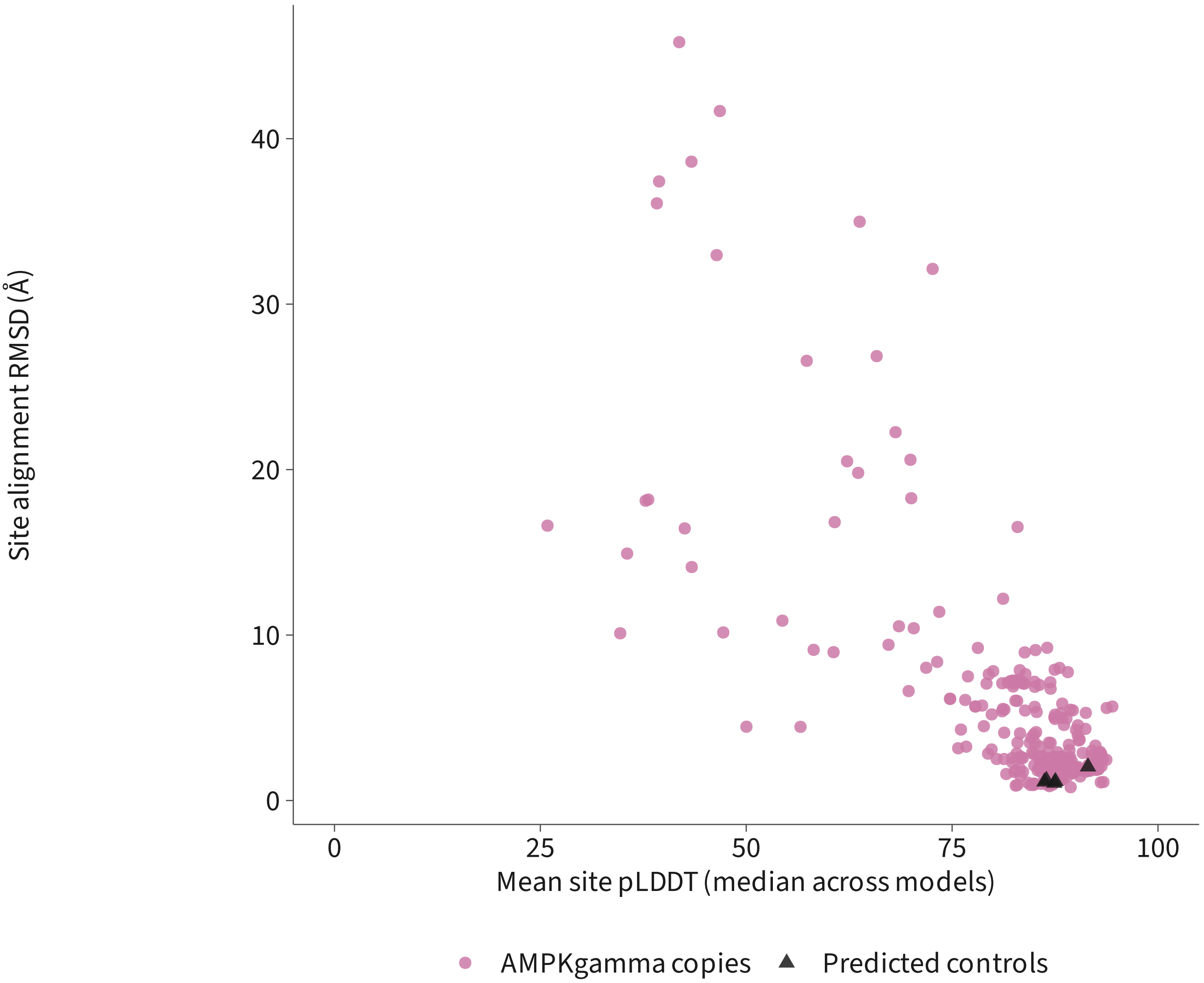


| | Measurement unavailable | Point shown |
|---|---|---|
| Predicted controls | 0 | 5 |
| AMPKgamma copies | 4 | 235 |

One point per query: median of available model measurements.
Counts include unavailable measurements; no confidence cutoff applied.
Computational descriptions, not binding or functional-loss measurements.

### AMPKgamma structural predictions: AMP1328

239 copies and 5 predicted controls; all queries accounted for

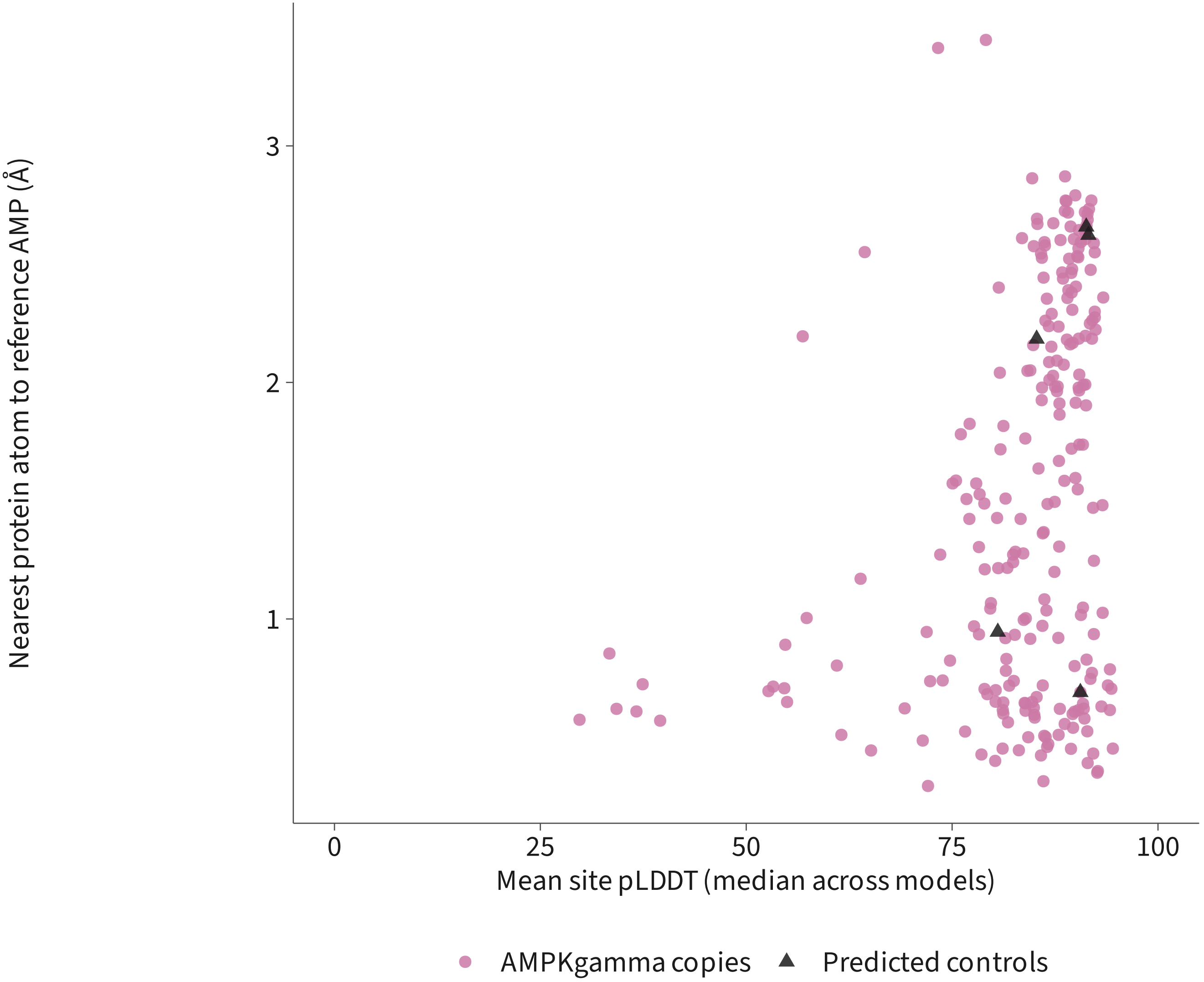


| | Measurement unavailable | Point shown |
|---|---|---|
| Predicted controls | 0 | 5 |
| AMPKgamma copies | 4 | 235 |

One point per query: median of available model measurements.
Counts include unavailable measurements; no confidence cutoff applied.
Computational descriptions, not binding or functional-loss measurements.

### AMPKgamma structural predictions: AMP1328

239 copies and 5 predicted controls; all queries accounted for

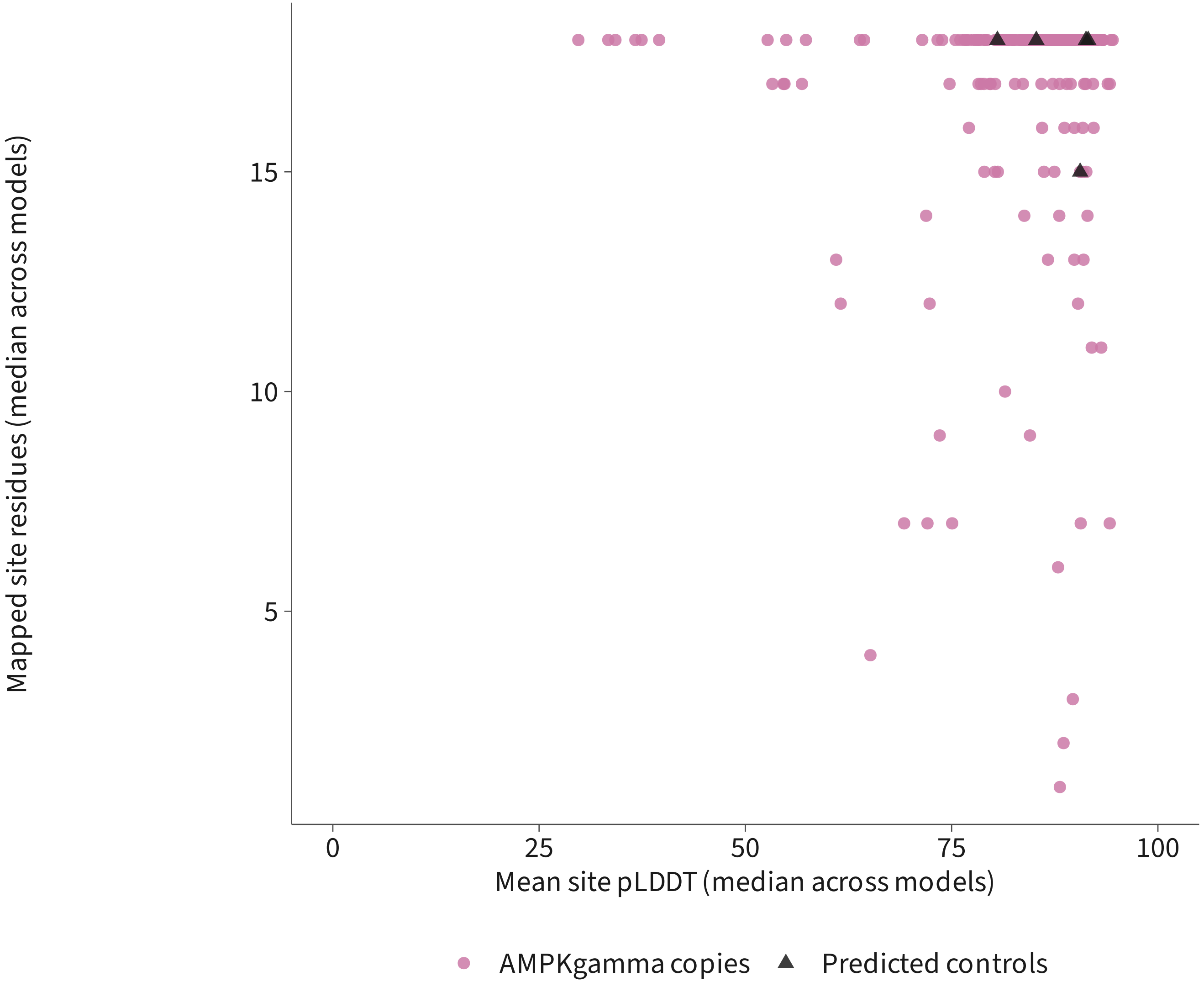


| | Measurement unavailable | Point shown |
|---|---|---|
| Predicted controls | 0 | 5 |
| AMPKgamma copies | 2 | 237 |

One point per query: median of available model measurements.
Counts include unavailable measurements; no confidence cutoff applied.
Computational descriptions, not binding or functional-loss measurements.

### AMPKgamma structural predictions: AMP1328

239 copies and 5 predicted controls; all queries accounted for

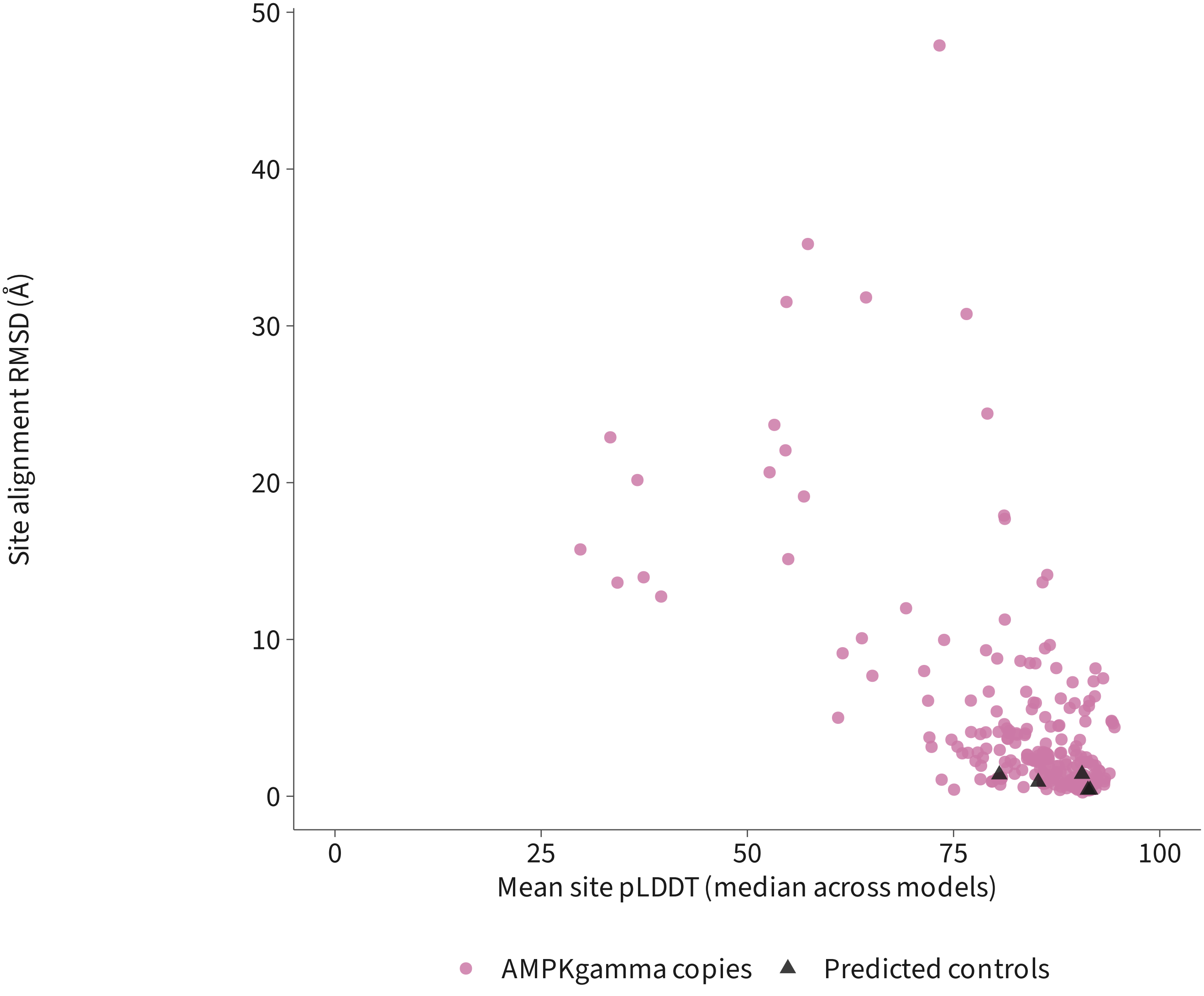


| | Measurement unavailable | Point shown |
|---|---|---|
| Predicted controls | 0 | 5 |
| AMPKgamma copies | 4 | 235 |

One point per query: median of available model measurements.
Counts include unavailable measurements; no confidence cutoff applied.
Computational descriptions, not binding or functional-loss measurements.

### AMPKgamma structural predictions: AMP1329

239 copies and 5 predicted controls; all queries accounted for

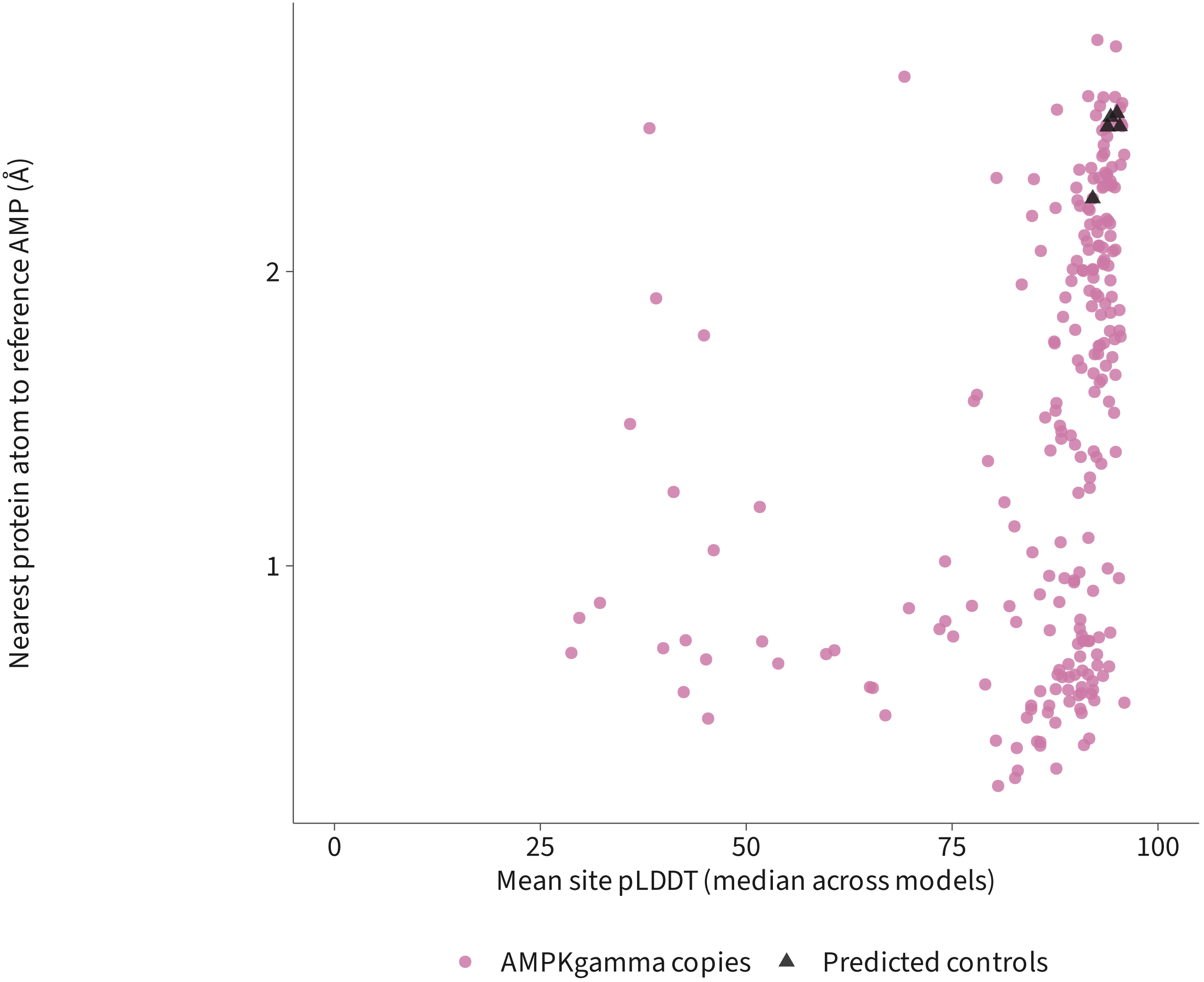


| | Measurement unavailable | Point shown |
|---|---|---|
| Predicted controls | 0 | 5 |
| AMPKgamma copies | 4 | 235 |

One point per query: median of available model measurements.
Counts include unavailable measurements; no confidence cutoff applied.
Computational descriptions, not binding or functional-loss measurements.

### AMPKgamma structural predictions: AMP1329

239 copies and 5 predicted controls; all queries accounted for

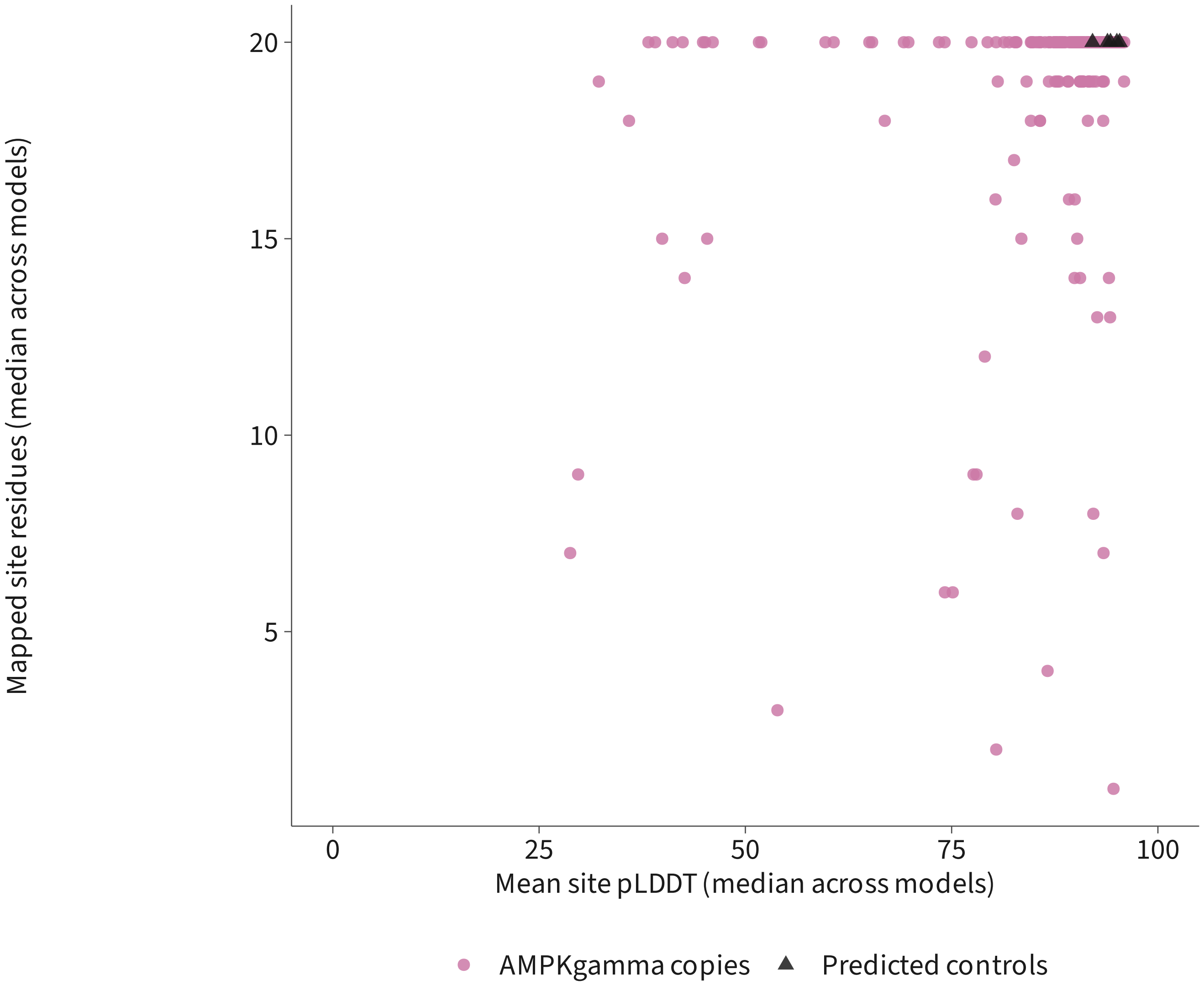


| | Measurement unavailable | Point shown |
|---|---|---|
| Predicted controls | 0 | 5 |
| AMPKgamma copies | 2 | 237 |

One point per query: median of available model measurements.
Counts include unavailable measurements; no confidence cutoff applied.
Computational descriptions, not binding or functional-loss measurements.

### AMPKgamma structural predictions: AMP1329

239 copies and 5 predicted controls; all queries accounted for

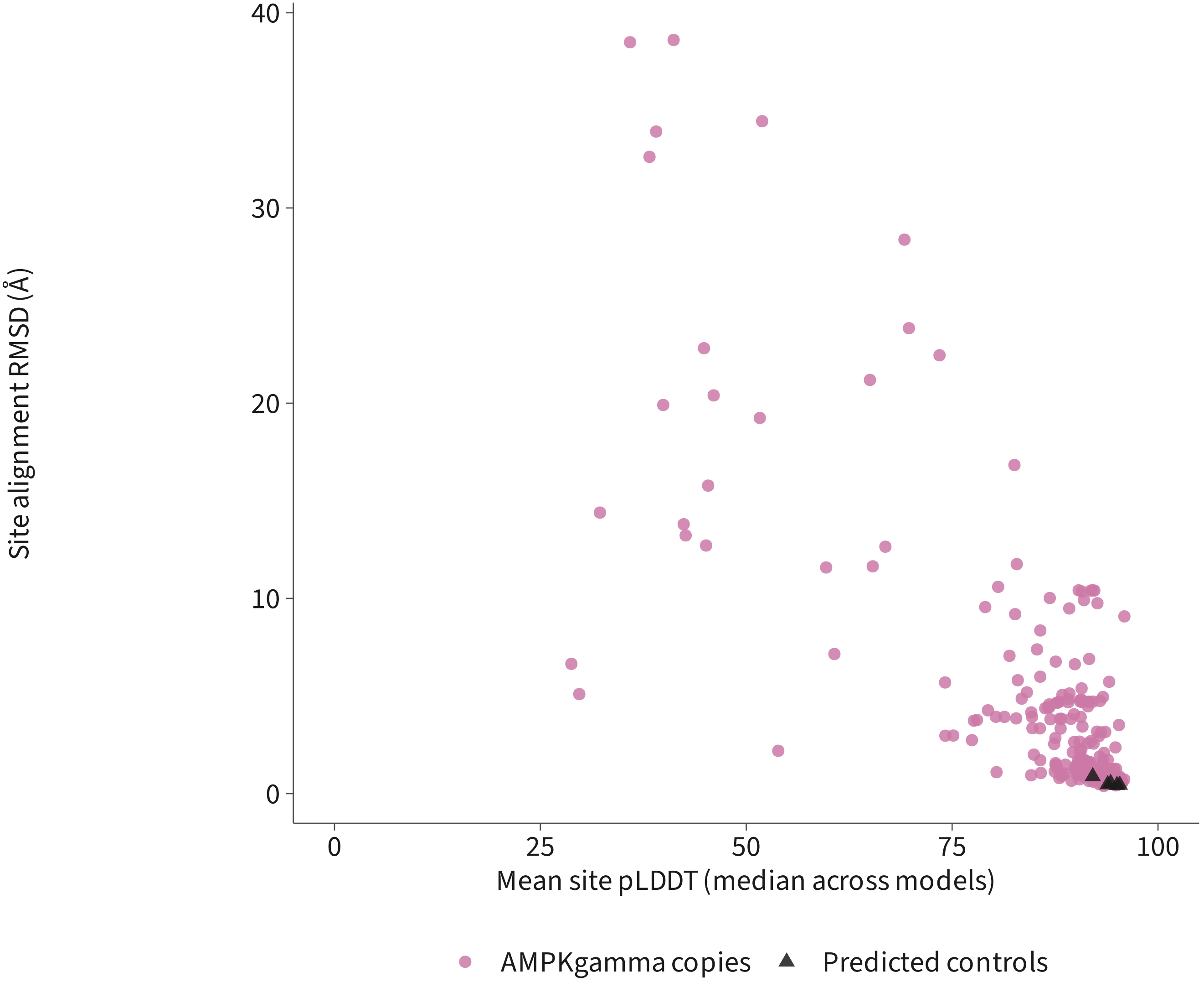


| | Measurement unavailable | Point shown |
|---|---|---|
| Predicted controls | 0 | 5 |
| AMPKgamma copies | 4 | 235 |

One point per query: median of available model measurements.
Counts include unavailable measurements; no confidence cutoff applied.
Computational descriptions, not binding or functional-loss measurements.
