## Supplementary material for "Divergence of TOR and AMPK regulators across 210 eukaryotes predicts a dispensable TOR in Toxoplasma gondii": S8

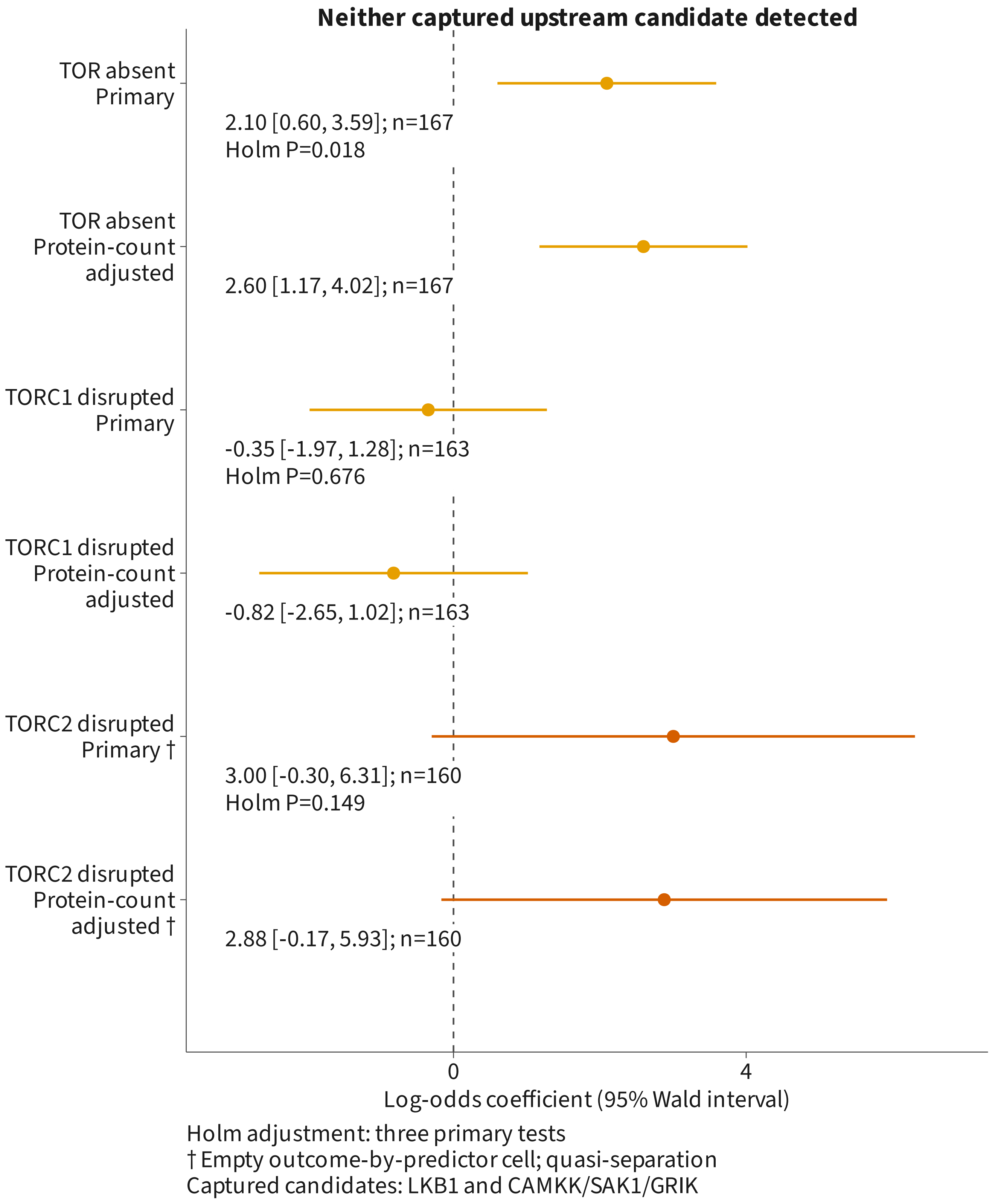
Neither captured upstream candidate detected
TOR absent
Primary
2.10 [0.60, 3.59]; n=167
Holm P=0.018
TOR absent
Protein-count
adjusted
2.60 [1.17, 4.02]; n=167
TORC1 disrupted
Primary
-0.35 [-1.97, 1.28]; n=163
Holm P=0.676
TORC1 disrupted
Protein-count
adjusted
-0.82 [-2.65, 1.02]; n=163
TORC2 disrupted
Primary †
3.00 [-0.30, 6.31]; n=160
Holm P=0.149
TORC2 disrupted
Protein-count
adjusted †
2.88 [-0.17, 5.93]; n=160
0
4
Log-odds coefficient (95% Wald interval)
Holm adjustment: three primary tests
† Empty outcome-by-predictor cell; quasi-separation
Captured candidates: LKB1 and CAMKK/SAK1/GRIK
