## Supplementary material for "Divergence of TOR and AMPK regulators across 210 eukaryotes predicts a dispensable TOR in Toxoplasma gondii": S9

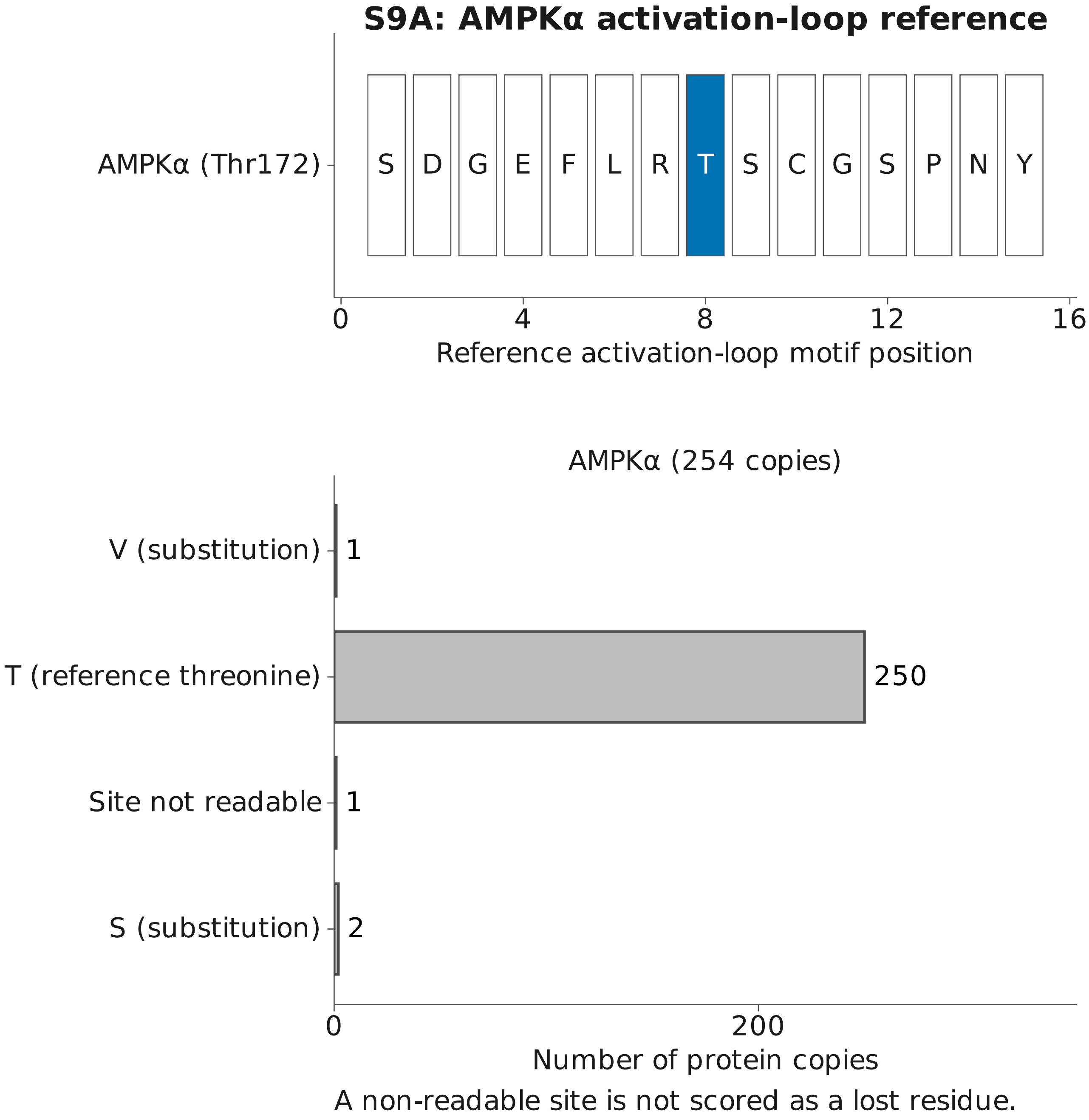
S9A: AMPKα activation-loop reference
AMPKα (Thr172)
S D G E F L R T S C G S P N Y
0
4
8
12
16
Reference activation-loop motif position
AMPKα (254 copies)
V (substitution)
1
T (reference threonine)
250
Site not readable
1
S (substitution)
2
0
200
Number of protein copies
A non-readable site is not scored as a lost residue.

#### S9B: activation-loop sequence identity

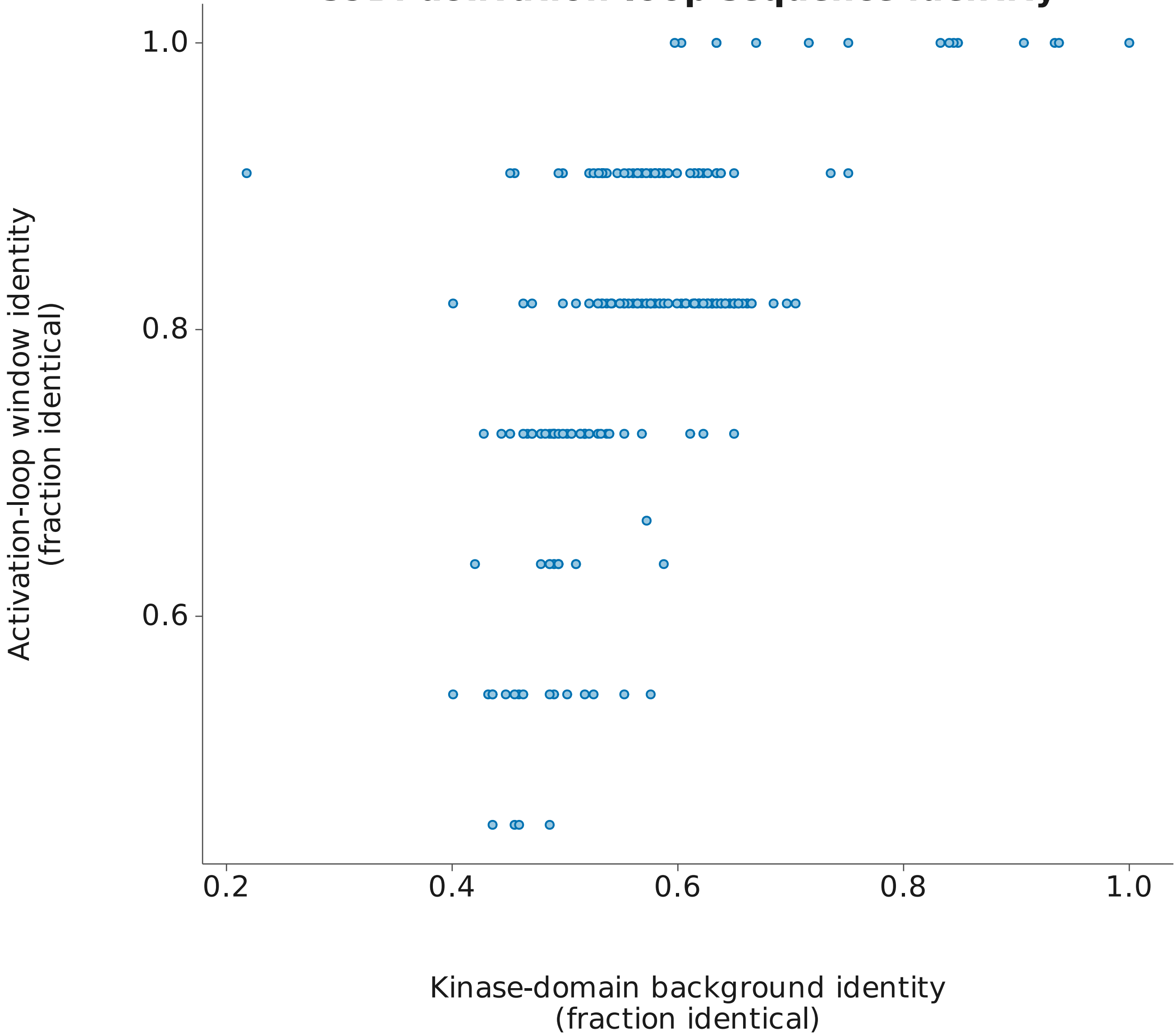


One point per scored protein copy. No regression is fitted.

**S9C: site calls and upstream kinase detection**

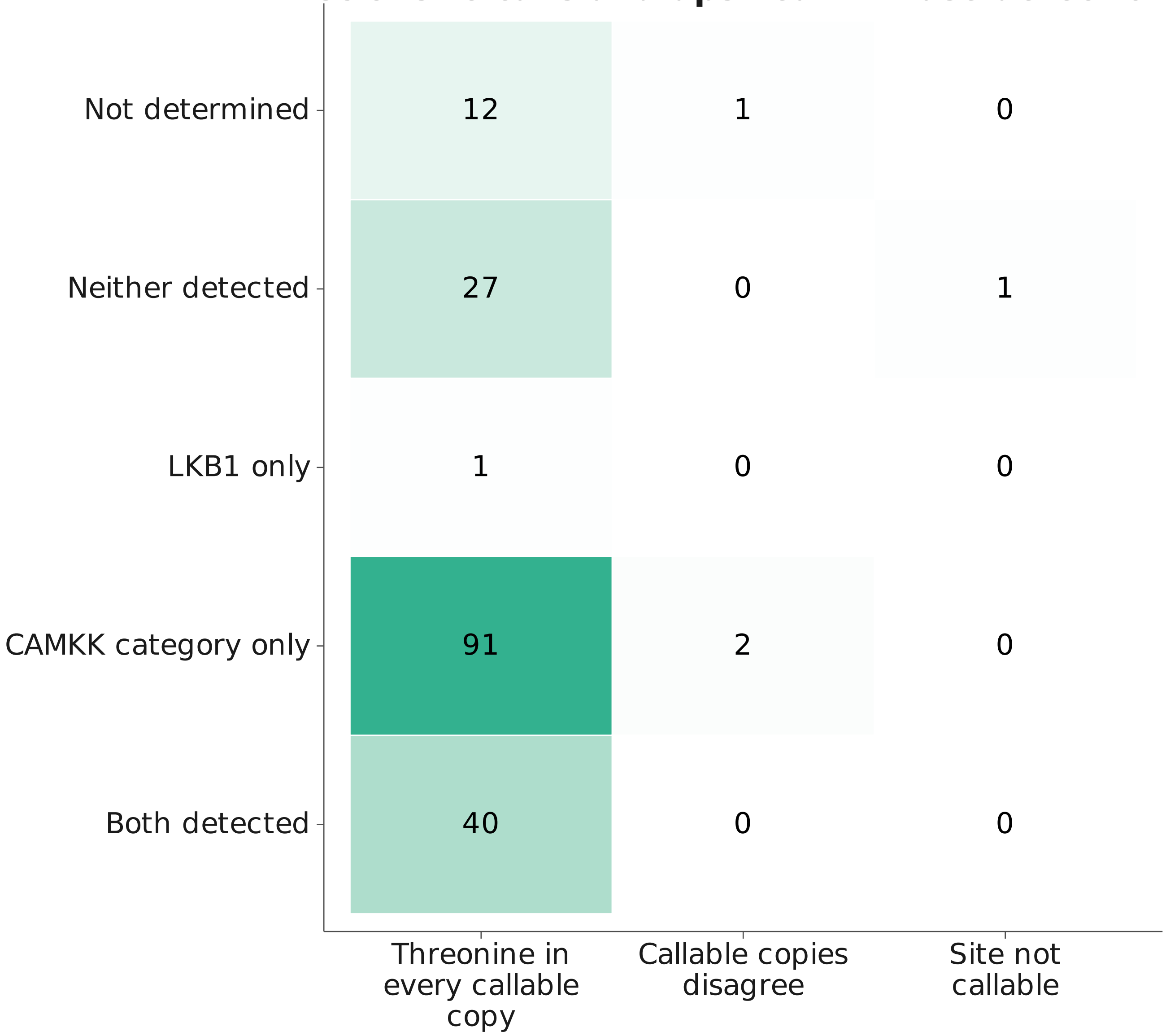


Numbers are species counts, including zero observations.
Neither detected refers only to the two searched upstream kinase categories.

### S9D: activation-loop regression results

Site not a threonine; No covariate; Phylogenetic logistic
n=171 species; Not estimable (retained 171, not retained 0)
No coefficient estimated

Median window identity; No covariate; Phylogenetic linear (lambda)
n=168 species; Fitted
Coefficient -0.0041; 95% Wald interval [-0.0179, 0.0098]

Median window identity; Covariate: Median background identity; Phylogenetic linear (lambda)
n=168 species; Fitted
Coefficient -0.0113; 95% Wald interval [-0.0281, 0.0054]

Median window identity; No covariate; Phylogenetic linear (Brownian)
n=168 species; Fitted
Coefficient -0.0041; 95% Wald interval [-0.0179, 0.0098]

Predictor: neither searched upstream kinase category detected.
Intervals are nominal and unadjusted for multiple comparisons.

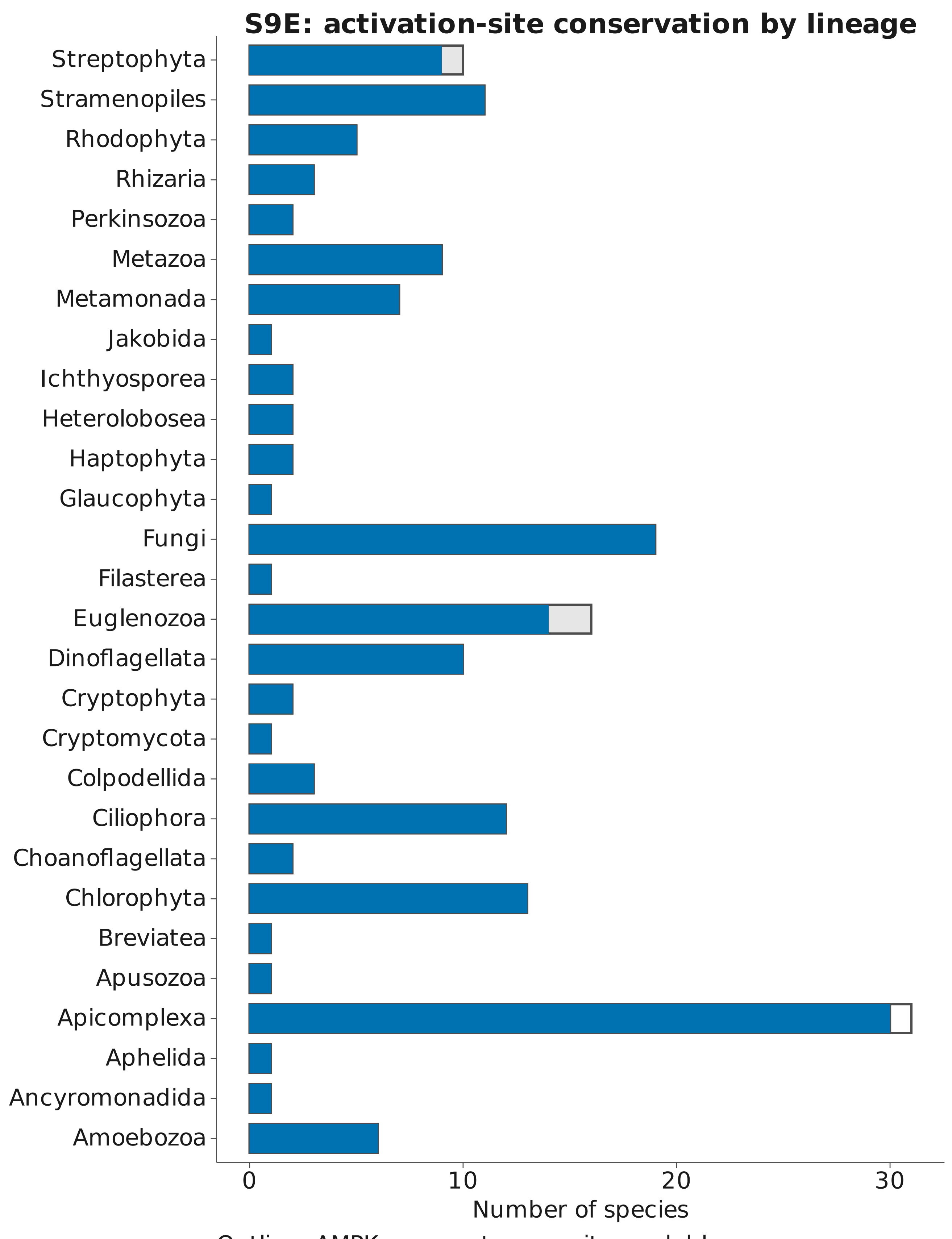


Outline: AMPKα present; grey: site readable;
solid color: threonine in every readable copy.
