## Supplementary material for "Divergence of TOR and AMPK regulators across 210 eukaryotes predicts a dispensable TOR in Toxoplasma gondii": S10

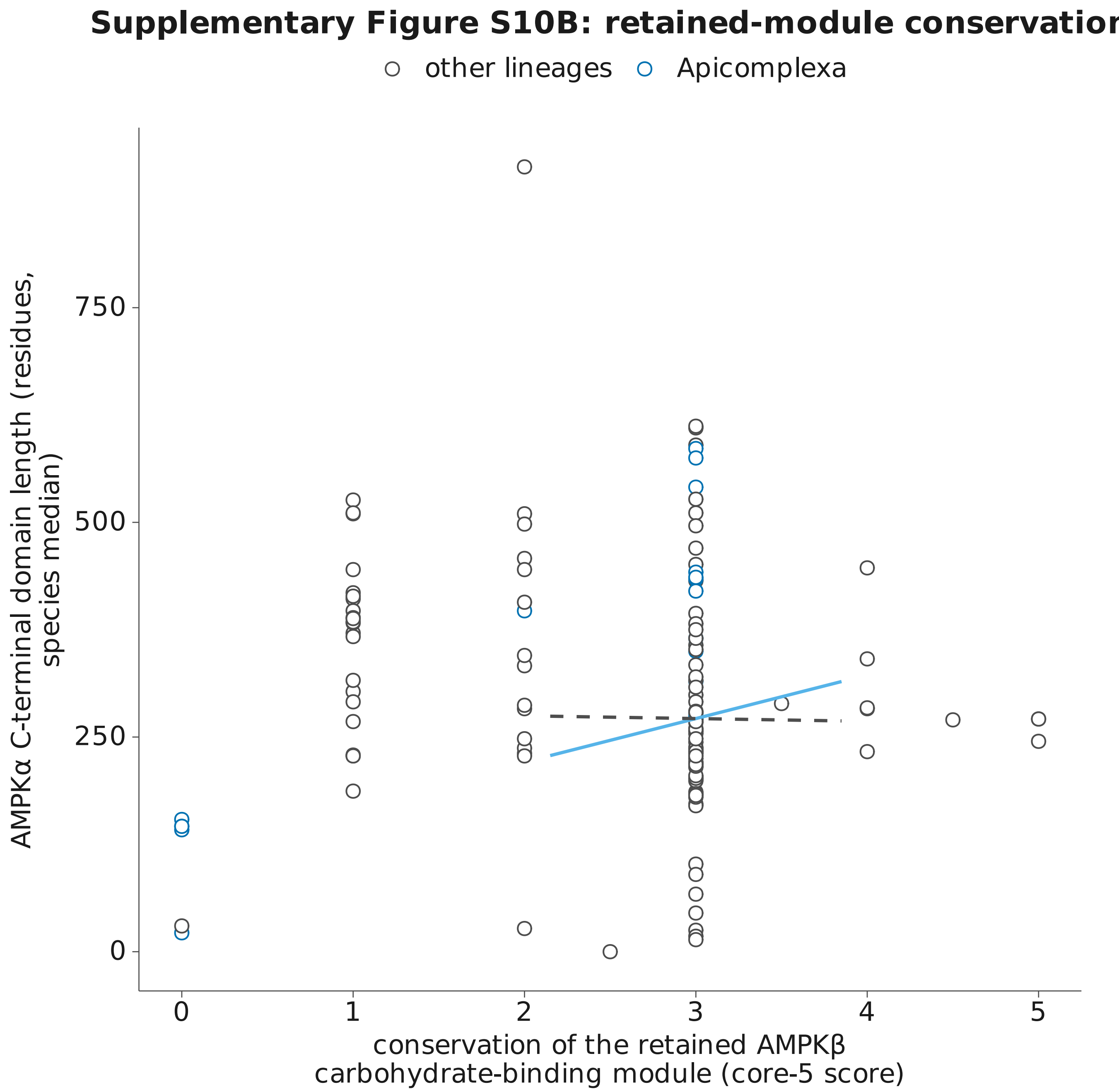


Solid: with phylogenetic correction: 50.7 residues per point, adjusted p 0.00067
Dashed: without: -3.2 residues per point, p 0.78
Segments show model slopes anchored at the species medians; they are not fitted lines through these plotted points.

**G Model-sensitivity status by endpoint family**

One cell is one endpoint family under one kind of sensitivity check.

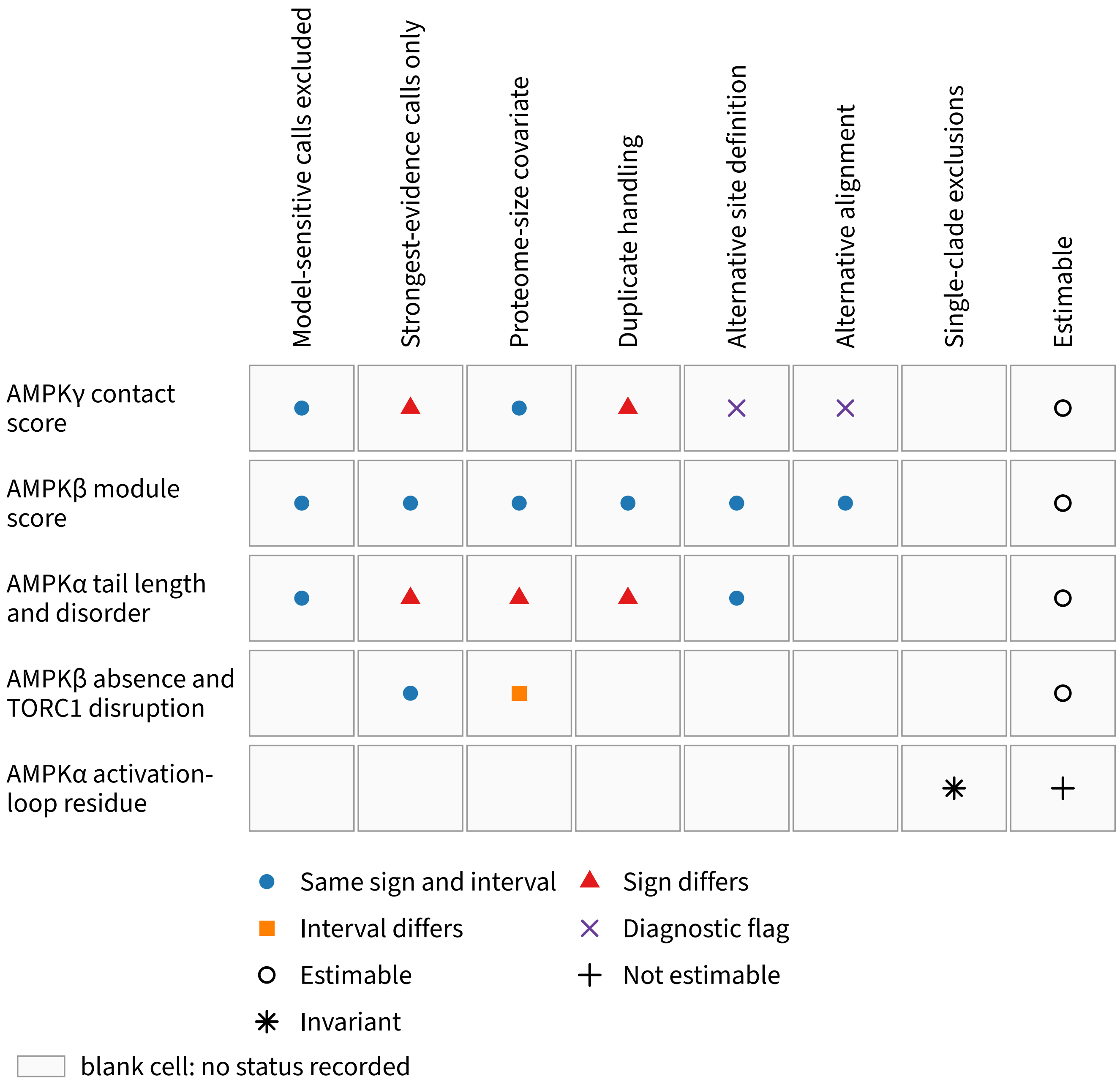


Supplementary Figure [redacted] odel-sensitivity status matrix
