## Supplementary material for "Divergence of TOR and AMPK regulators across 210 eukaryotes predicts a dispensable TOR in Toxoplasma gondii": S11

### S11: recorded absence associations, page 1

Predictor: recorded component absence (rows); Response: recorded component absence (columns)

| | TOR | RAPTOR | RICTOR | LST8 | SIN1 |
|---|---|---|---|---|---|
| TOR | | 0.33 | 2.18 | 1.19 | 0.71 |
| RAPTOR | 0.90 | | 0.09 | 1.05 | 0.57 |
| RICTOR | 2.85 | 0.15 | | 0.72 | 3.75 |
| LST8 | 1.52 | 0.79 | 1.29 | | 0.49 |
| SIN1 | 1.94 | 0.36 | 4.19* | 1.51 | |
| AMPKα | 0.67 | 1.16 | 1.97 | 1.65 | 1.68 |
| AMPKβ | 0.00 | 0.89 | 0.59 | 1.17 | 0.77 |
| AMPKγ | 0.00 | 0.29 | 1.37 | 0.64 | 0.62 |
| LKB1 | 1.13 | 0.00 | 0.07 | 1.13 | 0.00 |
| CAMKK | 0.89 | 0.65 | 0.97 | 0.95 | 3.42 |

* Holm P < 0.05 across all 90 directed primary comparisons.
Dagger: alpha boundary warning. Grey: untested self-comparison.
Values: log-odds coefficients; no causal direction is established.

### S11: recorded absence associations, page 2

| Predictor: recorded component absence | AMPKα | AMPKβ | AMPKγ | LKB1 | CAMKK |
|---|---|---|---|---|---|
| TOR | 0.56 | 0.98 | 1.05 | 0.00 | 1.66 |
| RAPTOR | 0.84 | 3.13 | 0.83 | 0.00 | 1.18 |
| RICTOR | 3.41 | 2.83 | 0.81 | 0.00 | 1.40 |
| LST8 | 1.44 | 3.23* | 0.92 | 0.47 | 1.60 |
| SIN1 | 3.13 | 2.90 | 0.72 | 0.00 | 4.80 |
| AMPKα | | 4.13* | 2.25* | 1.03 | 1.81 |
| AMPKβ | 4.11*† | | 2.11* | 0.00 | 1.26 |
| AMPKγ | 2.01* | 3.63 | | 0.00 | 1.28 |
| LKB1 | 0.79 | -0.49 | 0.38 | | 2.01 |
| CAMKK | 1.26 | 1.06 | 0.91 | 1.97 | |

Response: recorded component absence

* Holm P < 0.05 across all 90 directed primary comparisons.
Dagger: alpha boundary warning. Grey: untested self-comparison.
Values: log-odds coefficients; no causal direction is established.

### S11: rate dependence between paired absences, page 3

45 unordered pairs of the ten targets; Holm within each of 28 pruned trees

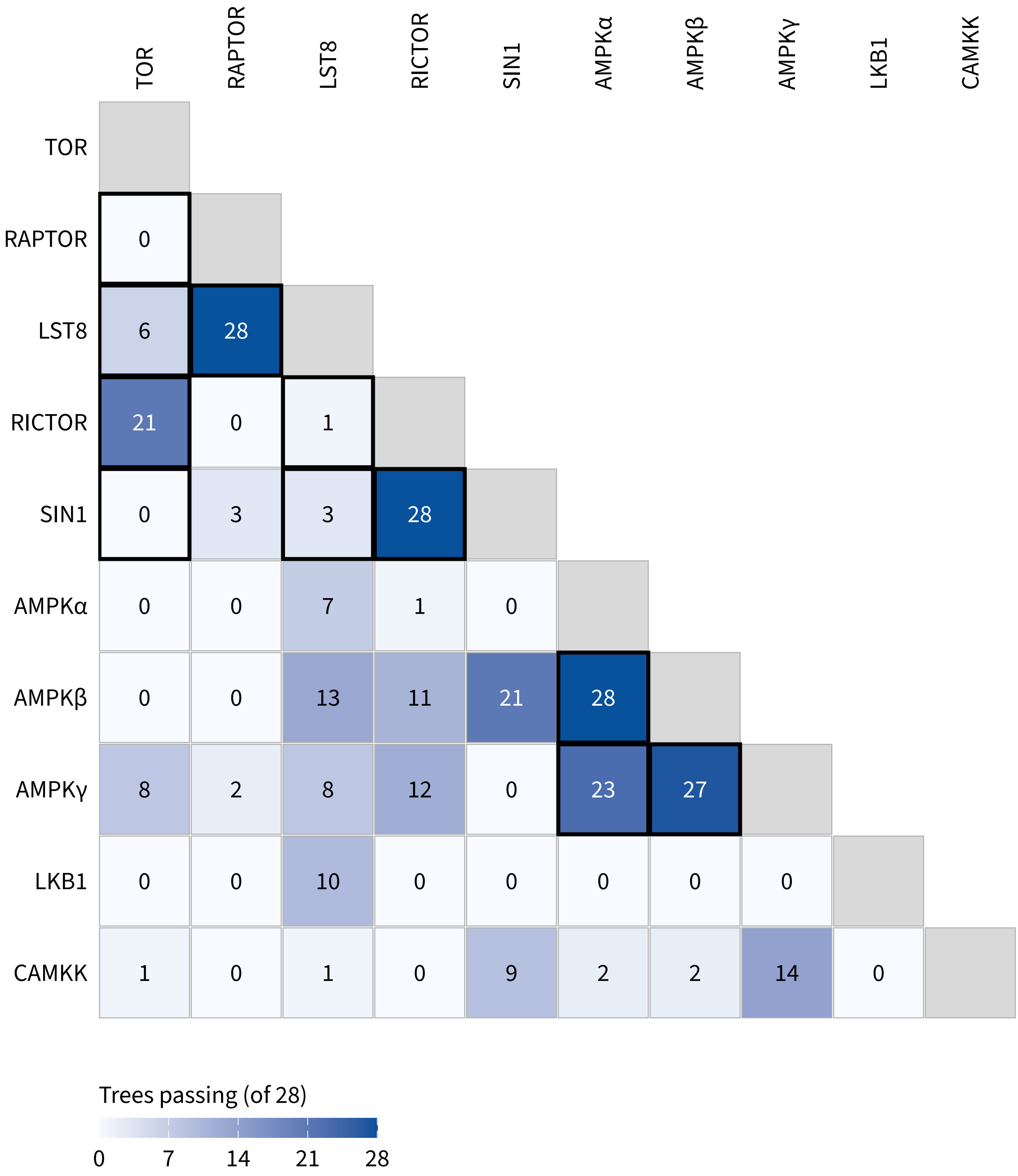


Cells: trees in which the pair passes Holm, of 28.
Heavy border: both proteins in one assembly. Grey: untested self-pair.
Each pair is drawn once; the trees are not independent of one another.
