## Supplementary material for "Divergence of TOR and AMPK regulators across 210 eukaryotes predicts a dispensable TOR in Toxoplasma gondii": S12

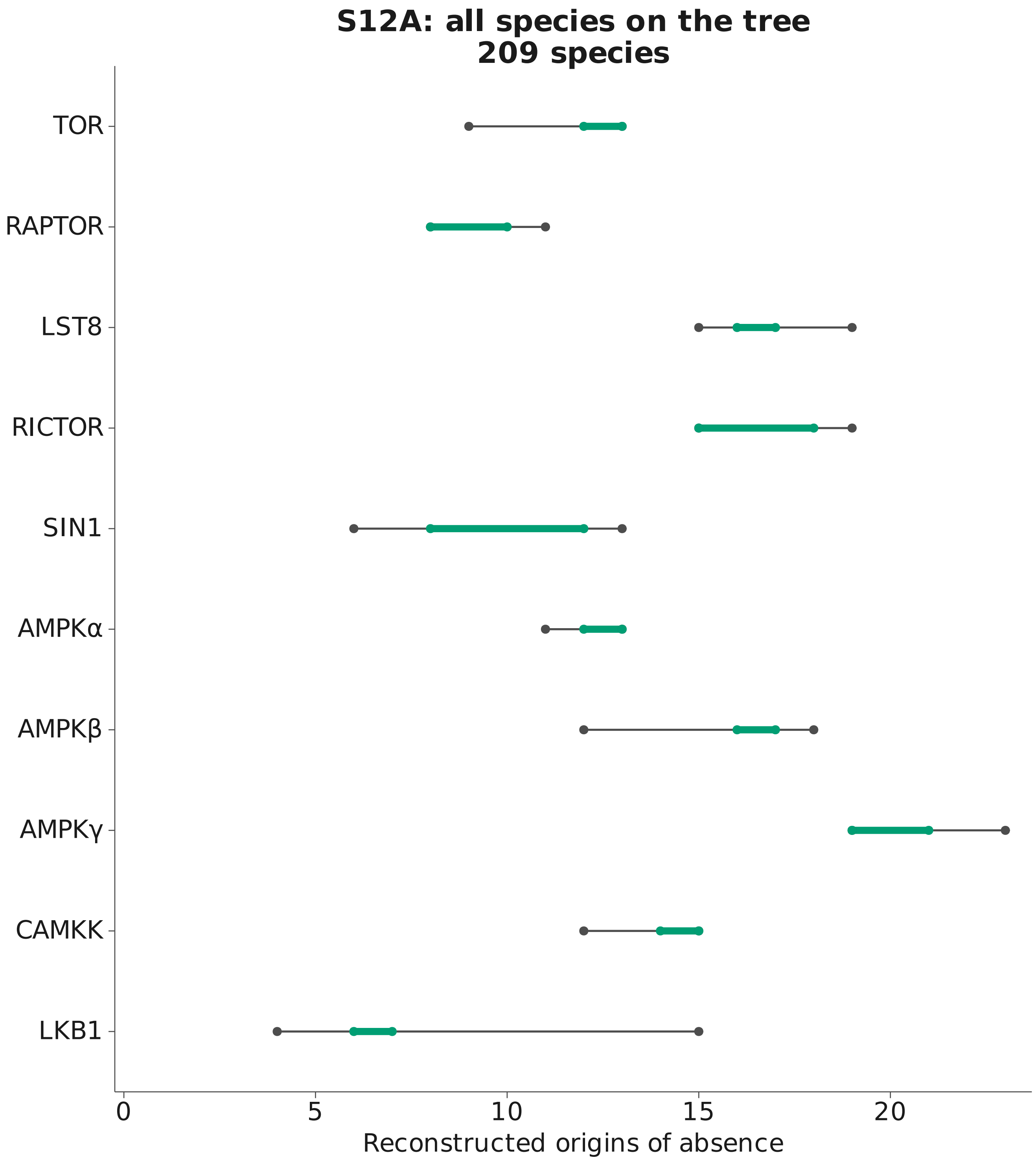


Dark range: all 28 reconstruction conditions
Green range: parsimony conditions only
Ranges describe alternative reconstructions of the same history.

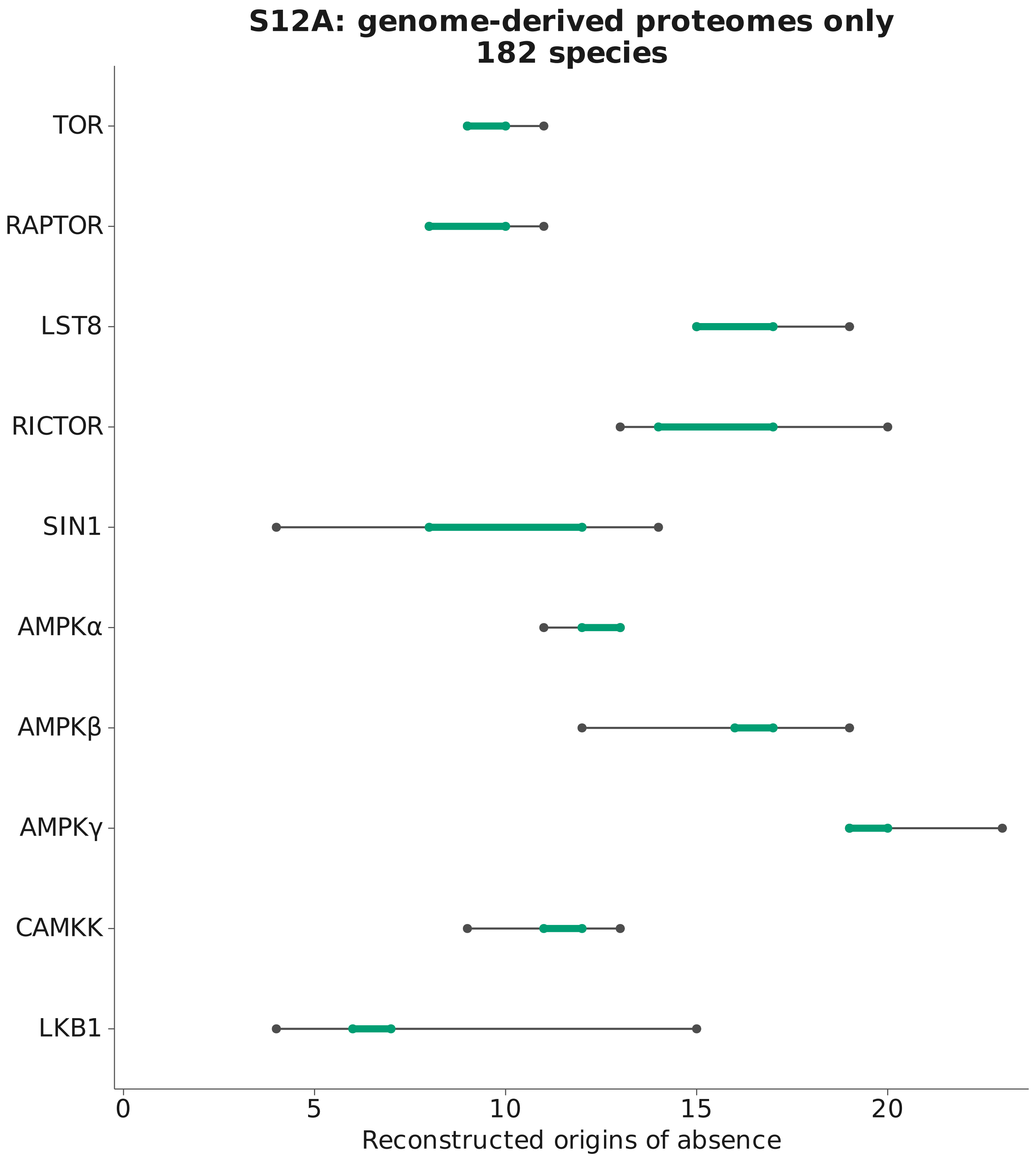


Dark range: all 28 reconstruction conditions
Green range: parsimony conditions only
Ranges describe alternative reconstructions of the same history.

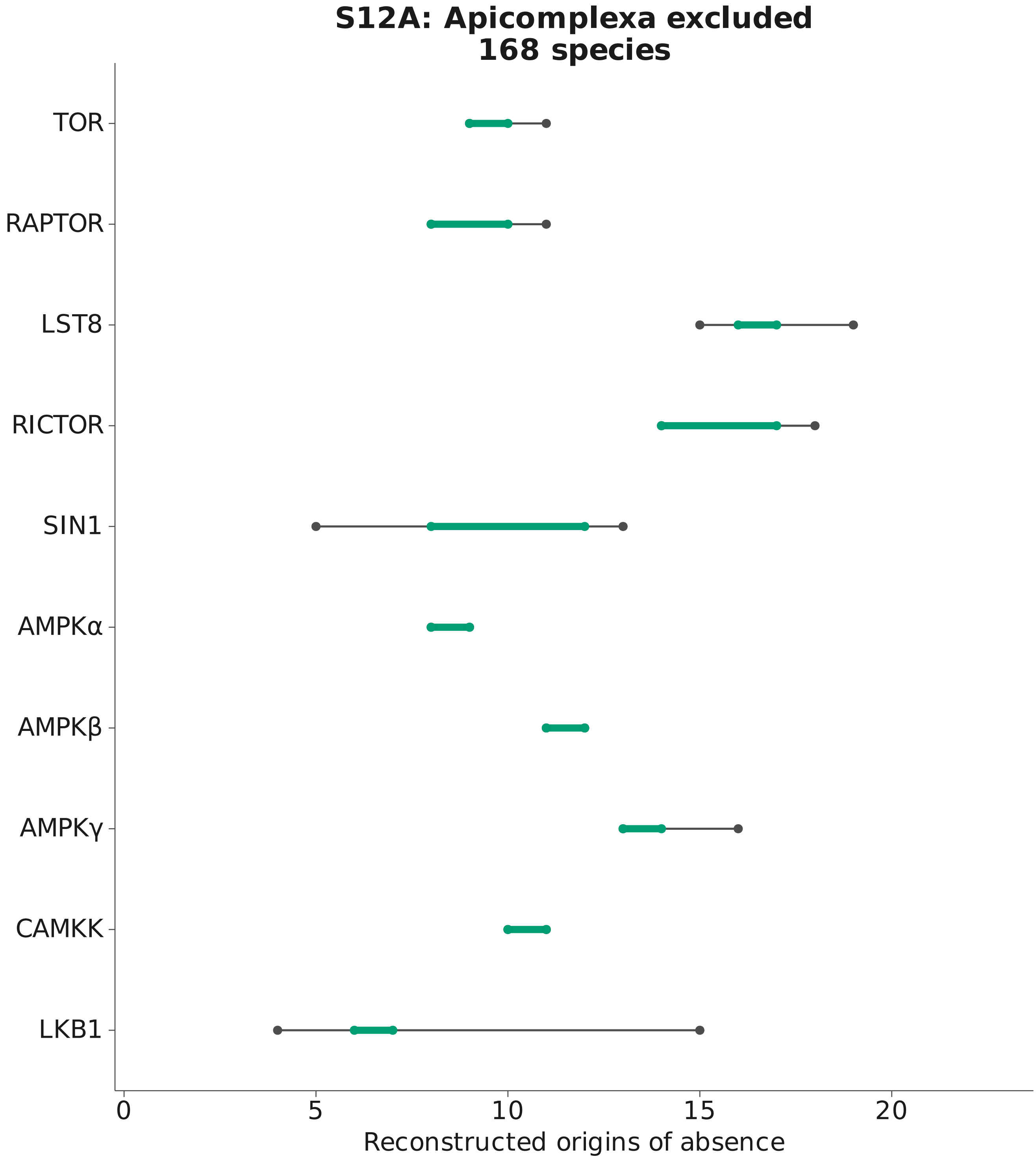

Dark range: all 28 reconstruction conditions
Green range: parsimony conditions only
Ranges describe alternative reconstructions of the same history.

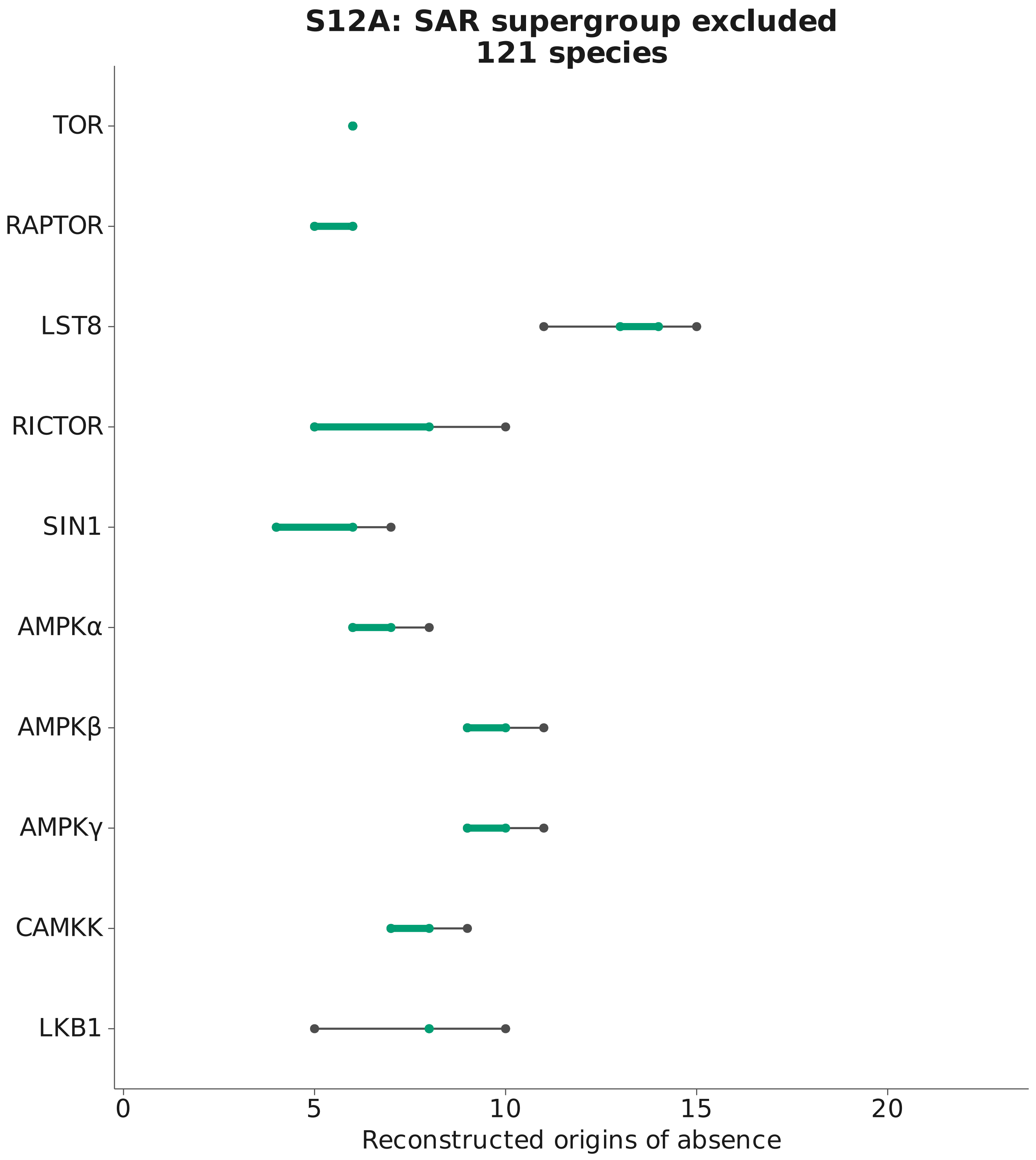


Dark range: all 28 reconstruction conditions
Green range: parsimony conditions only
Ranges describe alternative reconstructions of the same history.

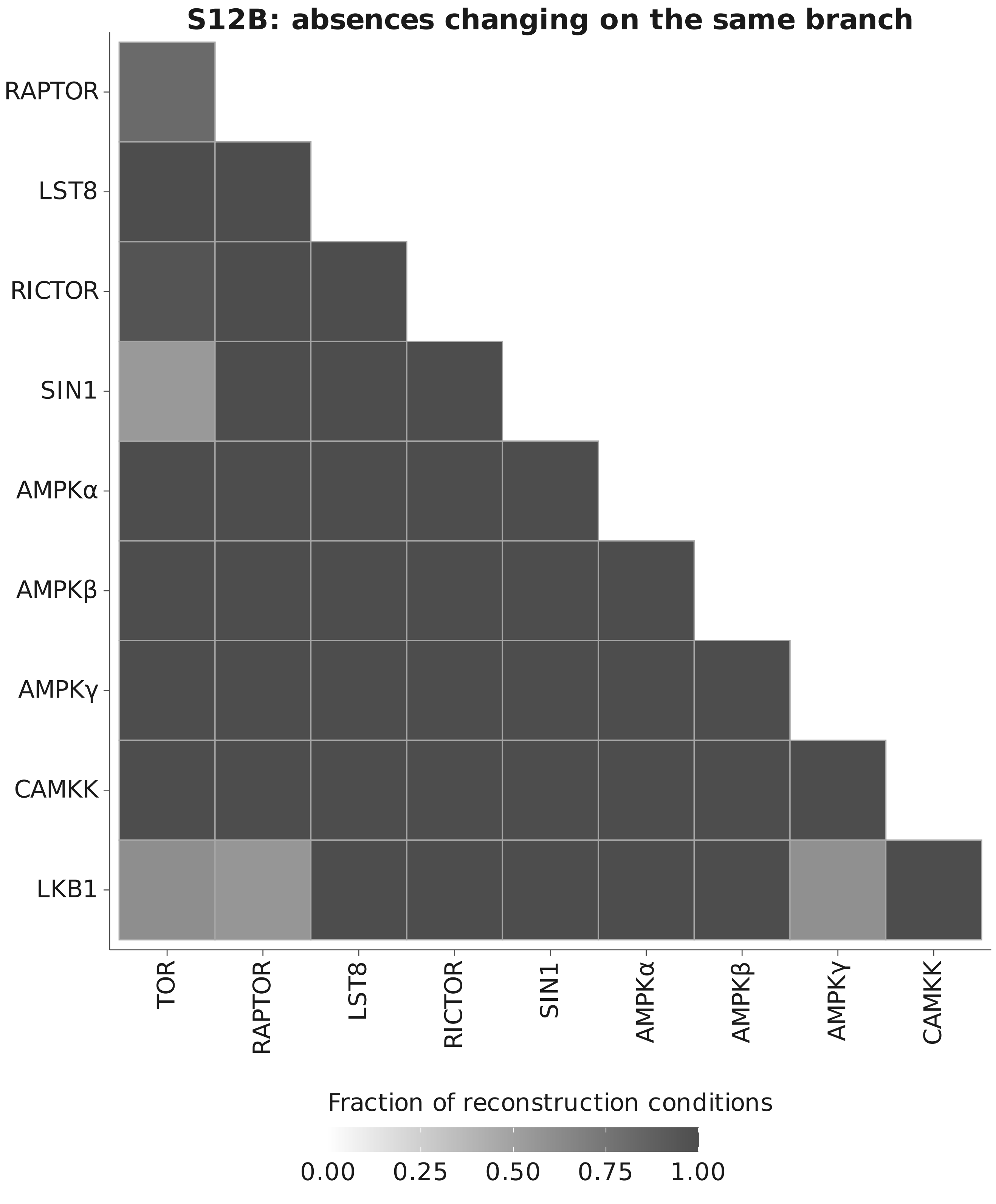


Fraction of 112 reconstruction conditions with a shared branch
Outlined white cells are measured zeros.
Blank positions are self-pairs or the mirrored triangle.
