## Supplementary material for "Divergence of TOR and AMPK regulators across 210 eukaryotes predicts a dispensable TOR in Toxoplasma gondii": S13

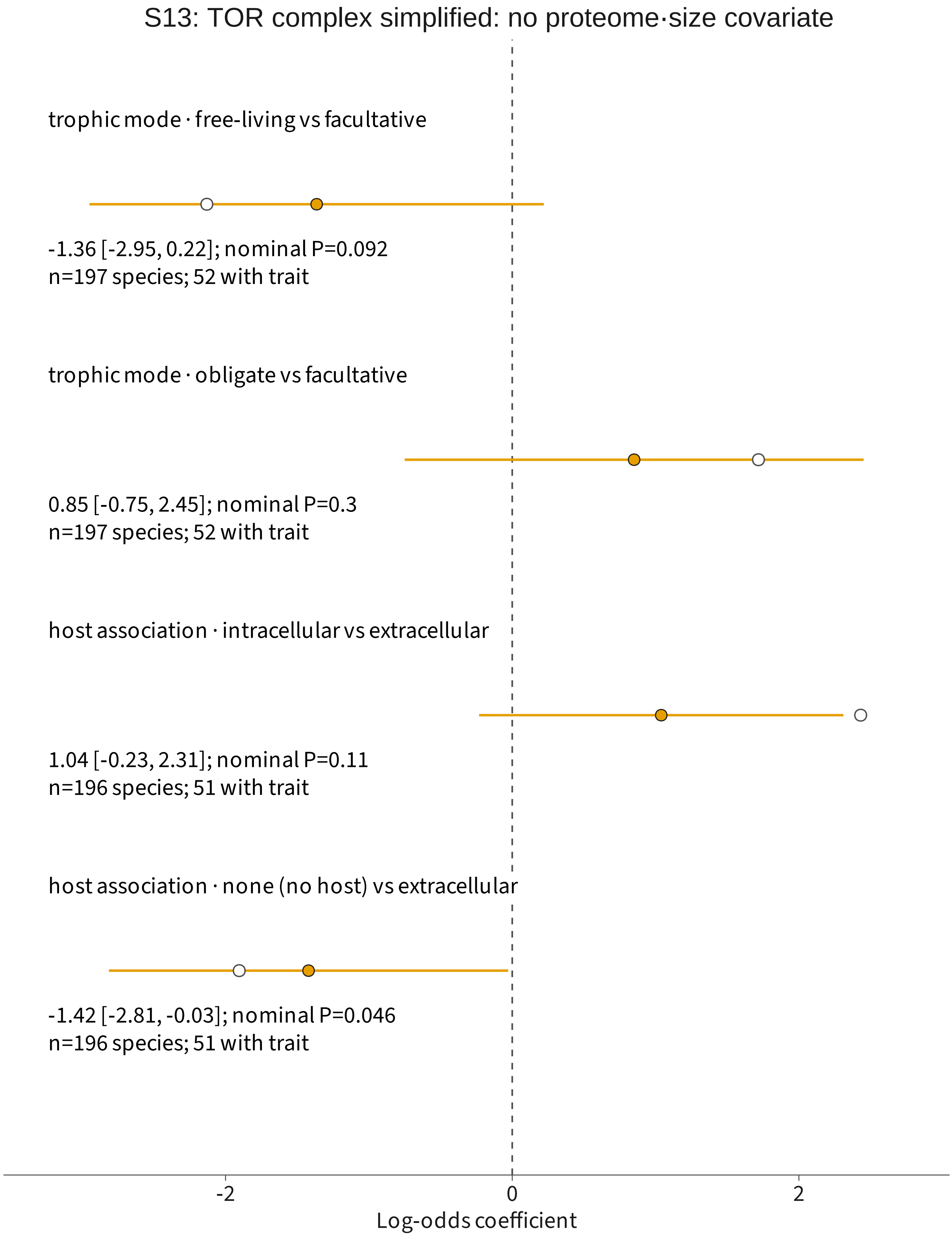


Filled: phylogenetic estimate with 95% Wald interval
Open: no phylogenetic correction; interval unavailable
P values and intervals are unadjusted.

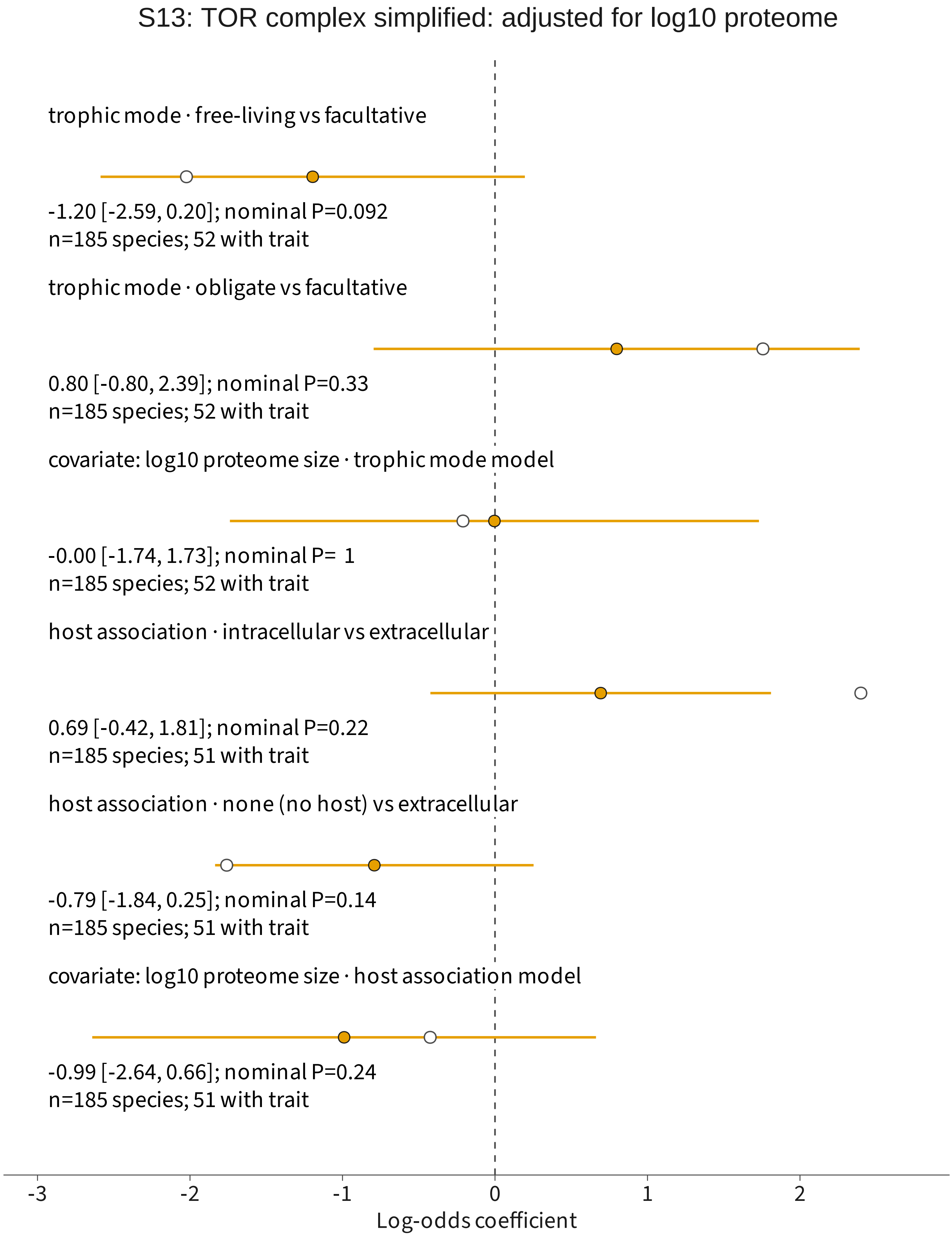


Filled: phylogenetic estimate with 95% Wald interval
Open: no phylogenetic correction; interval unavailable
P values and intervals are unadjusted.

S13: AMPK· subunit absent: no proteome·size covariate

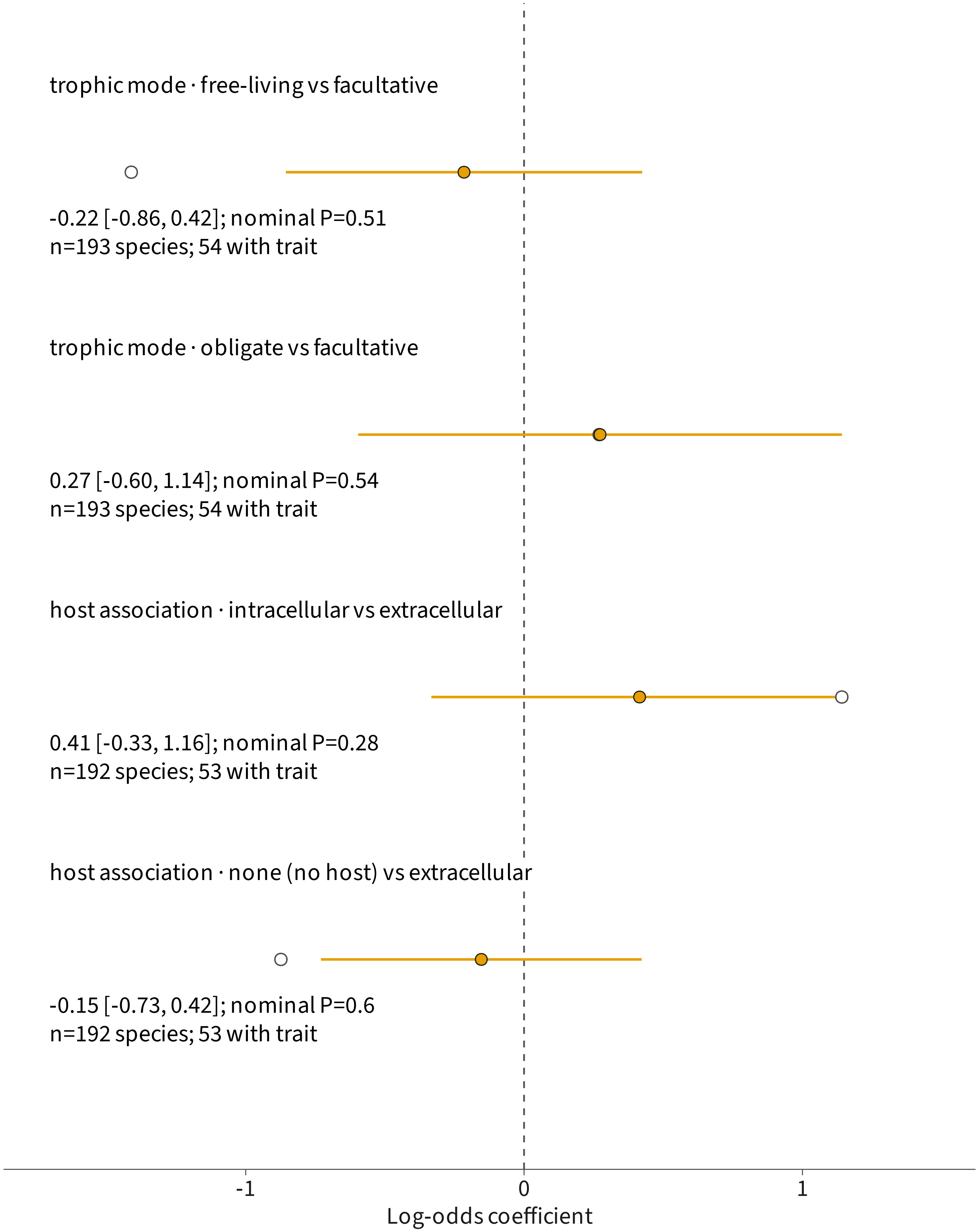


Filled: phylogenetic estimate with 95% Wald interval
Open: no phylogenetic correction; interval unavailable
P values and intervals are unadjusted.

S13: AMPK· subunit absent: adjusted for log10 proteome

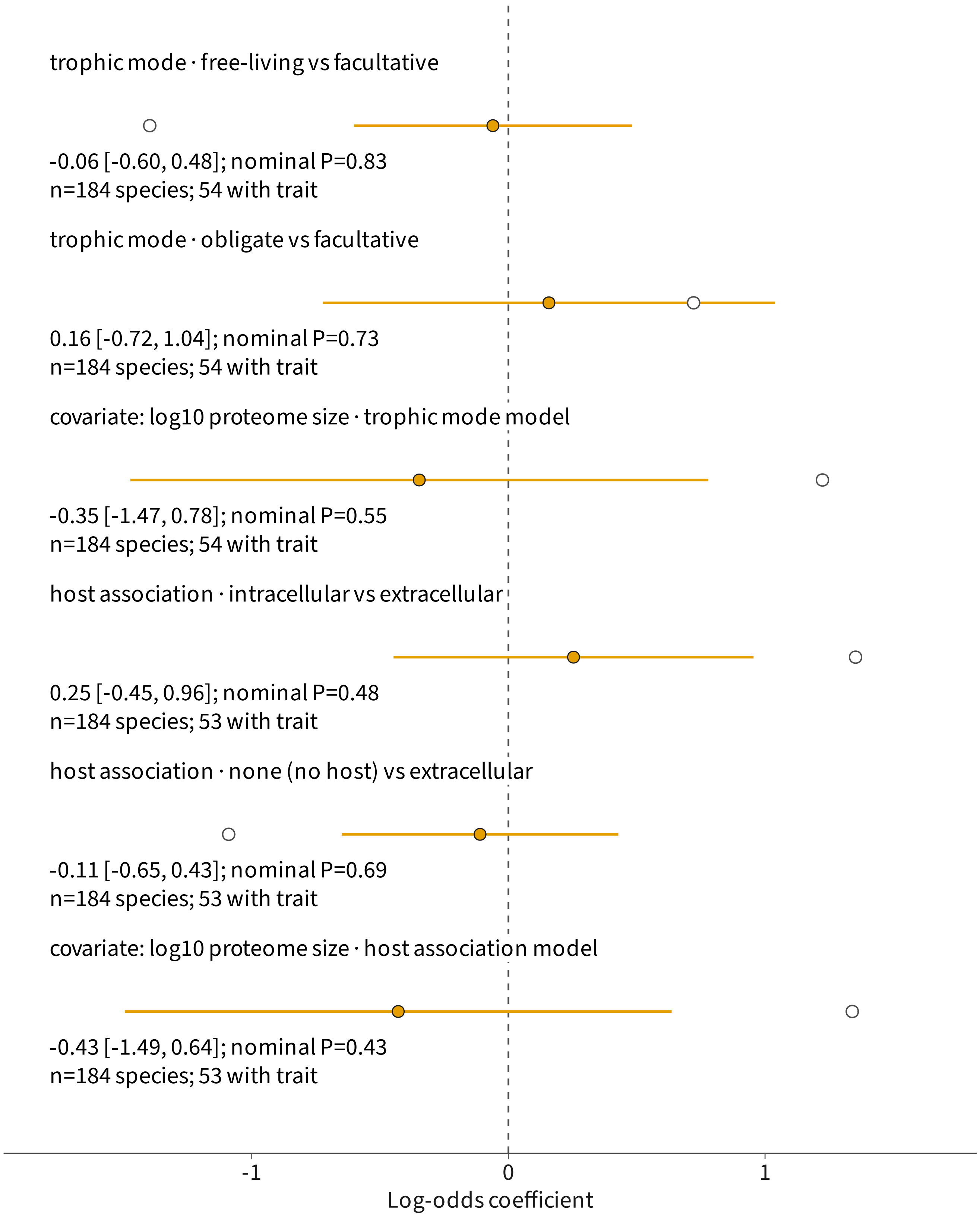


Filled: phylogenetic estimate with 95% Wald interval
Open: no phylogenetic correction; interval unavailable
P values and intervals are unadjusted.

S13: AMPK· nucleotide pocket eroded: no proteome·size

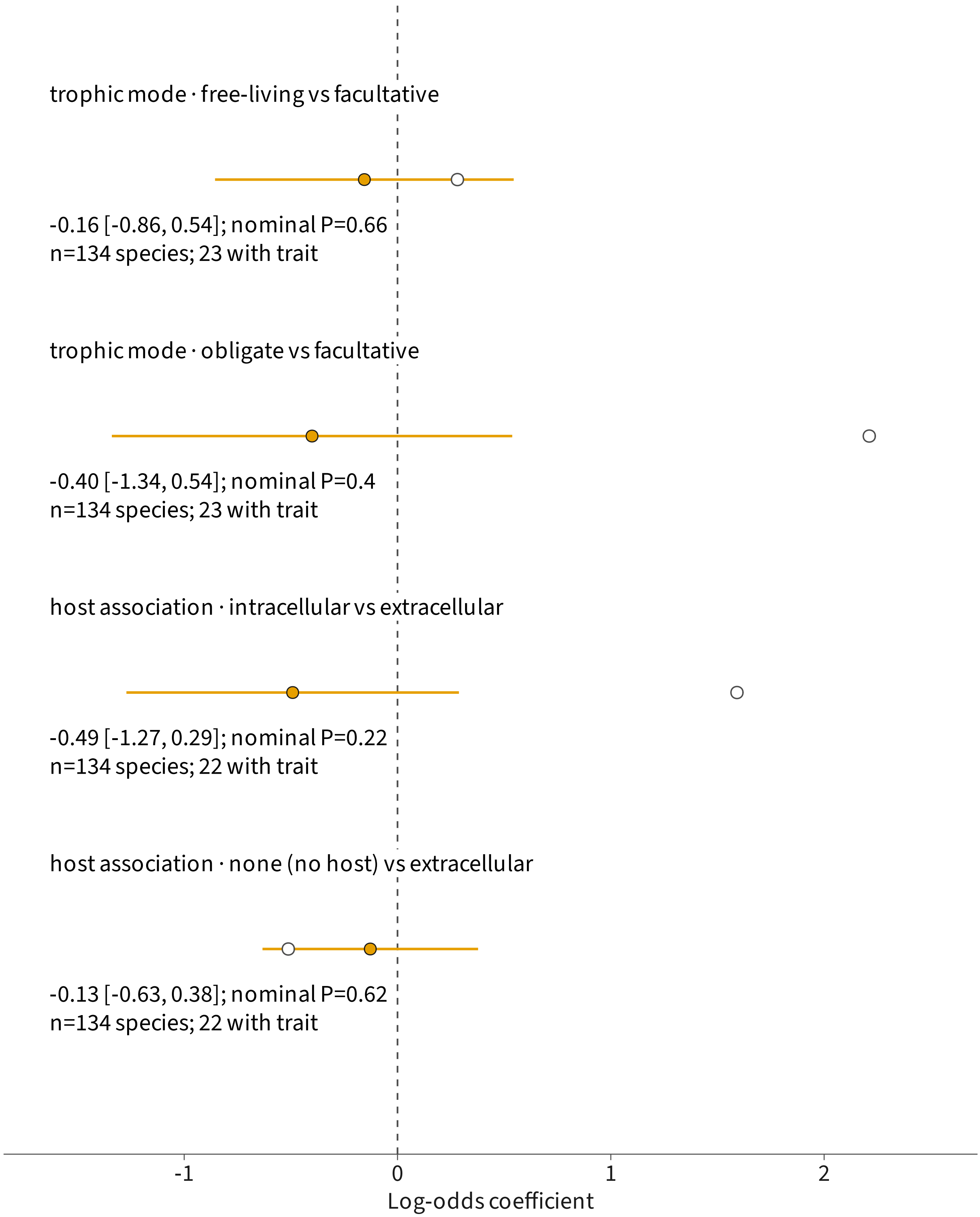


Filled: phylogenetic estimate with 95% Wald interval
Open: no phylogenetic correction; interval unavailable
P values and intervals are unadjusted.

S13: AMPK· nucleotide pocket eroded: adjusted for log10

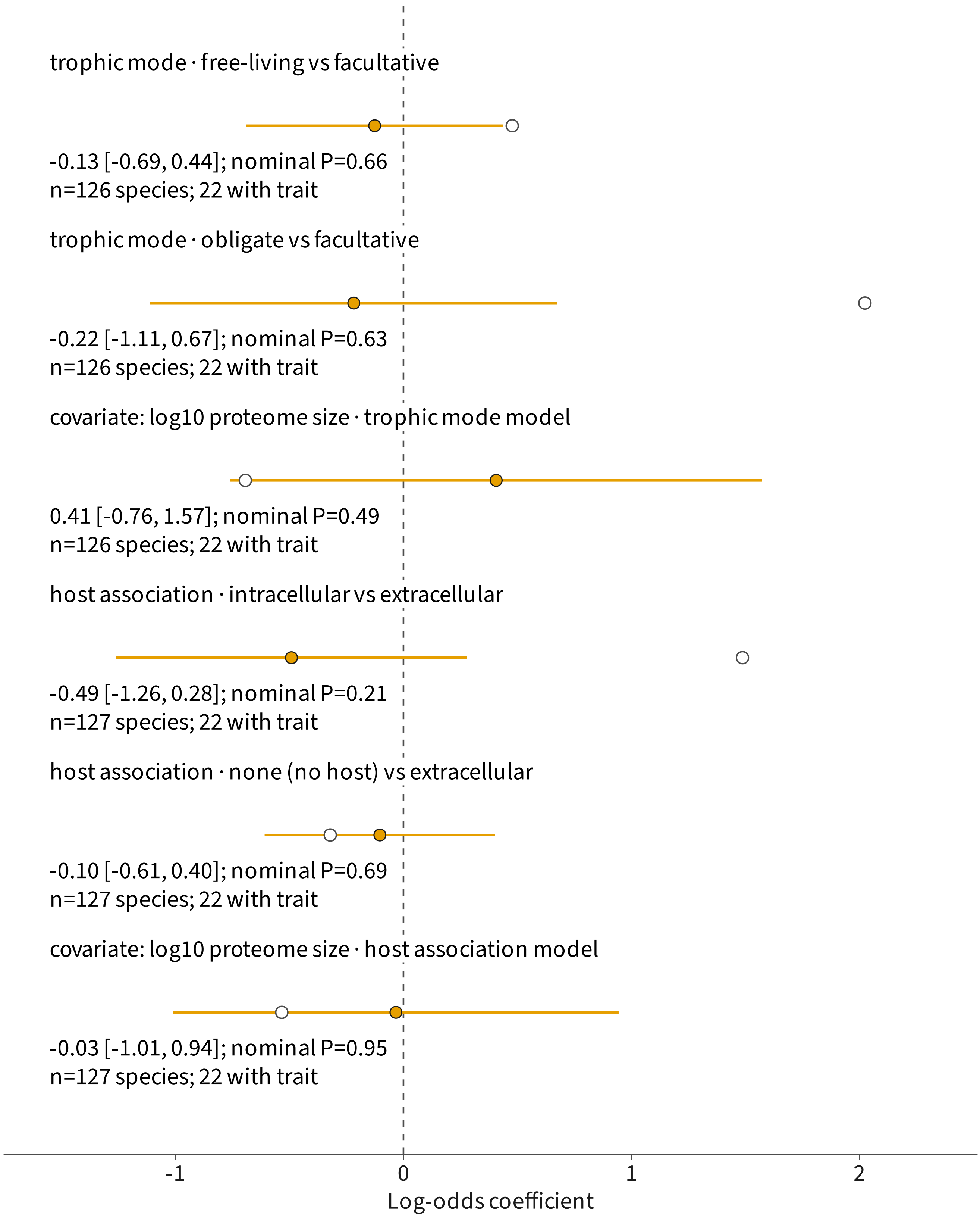


Filled: phylogenetic estimate with 95% Wald interval
Open: no phylogenetic correction; interval unavailable
P values and intervals are unadjusted.
