## Supplementary material for "Divergence of TOR and AMPK regulators across 210 eukaryotes predicts a dispensable TOR in Toxoplasma gondii": S14

### S14: existing multimer geometry and confidence

86 query summaries; 2,150 existing predictions

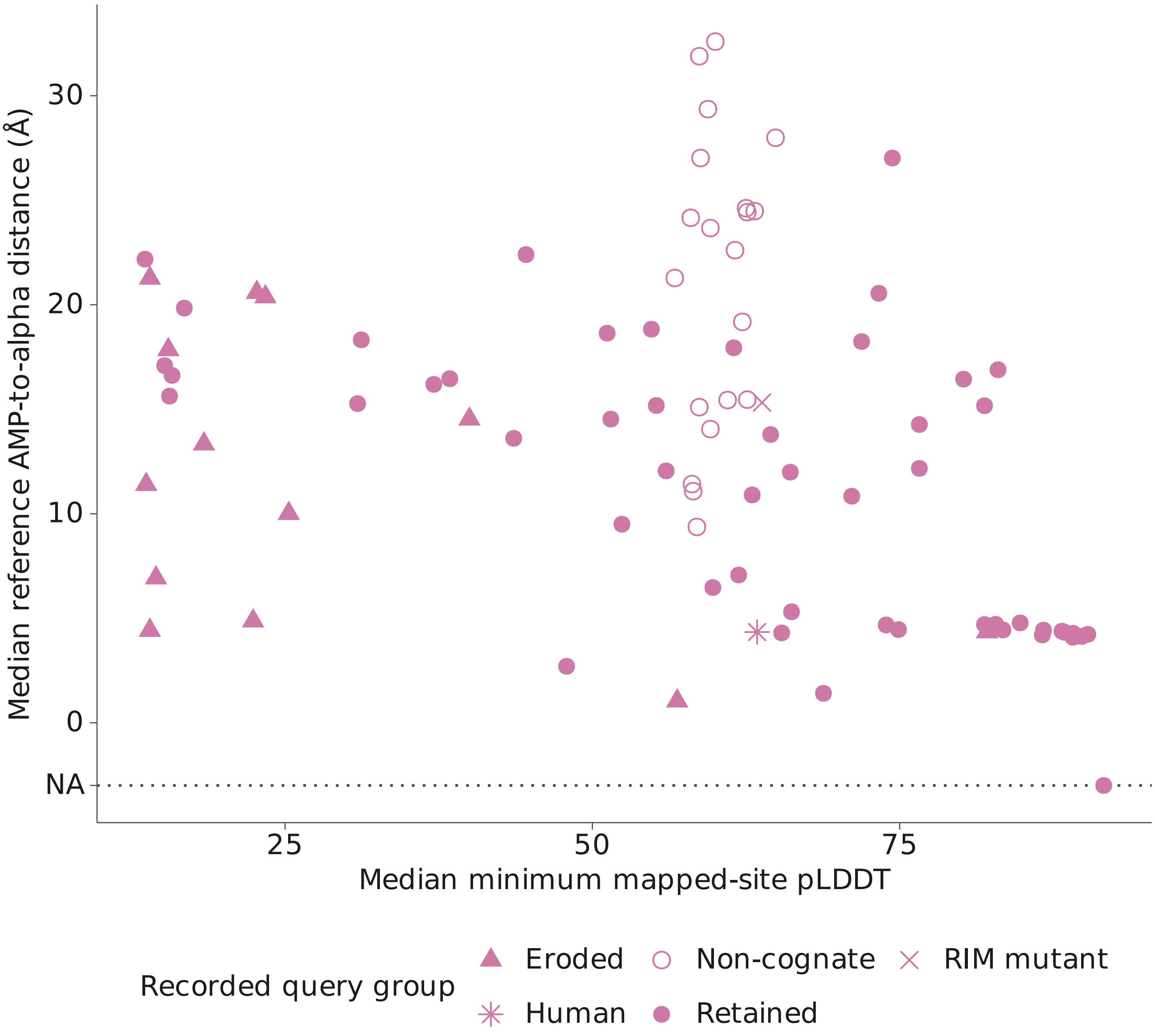


One point per query; 85 have recorded geometry, one is at NA.
Reference AMP is positioned by superposition. The distance
is a predicted geometric measurement, not a binding assay.
