## Supplementary material for "Divergence of TOR and AMPK regulators across 210 eukaryotes predicts a dispensable TOR in Toxoplasma gondii": S15

### A Human partner-context predictions

Reference-positioned AMP; 25 models per construct

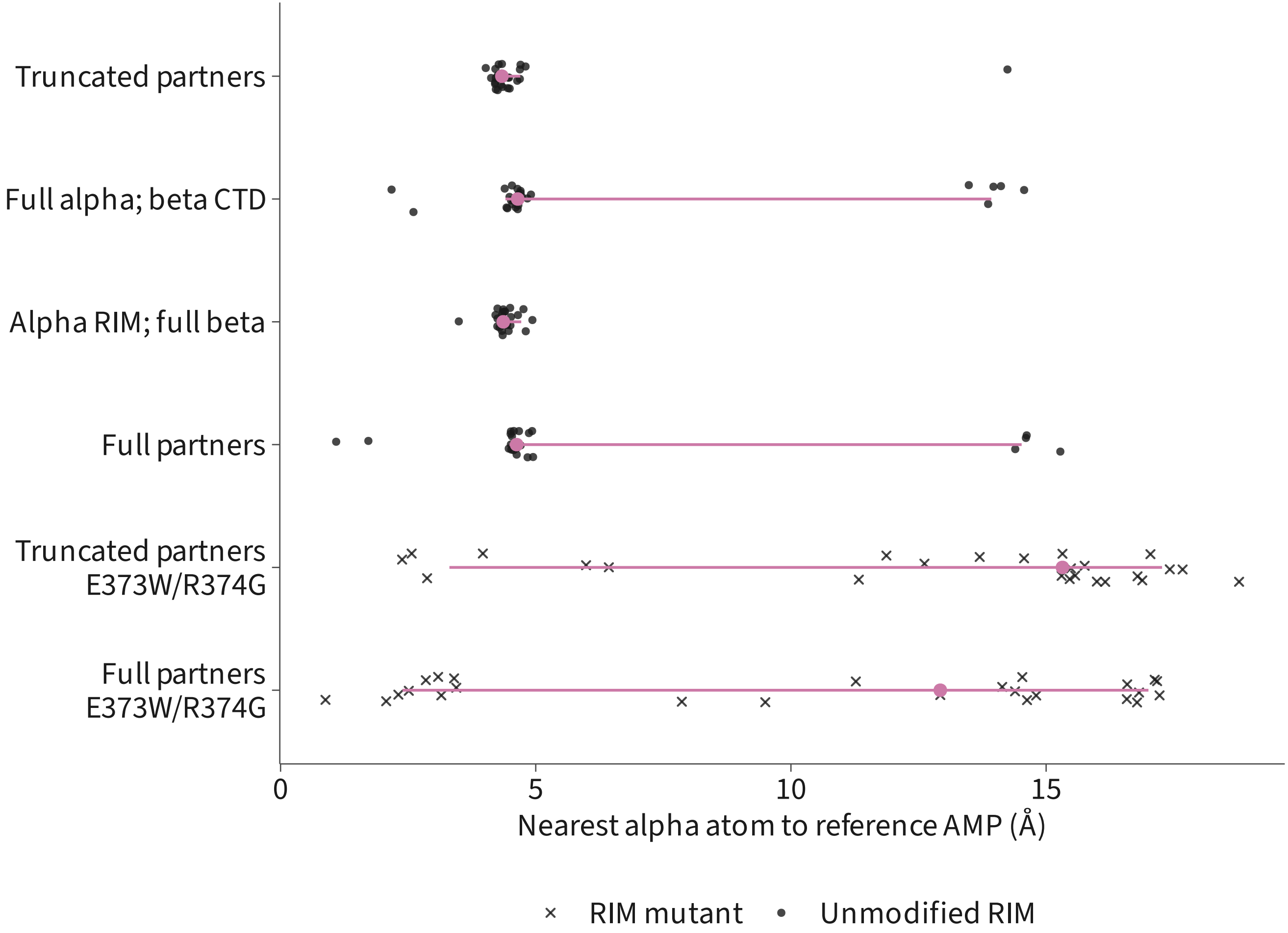


Colored point: median; line: 10th to 90th percentiles.
Dark marks: individual models; all 150 predictions retained.

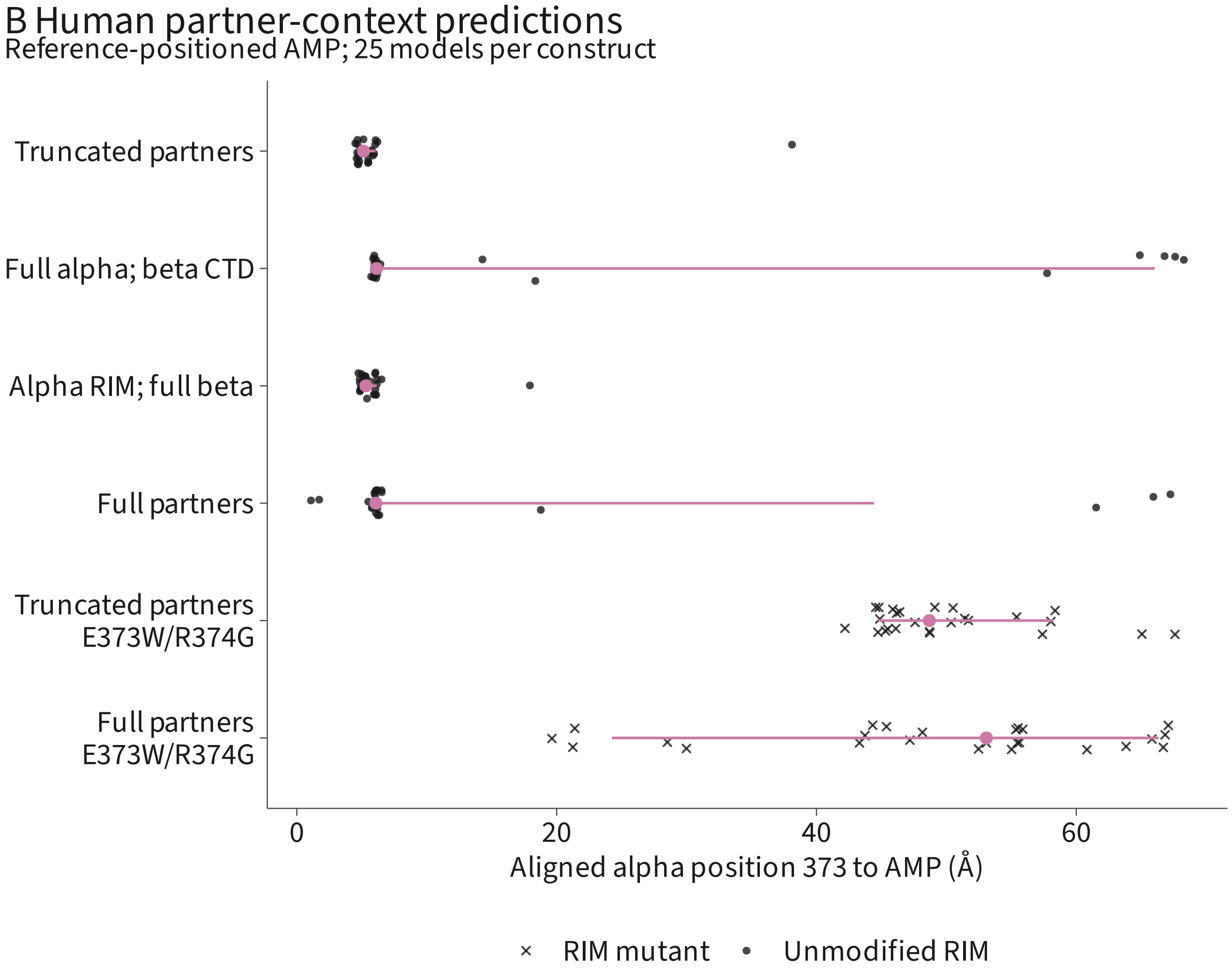


Colored point: median; line: 10th to 90th percentiles.
Dark marks: individual models; all 150 predictions retained.

### C Human partner-context predictions

Reference-positioned AMP; 25 models per construct

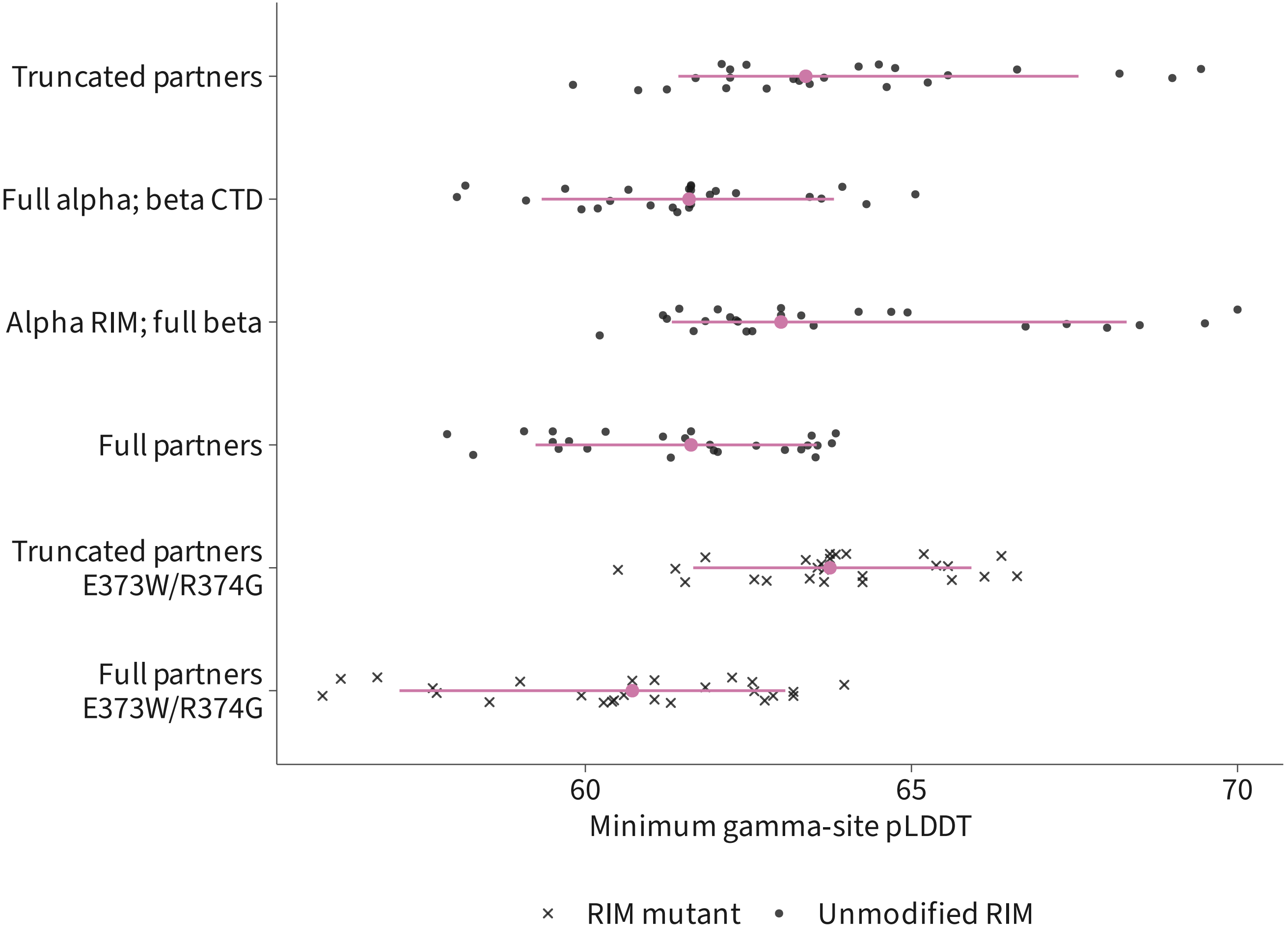


Colored point: median; line: 10th to 90th percentiles.
Dark marks: individual models; all 150 predictions retained.

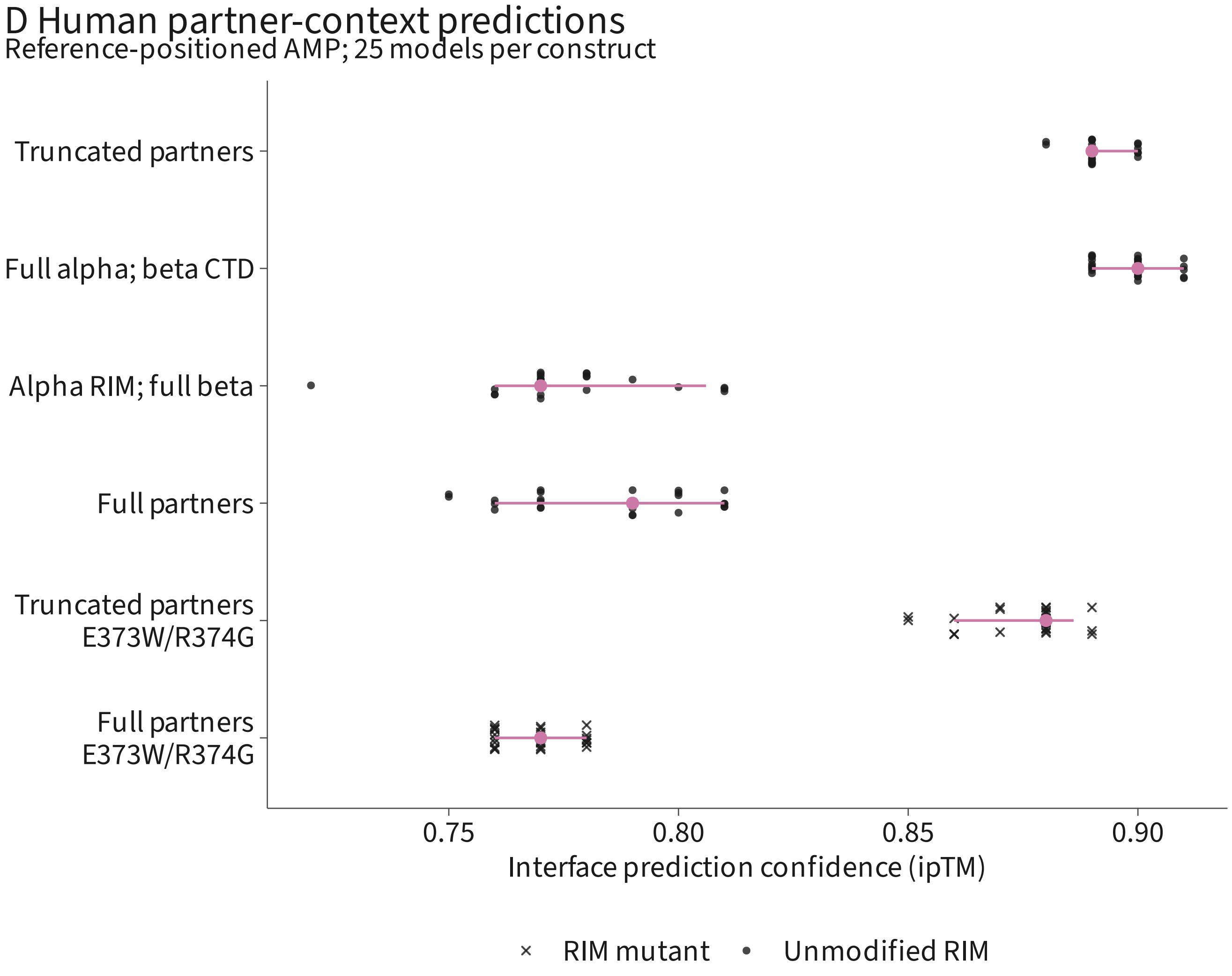


Colored point: median; line: 10th to 90th percentiles.
Dark marks: individual models; all 150 predictions retained.
