## Supplementary material for "Divergence of TOR and AMPK regulators across 210 eukaryotes predicts a dispensable TOR in Toxoplasma gondii": S16

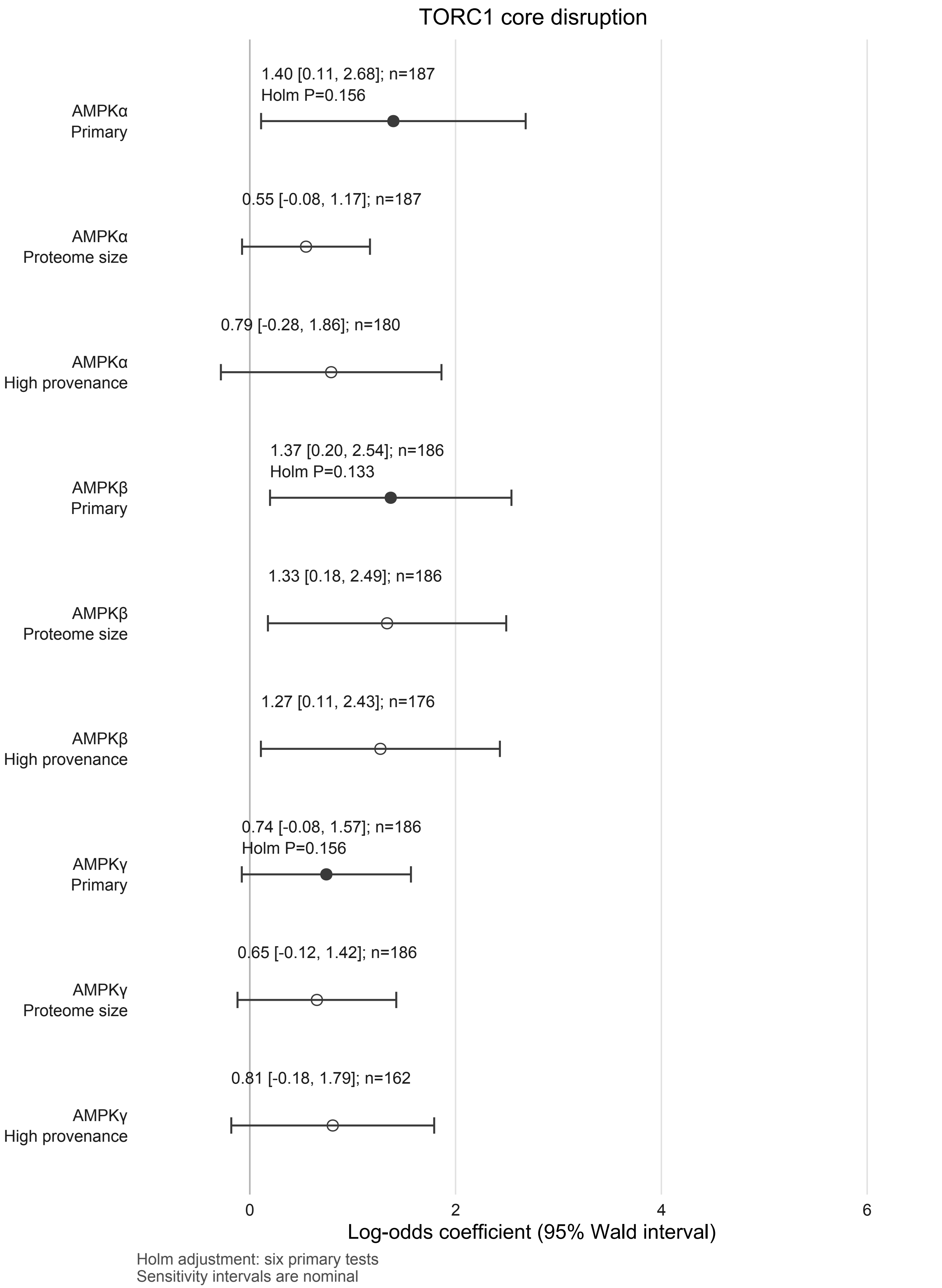
TORC1 core disruption
1.40 [0.11, 2.68]; n=187
Holm P=0.156
AMPKα
Primary
0.55 [-0.08, 1.17]; n=187
AMPKα
Proteome size
0.79 [-0.28, 1.86]; n=180
AMPKα
High provenance
1.37 [0.20, 2.54]; n=186
Holm P=0.133
AMPKβ
Primary
1.33 [0.18, 2.49]; n=186
AMPKβ
Proteome size
1.27 [0.11, 2.43]; n=176
AMPKβ
High provenance
0.74 [-0.08, 1.57]; n=186
Holm P=0.156
AMPKγ
Primary
0.65 [-0.12, 1.42]; n=186
AMPKγ
Proteome size
0.81 [-0.18, 1.79]; n=162
AMPKγ
High provenance
0
2
4
6
Log-odds coefficient (95% Wald interval)
Holm adjustment: six primary tests
Sensitivity intervals are nominal

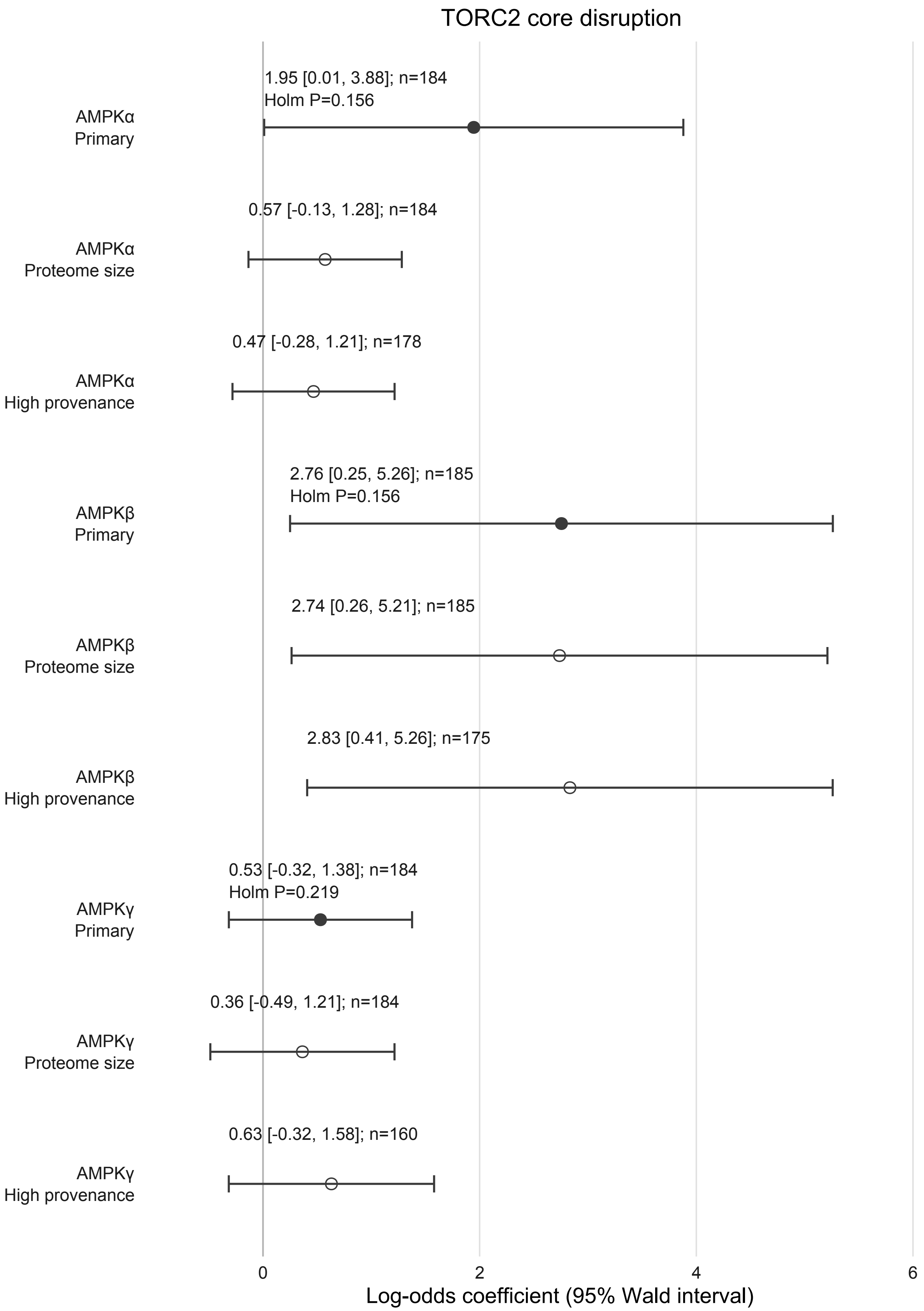

Holm adjustment: six primary tests
Sensitivity intervals are nominal
